# Evolutionary dynamics of temporal transcription factor series in the insect visual brain

**DOI:** 10.64898/2026.01.08.698497

**Authors:** Konstantina Filippopoulou, Elisavet Iliopoulou, Claire Julliot de La Morandière, Christy Lee, Marina Marcet-Houben, Toni Gabaldón, Jingyi Jessica Li, Nikolaos Konstantinides

## Abstract

The nervous system consists of a wide variety of neuronal cell types arranged into complex circuits that support a broad range of behaviors. Patterning of neural stem cells in time through the sequential expression of series of temporal transcription factors is a key contributor to the generation of neuronal diversity. How do temporal series arise and diversify across species to support the evolution of neuronal type? Here, we reconstruct the evolutionary history of the temporal series in the visual brain of insects, spanning 400 million years of evolution; we identify a conserved temporal ground plan, as well as species-specific variations. We find that temporal programs evolve through recurrent modifications of a shared scaffold. Finally, we show that such evolutionary changes in temporal patterning can result in different neuronal type identities. Together, our results reveal both the deep conservation and evolutionary plasticity of temporal patterning programs, and establish temporal transcription factor series as a tractable substrate for the evolution of neuronal diversity.

## Main text

The morphological diversity of the different neuronal types has been appreciated for more than a century. Apart from diverse morphologies, neuronal types differ in other fundamental characteristics, such as connectivity, physiology, and molecular identity. In evolutionary neurobiology, comparative studies examine how these neuronal properties diverge across species, providing insight into how nervous systems adapt to distinct ecological pressures and behavioral demands ^1–7^. All these properties that endow every neuron with a unique identity emerge during development, when the neuronal identity is specified. Therefore, understanding neuronal evolution requires uncovering how developmental programs have changed, as modifications in gene regulation and stem cell lineage progression alter the neuronal output, to give rise to diverse neuronal types.

Research over the last decades has shown that the combination of spatial and temporal patterning of neuronal progenitors, i.e. their position within the tissue and their developmental stage, respectively, is the major mechanism underlying the generation of neuronal diversity in both invertebrates and vertebrates ^8–10^. Temporal patterning is driven by the sequential, temporally-restricted expression of transcription factors (temporal transcription factors - tTFs) in proliferating neural stem cells which alters their ability to produce distinct neuronal types^11,12^.

Although temporal patterning was first functionally described in the vertebrate cortex ^13,14^, the first tTF series was identified in the *Drosophila* ventral nerve cord neuronal progenitors (termed neuroblasts in insects) ^15,16^, the counterpart of the mammalian spinal cord. Since then, multiple tTF series have been discovered across different neuroblast populations in the central nervous system ^11,17–20^. Interestingly, while the principle of temporal patterning is broadly conserved, the specific tTFs that are involved differ markedly across neuronal tissues within the same or between different species ^21^ (**Fig. 1A**). Notably, while the vertebrate spinal cord tTF series shares no homology with any of the fly temporal cascades, it appears to be conserved across vertebrate species and largely shared within the developing vertebrate brain ^12^.

**Fig. 1.**
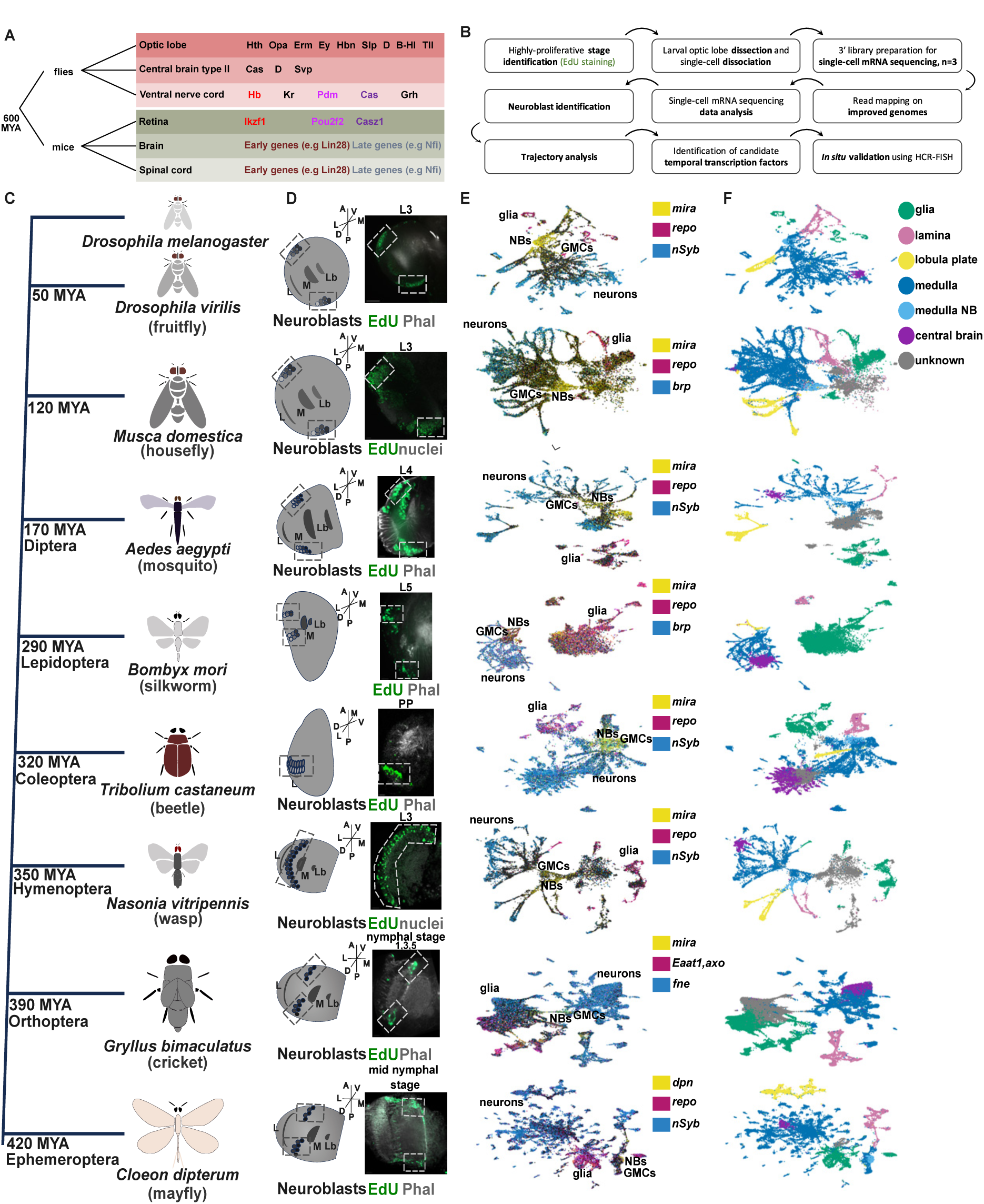
Single-cell profiling of developing insect optic lobes. **(A)** Examples of tTF series from different neuronal structures in flies and mice. Common tTFs between series are color-coded. **(B)** Overview of the experimental workflow, from the identification of highly proliferative developmental stages, through single-cell mRNA sequencing, to the computational identification of candidate tTFs and their *in situ* visualisation. **(C)** Phylogenetic relationships of the insect species analyzed in this study, with divergence times relative to *Drosophila melanogaster* indicated in million years ago (MYA). **(D)** Schematic of the developing insect brains. Dashed boxes highlight the medulla neuroblast region. EdU labeling of highly proliferative developmental stages selected for single-cell mRNA sequencing. Samples are counterstained with phalloidin to label neuropils or DAPI to label nuclei. Dashed boxes indicate proliferating medulla neuroblasts. **(E)** UMAP representations of single-cell mRNA sequencing data from developing optic lobes, showing expression of neuroblast (*mira* or *dpn*; yellow), glial (*repo* or *Eaat1*/*axo*; magenta), and neuronal markers (*nSyb, fne,* or *brp*; blue). **(F)** The same UMAP plots colored by cell type (glia, medulla neuroblasts) or brain region (lamina, medulla, lobula plate, central brain). Abbreviations: M, medulla; Lb, lobula; L, larval stage; PP, prepupa; NBs, neuroblasts; GMCs, ganglion mother cells. Scale bar, 20 µm.

At the same time, several isolated cases of conserved tTF series have emerged. The most compelling example is the expression of three *Drosophila* ventral nerve cord tTF orthologs - Hb/Ikzf1, Cas/Casz1, and Pdm/Pou2f2 - as temporal factors in the developing mouse retinal progenitor cells ^22–24^ (**Fig. 1A**). Similarly, individual *Drosophila* tTFs are deployed temporally in various vertebrate neural stem cell populations even when the remainder of the cascade is not conserved ^11,25^. These scattered but intriguing parallels suggest that some components of temporal patterning may be evolutionarily ancient, while others might have diversified or been replaced. Thus, understanding the evolution of neuronal development requires examining how temporal series in neural stem cells arise and diversify across species.

This question cannot be addressed by comparing animals as evolutionarily distant as flies and mice that diverged ∼600 million years ago (**Fig. 1A**). Moreover, to study the evolution of temporal series, one needs a neuronal tissue model that is comprehensively described, cell-type resolved, and comparatively accessible, allowing the identification of both conserved and divergent mechanisms. In the insect brain, the optic lobes - the part of the brain located behind the retina processing visual information - consist of 3 main neuropils (lamina, medulla, and lobula complex), which are conserved across insects both structurally and functionally, providing exactly such a model ^26^. More specifically, the medulla region of the optic lobes enables meaningful comparative analyses, as it is the richest in neuronal diversity, experimentally tractable and its temporal patterning program has been dissected in exceptional detail by our group and others ^11,17^: in the developing *Drosophila* medulla, a series of 14 tTFs (*hth, dmrt99B, erm, opa, L, ey, hbn, scro, slp1, slp2, D, BarHI, BarH2,* and *tll*) has been identified, forming the longest and best-resolved tTF cascade described in any neural stem cell population, contributing to the generation of over a hundred distinct neuronal types ^27–30^. The progression of this series is largely governed by sequential activation of successive tTFs and repression of the preceding factor; however, this core cascade operates within a broader gene regulatory network that includes branching regulatory inputs, collectively coupling stem cell age to neuronal identity.

Here, we present a systematic study of how this fundamental neurodevelopmental program evolves across the insect phylogeny spanning ∼400 million years. We performed single-cell mRNA sequencing of the developing optic lobes of eight different species, reconstructed the neuroblast trajectories to identify candidate tTFs and cross-validated their expression by HCR-FISH (**Fig. 1B**). By tracing the evolutionary history of optic lobe temporal series, we uncover a conserved core of tTFs that was present in the last common ancestor of hemimetabolous and holometabolous insects, alongside divergent factors that likely allowed insects to adapt their neuronal repertoire to defined ecological pressures. We further identify evolutionary mechanisms shaping temporal series divergence and infer their impact on lineage-specific neuronal identities. Together, these results provide the first unified view of how temporal patterning programs evolve, revealing the rules by which neural stem cells diversify neuronal types over hundreds of millions of years.

### The developing optic lobe in single-cell resolution across insect phylogeny

We first selected representatives from different insect orders that diverged from *D. melanogaster* between 50 and 420 million years ago (mya), spanning the insect phylogenetic tree and with high-quality genomes available: the fruit fly *Drosophila virilis* (Diptera, 50 mya), the house fly *Musca domestica* (Diptera, 120 mya), the mosquito *Aedes aegypti* (Diptera, 170 mya), the silkworm *Bombyx mori* (Lepidoptera, 290 mya), the beetle *Tribolium castaneum* (Coleoptera, 320 mya), the wasp *Nasonia vitripennis* (Hymenoptera, 350 mya), the cricket *Gryllus bimaculatus* (Orthoptera, 390 mya), and the mayfly *Cloeon dipterum* (Ephemeroptera, 420 mya) (^31^ (**Fig. 1C**). Second, we identified the developmental stages at which neurogenesis occurs in the optic lobes of each species. Given the substantial differences in life cycles (**Fig. S1**), we focused on stages showing extensive proliferation in the developing optic lobes as previously described morphologically ^32–36^ (**Fig. 1D**) and performed EdU pulse-chase experiments (**Fig. 1D** and **Fig. S2A–D**), which revealed the following proliferative stages: *D. virilis* larval stage 3 (L3), *Musca* L3, *Aedes* L4, *Bombyx* L5, *Tribolium* prepupa, and *Nasonia* L3. For hemimetabolous insects lacking true larval stages, we assayed nymphal stages and selected *Gryllus* nymphal stages 1, 3, and 5 ^37^, and *Cloeon* mid-nymphal stage, as periods of high optic lobe neuroblast proliferation (**Fig. S2E–F** and **Methods**). Finally, we developed tissue dissociation protocols for the optic lobes of each species to obtain single cells for downstream applications (**Methods**).

We performed single-cell 3’ mRNA sequencing using the 10X Genomics platform. To improve genome annotations, we combined reads from our single-cell data with additional reads from adult single-cell RNA-seq datasets from the same species and applied GeneExt to extend the 3’ end of the mRNAs ^38^ . It has been observed that 3′ ends are often misannotated for neuronal mRNAs, as transcripts in the nervous system tend to have longer 3′ UTRs ^39–41^ (**Methods** and **Fig. S3**). After read mapping, initial quality control, and preprocessing, we recovered a total of 304,485 developing optic lobe cells across all species (**Methods** and **Fig. S4**).

In the insect developing optic lobes, a wave of neurogenesis converts the neuroepithelial cells into neuroblasts. Neuroblasts divide asymmetrically to self-renew and produce intermediate precursors, called ganglion mother cells (GMCs). GMCs then divide once to generate two different neurons or glial cells ^42^. To identify the different progenitors and differentiated cell types, we examined the expression of conserved markers: *shotgun* (*shg*) for neuroepithelial structures and neuroblasts ^43^, *miranda* (*mira*), *deadpan* (*dpn*) and *worniu* (*wor*) for neuroblasts ^44,45^, *asense* (*ase*) for neuroblasts and GMCs ^46^, *found in neurons (fne)*, *embryonic lethal abnormal vision* (*elav*) ^47^, *neuronal Synaptobrevin* (*nSyb*), and *bruchpilot* (*brp*) ^48^ for neurons, and *reversed polarity* (*repo*), *Excitatory amino acid transporter 1* (*Eaat1*), and *axotactin* (*axo*) for glia ^49^ (**Fig. 1E** and **Fig. S5**). Because neuroblasts are generated progressively during the progression of the neurogenic wave, neuroblasts of every age are recovered at any given time point. As observed in the equivalent *D. melanogaster* dataset ^11^, the two-dimensional Uniform Manifold Approximation and Projection (UMAP) representations recapitulated the progression from progenitors to differentiated cells: neuroblasts (yellow) occupy central positions in the UMAP plots, leading into neuronal trajectories (blue), while most glia form relatively isolated clusters (red). The only exception is in hemimetabolous insects (*Gryllus* and *Cloeon*), where some differentiated neurons already form separate, distant clusters from neuroblasts; these isolated neurons likely represent embryonically-born neurons that have differentiated and become active in hatchlings, which have a functional visual system.

Finally, because our dissections and dissociations did not allow us to separate the individual optic lobe neuropils, we relied on conserved molecular markers to identify cells from these neuropils in our datasets. Both the lamina and medulla originate from the Outer Proliferation Center (OPC), which generates medulla neuroblasts medially and lamina precursor cells laterally. To identify lamina neurons, we searched for conserved markers of this lineage - *dachsund* (*dac*), *eyes absent* (*eya*), *single-minded* (*sim*), *glial cells missing* (*gcm*), and *tailless* (*tll*) ^11,50,51^ (**Fig. S6**). Most of these genes are reliable lamina markers in all insects, the most conserved ones being *eya*, *sim*, and *dac*. Neurons of the lobula plate (T4/T5) as well as T and C neurons (T2/T3 and C2/C3) originate from the Inner Proliferation Center (IPC), which is defined by the expression of *tll*, *dac*, and *acj6* ^52,53^ (**Fig. S6**). Using these genes, we were able to define lobula plate cell types in all insects, except for *Gryllus* (**Fig. 1F**). Having identified the major neuropils, we next investigated the neural stem cell populations present in our datasets. We examined the expression of conserved *D. melanogaster* neuroblast markers such as *mira* (**Fig. 1E**), *dpn*, and *wor* (**Fig. S5**). To identify medulla neuroblasts and distinguish them from differentiating cells, we used the co-expression of *mira* and *dpn*, a well-established hallmark of medulla neuroblasts; the only exception was *Cloeon*, for which selection was based solely on *dpn* expression (**Methods**). As in *D. melanogaster*, medulla neuroblasts were positioned between the medulla and lamina neurons (**Fig. 1F**), consistent with the *in vivo* organization of the OPC.

In summary, we generated developmental atlases of eight insect optic lobes during major proliferative stages, annotated their constituent neuropils, and identified the medulla neuroblast populations.

### A conserved core of the temporal series

#### *In silico* identification of temporal transcription factors

To compare the temporal transcription factor (tTF) series across insects (**Fig. 2A**), we first identified orthologous genes. To do so, we reconstructed phylomes ^54^ - comprehensive collections of gene-specific phylogenetic trees - for all genes of each species. This allowed us to classify one-to-one, one-to-many, many-to-one, and many-to-many homologs, as well as species-specific genes (**Fig. S7, Methods,** and **Table S1**). Using these orthology relationships, we selected representative *D. melanogaster* tTFs for early (*hth*), mid (*opa* and *ey*), and late (*scro* and *BarHI*) temporal windows and examined their expression patterns in the neuroblast

**Fig. 2.**
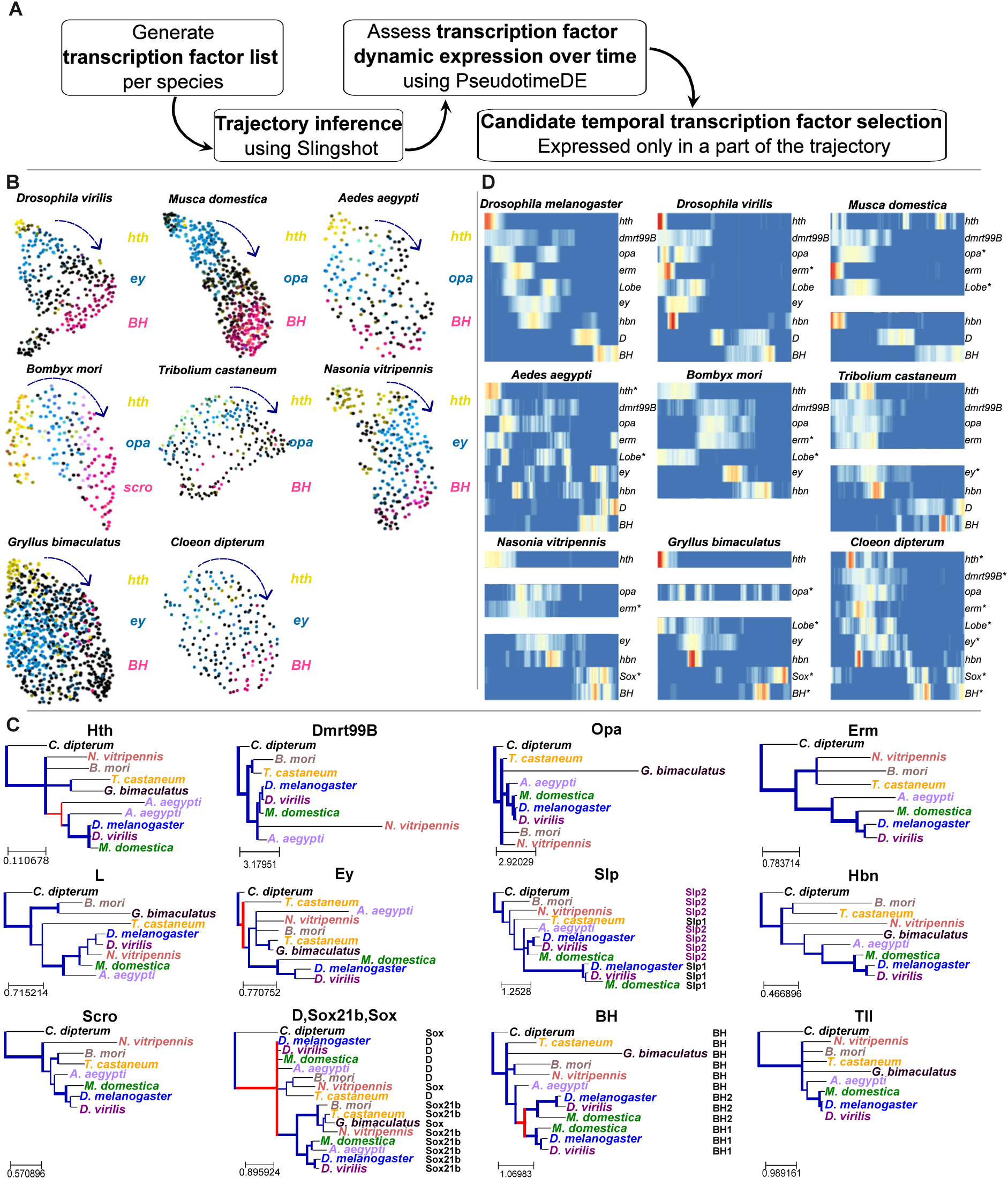
Trajectory analysis of developing neuroblasts reveals a conserved core of the temporal series. **(A)** Schematic overview of the computational pipeline used for trajectory inference and unbiased identification of candidate temporal transcription factors. **(B)** UMAP projections of neuroblasts from the different insect species, showing the expression of conserved temporal transcription factors corresponding to early (*hth*; yellow), mid (*ey* or *opa*; blue), and late (*BarH* or *scro*; magenta) temporal windows. Arrows indicate trajectory. **(C)** Phylogenetic trees for *Drosophila melanogaster* tTFs and their homolog counterpart expressed temporally in other species. Nodes are collapsed when bootstrap values are below 50, the branches are thicker when the bootstrap is high and are coloured in blue for speciation events and in red for duplication events. Scale bars represent the estimated number of substitutions per site along each branch. **D)** Heatmap showing the *Drosophila melanogaster* tTF homologs that are temporally expressed across all analyzed insect species, highlighting the conserved core of the temporal series. The asterisk (*) refers to non 1:1 ortholog genes. Gaps in the heatmaps illustrate the lack of temporal expression, or the absence of the gene from the genome of the respective species. Non-temporally expressed TFs are shown in **Fig. S9**.

UMAP plots across species (**Fig. 2B**). Their expression pattern revealed that (a) neuroblasts from all insects undergo temporal patterning at the developmental stages we profiled, (b) as in *D. melanogaster*, neuroblasts display time-restricted (temporal) expression of potential tTFs on the UMAP space, and (c) the temporal series appears conserved to a substantial extent across insects.

To investigate temporal regulation in these insects and identify medulla neuroblast tTFs in an unbiased manner, we first computationally isolated the medulla neuroblast clusters and performed trajectory inference using Slingshot ^55^. We then sought to identify all transcription factors with dynamic expression along this trajectory: first, we generated transcription factor lists for each insect species (**Table S2 and Methods**). Next, we developed a computational pipeline to identify TFs dynamically expressed along the inferred trajectory (**Fig. S8 and Methods**). Using PseudotimeDE ^56^, we applied three filters to define dynamically expressed transcription factors: (a) an expression threshold ensuring the gene is expressed along the trajectory; (b) a requirement that expression approaches zero at some point, ensuring temporality; and (c) a PseudotimeDE p-value to account for uncertainty in pseudotime estimation. Finally, we took into consideration their relative expression level in their temporal window, which is high in validated temporal transcription factors, as previously demonstrated for *D. melanogaster* ^11^. We then applied this standardized pipeline (**Fig. 2A**) to the medulla neuroblasts of all insects, including *D. melanogaster*, and identified tTF candidates in each species (**Fig. S9** and **Table S3**). In total, we found 14 candidate tTFs in *D. melanogaster* (similar to what was identified before ^11^), 15 in *D. virilis*, 15 in *Musca*, 18 in *Aedes*, 17 in *Bombyx*, 16 in *Tribolium*, 15 in *Nasonia*, 8 in *Gryllus*, and 14 in *Cloeon*. Strikingly, despite differences in developmental tempo, eye size, and ecological reliance on vision, the number of temporal transcription factors is broadly similar across species; the reduced number observed in *Gryllus* may reflect biological divergence or technical factors, including developmental stage, genome annotation, or expression sensitivity. This suggests that the consistent number of tTFs reflects a conserved developmental strategy for generating neuronal diversity.

It was immediately apparent that most *D. melanogaster* tTFs were present in the majority of species examined. To further investigate and validate the tTF evolutionary relationships, we reconstructed phylogenetic trees for all known *D. melanogaster* tTFs and their homologs across the other species (**Fig. 2C**). Overall, neuroblasts across species expressed a largely conserved core set of tTFs, with between 9 and 14 *D. melanogaster* tTFs detected depending on the species (with the exception of *Gryllus* that expressed 7 of them) (**Fig. 2C, Fig. S9**). We then examined whether the *D. melanogaster* tTFs that are missing were in fact absent from the corresponding genomes. We found that (a) the two *slp* paralogs are present only in the two drosophilid species; (b) *BarH2* arose in Diptera after the divergence of *Aedes* and *Musca*; (c) *dmrt99B*, *erm*, *scro*, and *slp* were absent from the *Gryllus* genome; and (d) strikingly, *ey* was not annotated in the *Musca* genome. To verify whether *ey* was truly absent in *Musca*, we used available bulk head transcriptomic data ^57^ to assemble a transcriptome (**Fig. S10Α** and **Methods**) and performed BLAST searches against it. We identified an *ey* ortholog and confirmed its identity based on a phylogenetic tree of Ey proteins (**Fig. S10B**) and incorporated the locus into the existing genome annotation. We confirmed by HCR-FISH that the identified *ey* gene is expressed in the eye field of the *Musca* eye-antennal disc (**Fig. S10C**). We then examined its expression in the developing optic lobe and confirmed that it is not expressed in the neuroblasts, while low neuronal expression can be detected in our *Musca* single-cell mRNA-sequencing atlas (**Fig. S10D**), as well as by HCR-FISH (**Fig. S10E**). This is also consistent with differential *ey* enhancer usage in neuroblasts and neurons in *D. melanogaster* ^30^, suggesting that only the neuronal enhancer is active in *Musca*.

These comparisons reveal a conserved “core” of tTFs (**Fig. 2D**) that can be traced back to the last common ancestor of almost all insects. In particular, 8 of the 14 *D. melanogaster* tTFs (*hth, dmrt99B, erm, opa, L, ey, hbn, D,* and *BarH*) were likely expressed in the common ancestor of hemi- and holometabolous insects and, hence, were deployed temporally for more than 400 million years. Furthermore, 11 of these factors were already expressed temporally in the last common ancestor of holometabolous insects, which lacked a second paralog of the *slp* and *BarH* genes.

### *In situ* cross-validation of temporal transcription factors

Although our previous work in *D. melanogaster* established trajectory inference as a reliable strategy for identifying temporal transcription factors, we cross-validated conserved and divergent candidates to verify their temporal expression pattern, including genes with noisy expression profiles, genes not expressed in medulla neuroblasts and tTF negative controls that are absent in *D. melanogaster* but expressed in other species. To this end, we performed HCR-FISH in the developing optic lobes of 7 species (*D. virilis, M. domestica, A. aegypti, T. castaneum, N. vitripennis, G. bimaculatus and D. melanogaster*) and have validated approximately 100 genes (**Figs. 3, 5, Fig. S10 and Table S4**). As a first step, we adapted and optimized the HCR-FISH protocol for each species, accounting for differences in tissue size and fixation requirements (**Methods**).

**Fig. 3.**
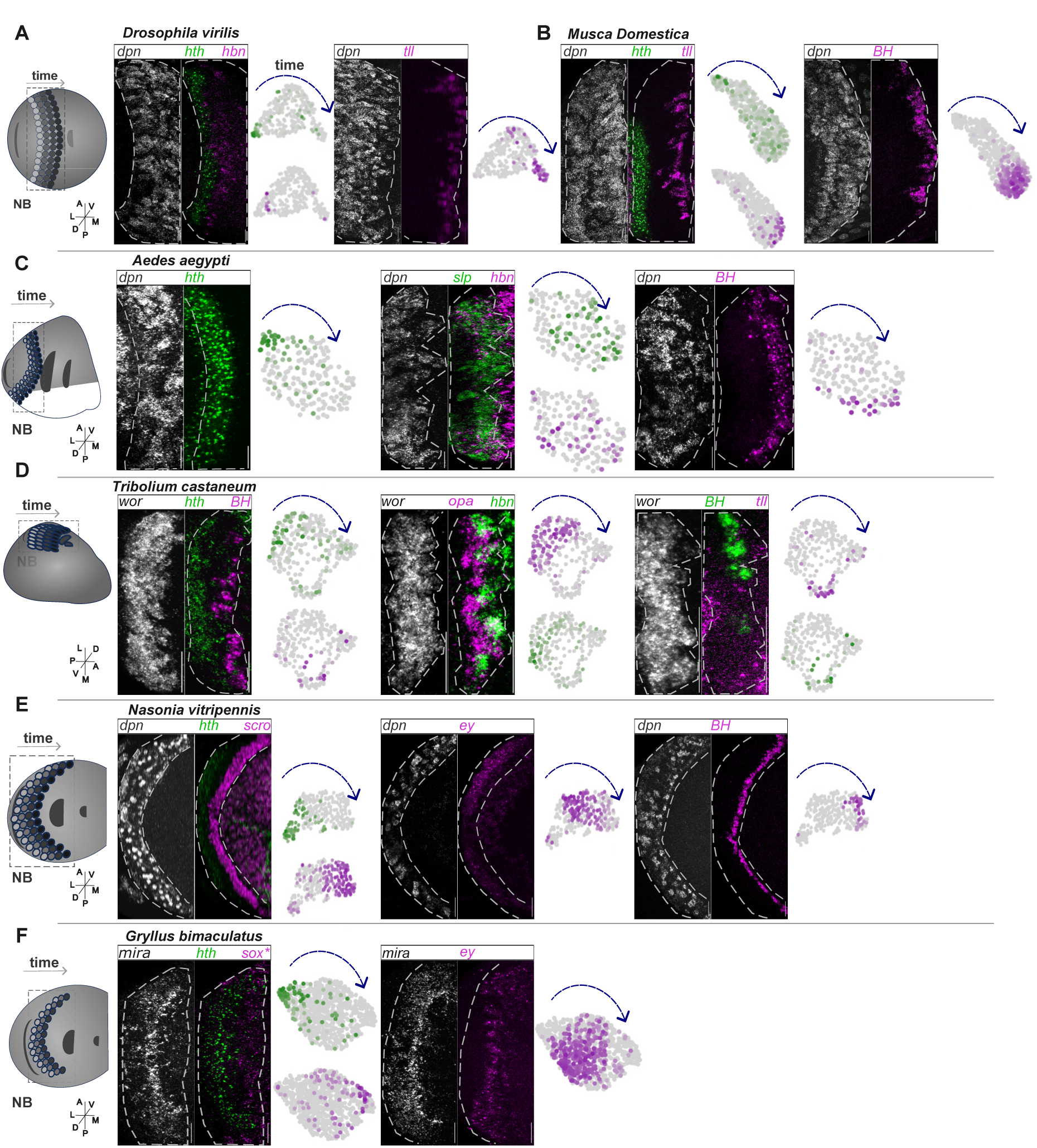
Experimental validation of conserved temporal transcription factors. **(A)** In *D. virilis*, *dpn* expression marks neuroblasts. Consistent with trajectory inference and UMAP analysis, *hbn* is expressed in neuroblasts that are older than those expressing *hth* (left), whereas *tll* is restricted to the oldest neuroblasts (right). **(B)** In *Musca*, *dpn* similarly labels neuroblasts. *hth* is expressed in young neuroblasts, while *tll* and *BarH1* (left and right, respectively) are confined to the latest temporal windows, as predicted by the UMAP. **(C)** In *Aedes*, *hth* marks young neuroblasts (left), *slp* and *hbn* show overlapping expression in a mid-temporal window (middle), and *BarH1* is expressed in old neuroblasts (right), consistent between HCR-FISH and UMAP analyses. **(D)** In *Tribolium*, coordinated expression patterns are observed between HCR-FISH and UMAP plots: *hth* and *BarH1* label young and old neuroblasts, respectively (left); *opa* and *hbn* are expressed in early temporal windows (middle); and *BarH1* and *tll* mark late neuroblasts (right). **(E)** In *Nasonia*, *hth* is expressed in young neuroblasts and *scro* in old ones (left), *ey* marks a mid-temporal window (middle), and *BarH* is restricted to late neuroblasts (right). **(F)** In *Gryllus*, *hth* is expressed in young neuroblasts, *D* in old neuroblasts (left), and *ey* in a mid-temporal window (right). For each panel a dashed line delineates the medulla neuroblast region, visualised by the respective neuroblast marker at each species. Images are z-projections of slices containing neuroblasts. Scale bar, 20µm.

We selected a subset of conserved tTFs - at least two to three per species - and examined their expression directly in medulla neuroblasts. In all cases, the HCR-FISH patterns recapitulated the expression inferred from the single-cell trajectories: early factors such as *hth*, mid-series factors such as *opa, ey*, *hbn or slp*, and late factors such as *scro*, *D, BarH,* and *tll* were detected in sequential domains along the medulla neuroblast layer (**Fig. 3A-F**, respectively and **Fig. S11**). This congruence confirms that trajectory inference accurately captures the *in vivo* temporal progression of neuroblast tTF gene expression and that the conserved core of the series is indeed deployed similarly during optic lobe development across insects.

### Divergent factors and strategies of evolution of the temporal series

While a conserved core of the temporal series can be traced back to the last common ancestor of holometabolous (and even hemimetabolous) insects, we also identified numerous instances of temporal transcription factor recruitment and loss across lineages (**Fig. 4A**). Rather than rewiring the program, these changes predominantly occur as modifications of a shared, conserved module, with most species retaining the major portion of the ancestral series. Strikingly, we observe more gains (34) than losses (12) of tTF expression (**Fig. 4B, Table S3**); since gene loss is more readily achievable involving fewer mutational steps, this bias suggests that the preferential recruitment of new temporal regulators reflects selection rather than evolutionary accessibility. Importantly, these events do not appear to correlate with eye and optic lobe size and estimated neuron numbers (*T. castaneum* has a markedly smaller optic lobe than *D. melanogaster*, whereas *G. bimaculatus* possesses a substantially larger optic lobe, consistent with differences in compound eye size, with approximately 80, 700 and 7000 ommatidia for *Tribolium*, *Drosophila* and *Gryllus* respectively ^58,59^.

**Fig. 4.**
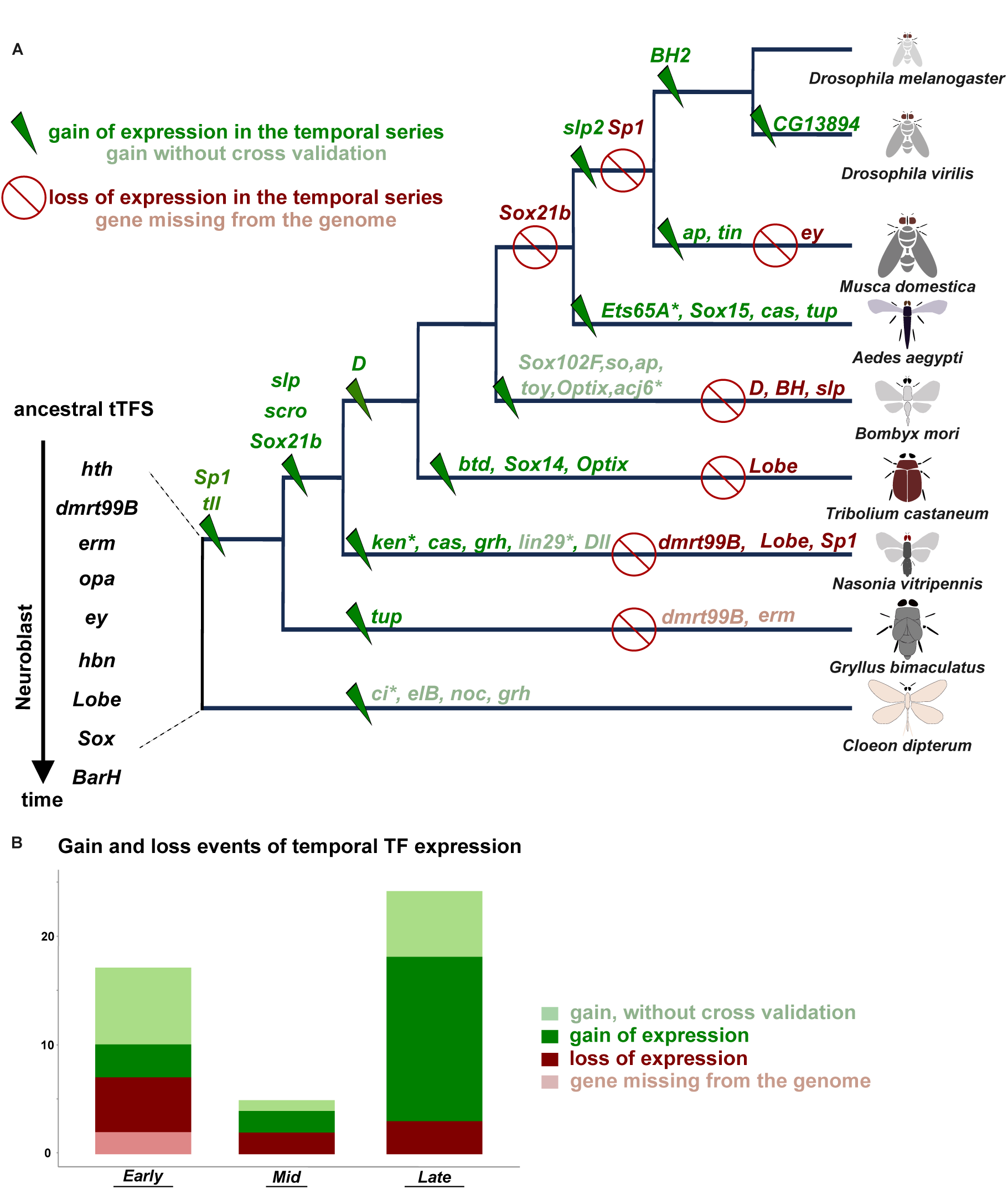
The evolutionary trajectory of the temporal series. **(A)** Phylogenetic tree depicting the inferred evolution of the medulla neuroblast tTF series from a conserved core present in the last common ancestor of hemi- and holometabolous insects (bottom left). Branches are annotated with inferred gains (green) and losses (red) of temporal expression, based on comparative single-cell and *in situ* analyses, as well as predicted gains lacking *in-situ* visualization (light green). Genes that are missing from the species genome are highlighted in light red. Terminal branches correspond to the nine insect species analyzed in this study. **(B)** Barplot comparing the instances of confirmed gains (green), unvalidated gains (light green), and losses (red), missing genes (light red) of tTF expression in the different insect lineages at early-, mid-, and late-stage neuroblasts.

**Fig. 5.**
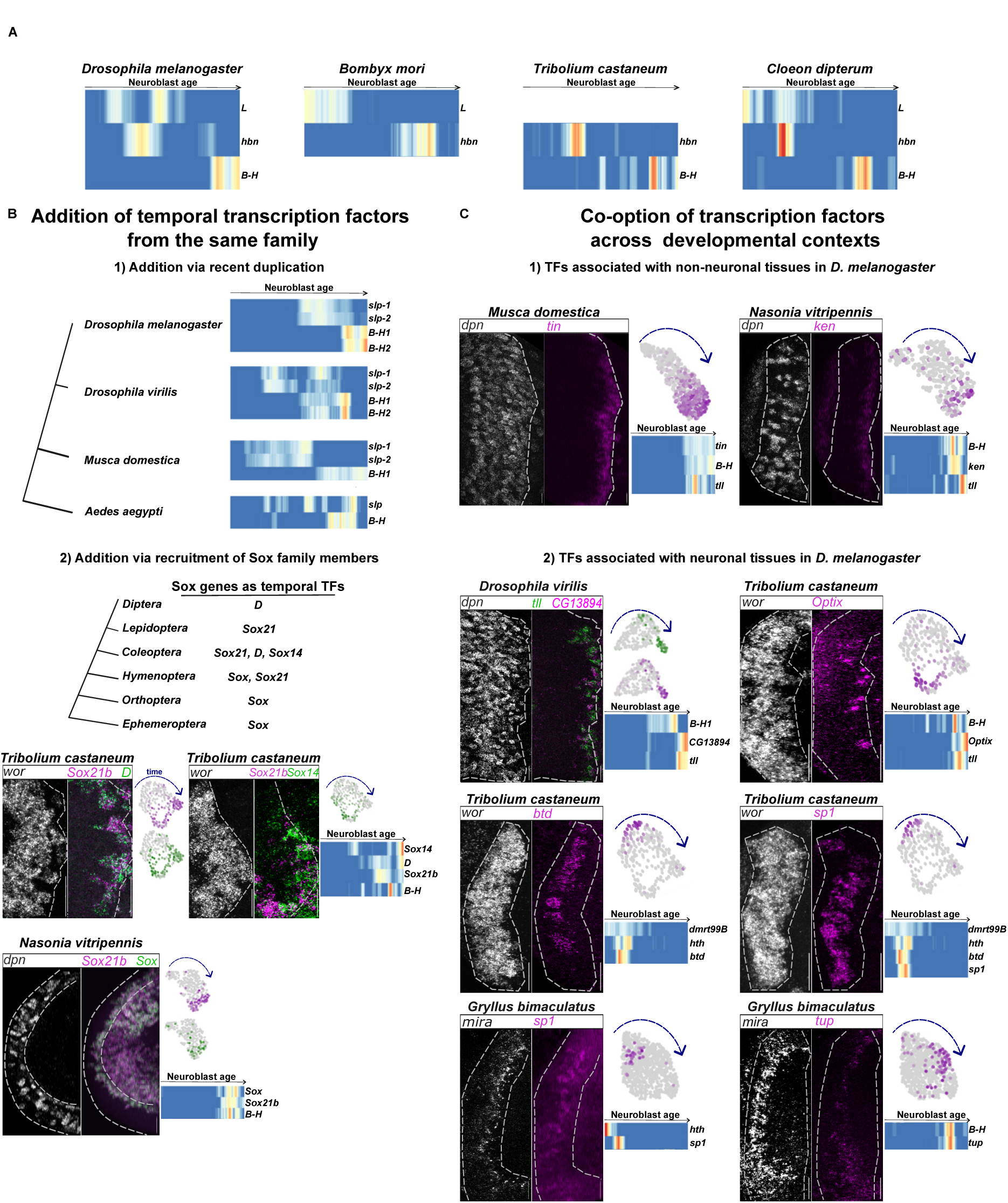
Mechanisms underlying the evolution of temporal patterning in insect neuroblasts. **(A)** Loss of tTF expression. Heatmaps showing conserved tTFs expressed both at *Drosophila* (left) and *Cloeon* (right), and examples of loss of temporal expression from either *Bombyx* or*, Tribolium*. Gaps in the heatmaps correspond to non-temporally expressed TFs, which are shown in **Fig. S9**. **(B)** 1) Addition of tTFs of the same family via recent duplication. Heatmaps showing the addition of *slp2* in the *Slp* window in *Musca* and *BarH2* in the *BarH* window in *Drosophila.* 2) Addition of Sox family members in the temporal series are shown in the different taxa. HCR-FISH of *Sox21b* (magenta), *D* (green-left), and *Sox14* (green-right) in *Tribolium* and *Sox21b* (magenta) and *Sox* (rgeen) in *Nasonia* show co-expression in neuroblasts, as indicated by the UMAP and heatmap. **(C)** 1) Addition of tTFs via co-option of TFs expressed in non-neuronal tissues in *D. melanogaster*: HCR-FISH against *tin* and *ken* (magenta) shows expression in old *Musca* and *Nasonia* neuroblasts (*dpn*- white), respectively, as indicated also by the respective UMAP and heatmap plots. 2) Co-option of TFs associated with neuronal tissues in *D. melanogaster*, as exemplified by *CG13894* (magenta) within the *tll* (green) window in *D. virilis*, *Optix* in *Tribolium*, *Sp1* and *btd* in young *Tribolium* neuroblasts, *Sp1* and *tup* in young/mid-age *Gryllus* neuroblasts. The dashed line delineates the medulla neuroblast region, visualised by the respective neuroblast marker at each species. Images are z-projections of slices containing neuroblasts. Scale bar, 20µm.

We then tested whether species-specific genes (**Table S1**) may contribute to temporal series innovations. We identified two species-specific genes among the temporally expressed transcription factors in *Gryllus*, a ZIC1 family TF sharing homology to *opa,* and a BH homolog (**Fig. 2D**). We also asked whether certain stages of the temporal series - early, mid, or late neuroblasts - were more prone to change; we find that while tTF expression loss is almost equally distributed, gains are enriched in the late part of the temporal series, indicating that the tempo of diversification increases as the temporal series progresses with the latest windows being the most divergent (**Fig. 4B, Table S3**).

Despite its apparent complexity, this phylogenetic reconstruction provides a conceptual map of temporal series evolution, which we then used to define general evolutionary mechanisms that act to diversify the series and, hence, neuronal diversity.

#### 1. Loss of expression of temporal transcription factors

One key mechanism, observed less frequently, involves the loss of temporal expression of specific tTFs, such as *BarH* in *B. mori* or *L* in *T. castaneum* (**Fig. 5A**). This likely represents a more drastic mode of network modification, given the severe phenotypes observed in tTF-mutant clones in *D. melanogaster* ^11^ and thus the apparently limited tolerance for removing core components of the regulatory network (**Fig. 4B**).

#### 2. Gain of temporal transcription factors from the same family

A more frequent mode of evolution involves the addition of temporal transcription factors within a largely conserved temporal framework (**Fig. 5B**). In this case, the overall sequence of the temporal program is maintained, while the molecular composition of individual windows is expanded. We identify two distinct modes of diversification:

##### 2a) Addition via recent gene duplication

Diversification can occur through the deployment of paralogous genes (**Fig. 2D**) within an established window, such as the expression of *slp2* alongside *slp1* in the Slp window in *Drosophila* and *Musca*, or *BarH2* alongside *BarH1* in the BarH window in *Drosophila;* notably, in *Musca*, *BarH2* is present in the genome, but is not expressed temporally **(Fig. 5B)**.

Because these changes occur without disrupting the global order of the temporal series, diversification within existing windows may represent a comparatively conservative mode of temporal program evolution, allowing incremental modification of temporal identity and potentially facilitating later shifts in tTF usage.

##### 2b) Recruitment of different Sox family members

We also observe cases where paralogous genes from the same transcription factor family (**Fig. 2D**) are co-expressed or substitute for one another within a temporal window (**Fig. 5C**). For example, members of the Sox family - *D*, *Sox14*, *Sox15*, and *Sox21b* - interchange their temporal expression in the late window across different species. In the hemimetabolous insects, *Gryllus* and *Cloeon*, a single *Sox* gene occupies this window. In *Nasonia*, *Sox21b* is also expressed temporally alongside another *Sox* gene (**Fig. 5C**), while in *Tribolium Sox21b* window is enriched by the addition of expression of *D* and *Sox14* (**Fig. 5C**). In Diptera, *Sox21b* expression is lost (**Table S4**), and *D* is the only *Sox* gene expressed, except in *Aedes* where *Sox15* is also expressed. This pattern suggests that functional redundancy within transcription factor families can facilitate smooth evolutionary transitions, allowing one paralog to replace or supplement another without disrupting temporal patterning.

#### 3. Co-option of transcription factors across developmental contexts

A third mechanism involves the co-option of transcription factors that are expressed in other developmental contexts or tissues in *D. melanogaster*, but are deployed as temporal transcription factors in neuroblasts of other insects (**Fig. 5C** and **Fig. S12)**.

##### 3a. Temporal TFs associated with non-neuronal tissues in D. melanogaster

We have identified factors that in *D. melanogaster* are mainly expressed outside the nervous system. The most striking example is *tinman (tin)*, which in *D. melanogaster* is restricted to the mesoderm ^60^, yet displays a clear and robust temporal expression pattern in medulla neuroblasts of *Musca* (**Fig. 5C**). Similarly, *ken*, which is not expressed in the *D. melanogaster* optic lobe and instead plays a key role in genital development ^61^, is expressed temporally in medulla neuroblasts of *Nasonia* (**Fig. 5C**). These observations were unexpected, as the recruitment of transcription factors with such distinct roles implies substantial rewiring of temporal window identity. This suggests that the temporal series is sufficiently flexible to incorporate factors from disparate regulatory contexts while maintaining a coherent developmental progression.

##### 3b. Temporal TFs associated with neuronal tissues in D. melanogaster

We also observed instances where transcription factors that, in *D. melanogaster*, are normally restricted to specific neuronal subtypes are acting as tTFs in other insects (**Fig. 5C**). For example, *Optix*, a spatial transcription factor of the optic lobe neuroepithelium in *D. melanogaster* ^62^, shows temporally regulated expression in *Tribolium* neuroblasts (**Fig. 5C**). We also confirmed the temporal expression of *Sp1* and *btd* in *Tribolium* neuroblasts, and *Sp1* and *tup* in *Gryllus* (**Fig. 5C**). Notably, these genes - including *elB*, *noc*, *Ets65A*, *tup, Dll and Sox21b* - have been previously identified as candidate terminal selectors in *D. melanogaster* ^27,63^. Terminal selectors are persistent transcription factors throughout differentiation and maintenance of neuronal functional identities, present in unique combinations specific to the neuronal identity ^64^. This pattern suggests a relatively simple evolutionary route for generating new tTFs: shifting the regulatory program of a gene from the post-mitotic neuron back into the neuroblast lineage allows it to acquire a temporal role without extensive rewiring of the existing temporal cascade.

### How does temporal patterning affect neuronal diversity?

We next asked how evolutionary differences in neuroblast temporal patterning may influence neuronal diversity. Notably, the datasets presented here are not sufficient to definitively assign mature neuronal identities to specific temporal windows, as the neurons we profile are not fully specified at these developmental stages. In previous work in *D. melanogaster*, assigning identities required integration with later, pupal-stage datasets to annotate neuronal cell types ^27^. Nevertheless, several neuronal transcription factors begin to be expressed shortly after neuron birth ^11^, providing an opportunity to examine early signatures of neuronal diversification.

We focused this analysis on *Musca domestica*, which displays some of the most pronounced deviations from the *D. melanogaster* temporal series, including the co-option of a mesodermal transcription factor (*tin*), the lack of expression of a *D. melanogaster* tTF (*ey*), and the deployment of one paralog (*BarH2*) within an existing temporal window. Together, these differences provide an informative case study to explore potential downstream consequences of temporal series evolution (**Fig. 6**).

**Fig. 6:**
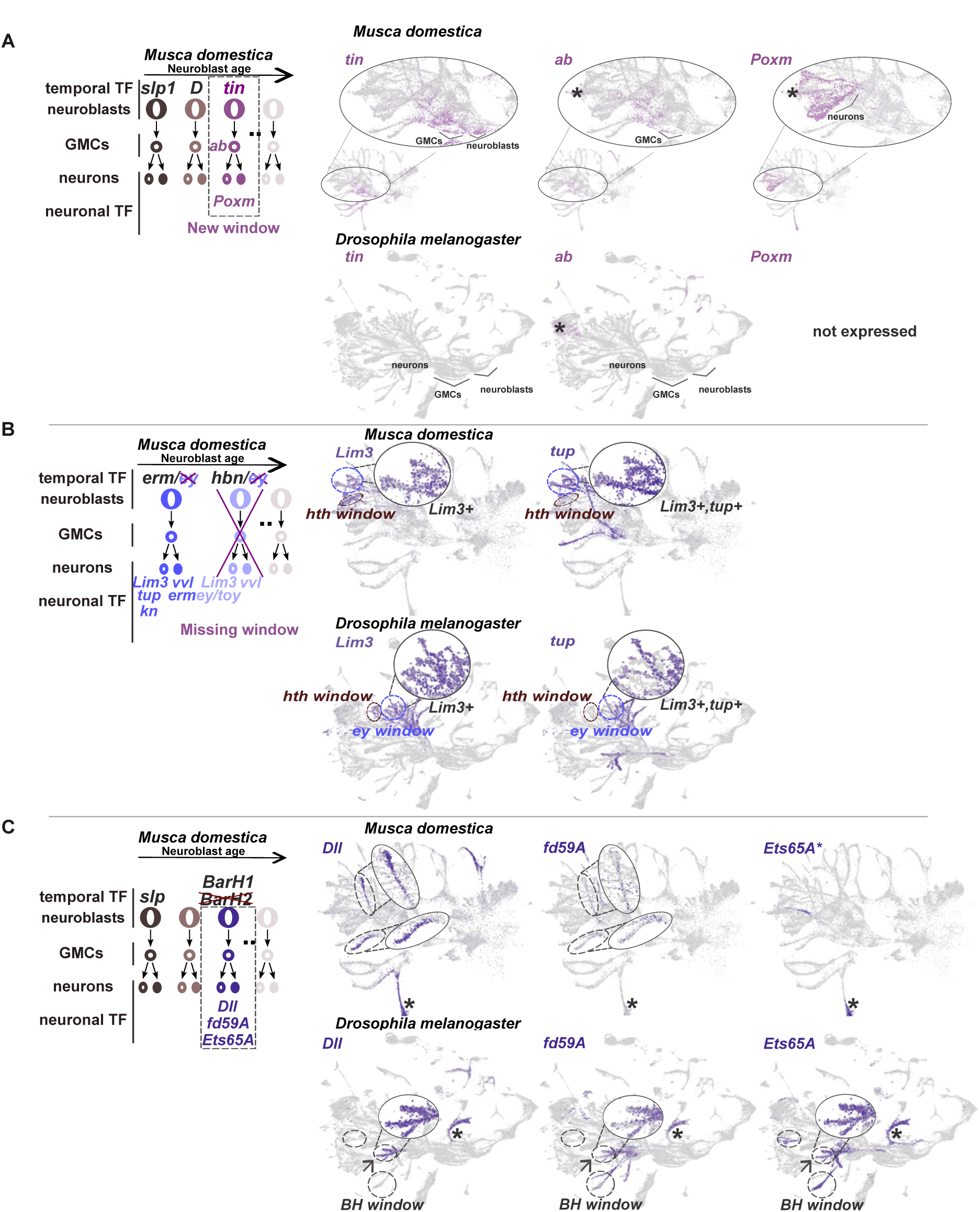
Impact of tTF evolution on early neuronal identity. **(A)** Neuroblasts, GMCs and neurons from the *tin* temporal window in *Musca* express *tin*, *ab*, and *Poxm,* respectively. UMAP plots showing the expression of *tin*, *ab*, and *Poxm* in the *Musca* and *Drosophila* developing brains. In *Musca, tin* is expressed in neuroblasts and GMCs from the same temporal window. The GMCs mark the beginning of the trajectories of neuronal progeny, along with *ab* expression and later *Poxm* in neurons. In *Drosophila, tin* and *Poxm* are not expressed at all, while *ab* is only expressed in few central brain neurons, marked by an asterisk (*). **(B)** The absence of the *ey* temporal window leads to the partial absence of its neuronal progeny in *Musca*. In *Drosophila*, Lim3+ neurons come either from the Hth temporal window (circled and highlighted as *hth* window) or from the Erm/Ey/Hbn windows (circled and highlighted as *ey* window). The neurons coming from the Ey/Hbn temporal window express *Lim3* and ey/*toy* (Lim3+,tup-) and are absent from *Musca*, while the ones coming from the Erm/Ey temporal window expressing *Lim3* and *tup* can be found in both *Musca* and *Drosophila*. **(C)** The absence of BarH2 expression in *Musca* correlates with absence of *Ets65A* expression and marked reduction in *Fd59A* expression levels, leading to the absence of co-expression of these two genes, together with *Dll* in *Musca*, while they are co-expressed in *Drosophila* medulla neurons (arrow), deriving from the D/BH window (all the descendent neurons from this window are circled in *Drosophila* and highlighted as *BH* window). There is co-expression of *Dll, Fd59A* and *Ets65A* in IPC-derived neurons, marked by an asterisk (*). The smaller asterisk next to *Ets65A* (*) indicates non 1:1 orthology relative to *Drosophila*.

We first examined the effects of *tin* co-option, whose expression in *Musca* coincides with that of *B-H1* and *tll* (**Fig. 5C**). In *Musca*, *tin* is robustly expressed in neuroblasts and GMCs, whereas it is entirely absent from the optic lobe in *D. melanogaster* (**Fig. 6A**). *tin* expression in the GMCs is accompanied by the expression of *ab*, which is also not expressed in *D. melanogaster*, with the exception of few central brain neurons (**Fig. 6A**). Using the expression of *tin* and *ab* at the root of neuronal trajectories, we identified neurons likely originating from the *tin* temporal window. Strikingly, these neurons express *Pox meso (Poxm)*, which is not expressed in *D. melanogaster* optic lobe neuroblasts or neurons (**Fig. 6A**). Notably, *Poxm* is involved in larval body wall muscle development ^65^ and is a direct target of *tin* in mesoderm development ^66^. Together, these observations suggest that the *tin* temporal window in *Musca* neuroblasts generates neurons with transcriptional identities not present in *D. melanogaster*, consistent with the emergence of species-specific neuronal populations.

We next examined the consequences of the absence of *ey* as a temporal factor in *Musca* (**Fig. 6B and Fig. S13**). In *D. melanogaster*, Lim3⁺ neurons represent the Notch^OFF^ progeny of the earliest neuroblast divisions and arise from several successive temporal windows (**Fig. 6B**): the Hth window (co-expressing *TfAP2* and *svp*; **Fig. S13**), the Hth/Opa window (co-expressing *run* and *tup*, the Erm/Ey window (co-expressing *kn*, *tup, ey*; **Fig. S13**), and the Ey/Hbn window (co-expressing ey/*toy*, but no *kn* or *tup*; **Fig. S13**) ^11^. In *Musca*, we identified all corresponding Lim3⁺ neuronal populations except those expressing *ey* (**Fig. 6B** and **Fig. S13)** and we did not identify a *toy* homolog (**Methods**). To determine whether this reflected the loss of a specific marker rather than the loss of an entire neuronal population, we searched for unaccounted Lim3⁺ neurons and found that all Lim3⁺ neurons are *tup*⁺, suggesting that they derived from the Erm temporal window in *Musca*, whereas the equivalent trajectories in *Drosophila* contain both *tup*⁺ and *tup*^−^*ey*⁺*/toy*⁺ neurons (**Fig. 6B and Fig. S13**). We also looked for the Notch^ON^ neurons that come from the Erm/Ey temporal window, which in *Drosophila* express *vvl and erm*; *ey* expression is not maintained in the neuronal progeny from this window ^11^. While *vvl* and *erm* are expressed in some *Musca* neuronal trajectories, these are markedly fewer than in *D. melanogaster* (**Fig. S13**), suggesting that these neurons come from the Erm temporal window rather than the Ey one. Together, these results provide strong evidence that the absence of an Ey temporal window in *Musca* is accompanied by a corresponding loss of its neuronal progeny.

Finally, we examined the potential effects of the absence of *BarH2* expression in *Musca* (**Fig. 6C**). In *D. melanogaster*, neurons born during the *BarH* window express transcription factors such as *Ets65A*, *Fd59A*, and/or *Dll* ^11^. In *Musca*, we identified neurons expressing *Fd59A* and *Dll* (**Fig.6C**) consistent with their origin from the BarH1 temporal window. However, *Fd59A* expression levels were markedly reduced, and *Ets65A* expression was entirely absent. On the other hand, we could observe the co-expression of the three markers in IPC-derived neurons, similar to the equivalent *D. melanogaster* cluster (**Fig.6C**). This suggests that the addition of *BarH2* expression in *D. melanogaster* may modulate downstream transcriptional programs by either amplifying the expression of neuronal transcription factors, consistent with findings in

*C. elegans*, where temporally acting TFs from the same family reinforce terminal selector activation ^67,68^, or by reshaping transcription factor combinations, thereby contributing to altered neuronal identity.

Although these observations are not conclusive in the absence of adult-stage neuronal data, they provide compelling evidence that evolutionary changes in the temporal transcription factor series can have tangible effects on the transcriptional identities of newly born neurons. Together, they support the idea that diversification of neuroblast temporal patterning through the loss or addition of temporal expression of TFs is a plausible mechanism for generating neuronal diversity across insect species.

## Discussion

In this study, we provide the first systematic analysis of how a temporal patterning program of neural stem cells evolves across insects. By combining single-cell transcriptomics, trajectory inference, and *in situ* analysis, we show that medulla neuroblasts across distantly related insect lineages share a conserved core of temporal transcription factors that was already present in the last common ancestor of hemi- and holometabolous insects more than 400 million years ago. At the same time, we uncover extensive variation in temporal series composition across species: our comparative analysis revealed a set of recurrent evolutionary strategies that shape temporal patterning programs across species and contribute to the diversification of temporal windows. These include the loss of temporal factors, the expansion of existing programs through the addition of transcription factors from the same family - either via recent duplication or positional substitution among transcription factor family members - as well as the co-option of transcription factors from other developmental or other neuronal contexts. Finally, neuronal transcription factor analyses suggest that such evolutionary changes in temporal patterning do translate into altered neuronal transcriptional identities. Together, our results reveal both the deep conservation and the evolutionary plasticity of temporal patterning programs, and establish temporal transcription factor series as a tractable substrate for the evolution of neuronal diversity.

It has not escaped our attention that each of the evolutionary strategies described above to reshape temporal patterning programs is likely to affect temporal series (and, hence, the neuronal diversity they generate) to markedly different degrees. For example, the co-option of the mesodermal transcription factor *tin* as a temporal factor (as well as its downstream target *Poxm* in neurons) in *Musca* is expected to profoundly perturb neuronal output and lead to the acquisition of new cell types ^69^. On the other hand, the substitution of a temporal factor by a closely related paralog is likely to produce much subtler changes; different *Sox* genes can have overlapping binding motifs ^70^ but also have distinct targets ^71^. We propose that this spectrum of effects provides evolution with a flexible framework to tune neuronal diversity while preserving the overall stability of developmental programs.

Consistent with this idea, although temporal transcription factor series described across neuronal systems are highly diverse, our analyses reveal a strikingly conserved landscape in the insect optic lobe, with evolutionary changes occurring primarily at the earliest and latest positions of the cascade, and being most enriched at the end. This pattern suggests that intermediate temporal states are under stronger constraint, likely reflecting requirements for the precise progression of neuroblast differentiation. Rather than being maintained solely through constraint, these conserved temporal transcription factors may represent a robust developmental scaffold that can be flexibly modified at its boundaries to generate neuronal diversity. From a gene regulatory network perspective, such flexibility may arise not only through changes in the composition of temporal transcription factors, but also through alterations in their downstream regulatory outputs. Temporal transcription factors activate, among other targets, terminal selectors ^63,72^, and changes in this readout could therefore reshape neuronal identity without necessarily rewiring the upstream temporal cascade. In support of this idea, although Lim3⁺/Ey⁺ neurons are absent from *Musca*, we identify Lim3⁺/Poxm⁺ neurons, suggesting that alterations in downstream effectors of temporally patterned lineages may contribute to evolutionary divergence in neuronal identity. Even though Poxm lacks the homeodomain characteristic of Pax6 family members such as Ey, it retains a paired domain and may therefore partially converge on related regulatory outputs; however, whether these neurons adopt equivalent functional identities remains unresolved.

Our findings further suggest that temporal series evolve not through the incorporation of species-specific genes (we only observed two instances of species specific tTFs that, nevertheless, share similarity with known tTFs), but via recombination and repurposing of existing regulatory components within this conserved toolkit. This may explain the existence of a complex gene regulatory network driving the progression of the series ^11^: through redundancy and modularity, the temporal series can generate neuronal diversity without disrupting the overall developmental trajectory. Such built-in redundancy provides evolutionary flexibility, allowing species to adapt and diversify while maintaining the core temporal program. In this way, the temporal series gene regulatory network acts as a buffer, permitting changes to accumulate without destabilizing its fundamental structure, thereby supporting both stability and adaptability in nervous system evolution.

While our study provides a broad view of temporal program evolution, some limitations remain. First, temporal transcription factors are often expressed at low levels and over narrow developmental windows, raising the possibility that additional regulators may have been missed; emerging single-cell technologies with increased sensitivity, as well as methods that better capture transcriptional dynamics, will be well suited to refine and extend the temporal series identified here. Second, our analysis necessarily depends on the quality of available genome assemblies and annotations, which vary across non-model species. While we mitigated this by improving gene models using single-cell mRNA sequencing data, and by generating *de novo* transcriptomes from publicly available RNA-seq data (e.g. for *Musca*) we also provide detailed pipelines that can readily be reapplied as genomic resources improve with optimized gene annotations. These considerations highlight both the current scope and the future potential of comparative temporal analyses to uncover the principles guiding neuronal diversification across species.

Our study indicates that insect optic lobe temporal series are highly conserved over hundreds of millions of years. This deep conservation mirrors the evolutionary stability of the insect eye ground plan, which is similarly maintained while accommodating lineage-specific novelty and innovation ^73^. While comparable data are not yet available for other insect neural stem cell populations, such as the ventral nerve cord or central brain, it is reasonable to hypothesize that similar constraints and evolutionary dynamics apply. In vertebrates, temporal transcription factor cascades in both the spinal cord and developing brain are likewise conserved across species ^12^ suggesting a general principle of broad conservation. However the temporal series differ massively between vertebrates and invertebrates and between different neuronal tissues. One possibility is that temporal series may have evolved rapidly after the divergence of vertebrates, followed by lineage-specific stabilization. Temporal patterning may not arise *de novo* in each neural stem cell population but could be adapted from preexisting regulatory modules. Spatial cues could be co-opted to generate temporal progression, and vice versa, allowing conserved regulatory logic to be redeployed in novel contexts; the expression of some optic lobe tTFs along the anteroposterior axis of the developing *Drosophila* procephalic ectoderm supports this scenario ^74^. Understanding how extensive diversification of temporal series is reconciled with the conservation of their underlying regulatory logic will be key to explaining the evolution of neuronal diversity.

## Methods

### Insect strains

*Drosophila melanogaster:* Canton-S; animals were reared at 25°C.

*Drosophila virilis*: 15010-1051.86; animals were reared at 25°C.

*Musca domestica*: M3 strain ^75^ (derivative of the wild-type multimarked strain aabys); animals were reared at 25°C.

*Aedes aegypti*: the inbred Paea strain ^76^; animals were received the days of experiments.

*Bombyx mori*: p50 strain (Silkworm Genetic Resource Database); animals were reared at 25°C. (Shortly after performing the experiments, we were informed by Silkwormbase that genetic analyses had revealed inadvertent contamination of the p50 strain with genomic material from *Bombyx mandarina*, most notably the W chromosome, likely arising from introgression of semiconsomic lines generated through interspecific crosses.)

*Tribolium castaneum*: wild type San Bernardino strain; animals were reared at 30°C.

*Nasonia vitripennis*: AsymCx wild-type strain; animals were reared at RT and 30°C .

*Gryllus bimaculatus*: wild-type strain with white-eyes ^77^; animals were reared at 28°C.

*Cloeon dipterum*: inbred wild type strain; animals were reared at 25°C.

### Edu staining and selection of neurodevelopmental stages

EdU proliferation assays were performed to identify developmental stages in different insect species that most closely correspond to the late third larval stage (L3) of *Drosophila melanogaster*, when neuroblasts are highly proliferative and medulla neurons are actively generated. For each species, stages immediately preceding pupation and metamorphosis were targeted, except for *Gryllus bimaculatus* and *Cloeon dipterum*, where neuroblasts and differentiated neurons coexist in nymphs immediately after hatching. Stage selection was based on available literature descriptions of optic lobe morphology, including the presence of Outer

Proliferation Centers and expansion of the medulla neuropil as indicators of ongoing neurogenesis. Accordingly, the following stages were analysed: *D. virilis* (late L3), *M. domestica* (late L3), *A. aegypti* (L4), *B. mori* (L5), *N. vitripennis* (late L3, white larvae, after gut cleaning, still moving), *T. castaneum* (L5 and early prepupa, when the neuropil is significantly expanded in beetles), all nymphal stages of *G. bimaculatus* ^78^ and early to mid–late nymphal stages of *C. dipterum* (2, 3, and 4 weeks after hatching, respectively). Brains were dissected in appropriate insect media (Schneiders insect medium (Sigma Aldrich), except for *B.mori* where Grace’s insect medium (GIBCO), was used) and incubated *ex vivo* ^79^ with EdU at a concentration of 20uM for a 4-hour pulse, followed by a 4-hour chase to allow detection of active proliferation. After incubation, samples were chemically fixed (3,7% FA/PBS (from a 37% FA solution, containing 10-15% methanol as stabilizer (Sigma Aldrich)), *D.virilis, M. domestica* 20 min RT, *A. aegypti, B.mori, G, bimaculatus, C. dipterum* ON at 4°C, *T. castaneum* 1h15min RT, *N. vitripennis* 2h15min RT), rinsed 3x PBST 0.3% (1xPBS with Triton-X-100 0.3%) washed 2x PBST 0.3% for 15 min each, and processed using the Click-iT EdU Imaging Kit. Brains were rinsed with PBST 0.3% and were incubated for 2 days with primary antibodies at 4 °C. Phalloidin (Sigma Aldrich) and anti-acetylated tubulin antibody were used at a concentration of 1/100 in PBST 0.03%. Brains stained with phalloidin were washed with PBST 0.3% and mounted in Vectashield. For the acetylated tubulin staining, brains were incubated with secondary antibodies for 2 days at 4 °C. The secondary antibodies were washed, and the brains were mounted in Vectashield for observation. The ZEISS LSM 980 laser-scanning confocal microscope or CSU-X1 Zeiss Spinning was used to verify active proliferation in the medulla neuropil.

### Single-cell dissociation protocols for the different insects and library preparation

#### Single-cell dissociation

Animals of all species were synchronized to the larval stages described above, except for *T. castaneum*, for which early prepupae were selected; *G. bimaculatus* (stages 1, 3, and 5); and *C. dipterum* (mid-nymphal stage). On the day of single-cell dissociation and library preparation, brains from both males and females were dissected at the selected stages (except for *C. dipterum*, where only female brains were used). To maximize biological coverage, we included individuals spanning mid- to late-wandering stages through early prepupae for all species, except *G. bimaculatus* and *C. dipterum*, for which nymphs were selected accordingly. All dissections were performed in ice-cold insect medium (Schneider’s insect medium for all species except *B. mori*, for which Grace’s medium was used), on pre-chilled Sylgard silicone plates using Dumont #5.5 forceps thoroughly cleaned with 70% ethanol and double-distilled water. Intact brains were gently transferred to glass staining dishes containing cold medium and kept on ice. Care was taken to prevent tissue damage and to minimize adhesion to pipette tips and low bind 1.5 ml Eppendorf tubes by applying in advance coating with DPBS + 0.04% BSA. Dissections were completed within 1 h 30 min. After dissection, optic lobes were separated from the rest of the brain using Vannas spring scissors (2 mm cutting edge; Fine Science Tools) or finely sharpened forceps when appropriate.

For single-cell RNA-seq, optic lobes were pooled (numbers per preparation indicated below) and enzymatically digested with collagenase (GIBCO; 2 mg/mL in the respective insect medium), either alone or in combination with dispase (GIBCO; 2 mg/mL). For selected *G. bimaculatus* stages, collagenase was supplemented with TrypLE (Sigma-Aldrich). Digestions were performed at 25–30°C for species-specific durations, followed by mechanical dissociation. After enzymatic treatment, only plasticware and pipette tips compatible with 10x Genomics recommendations were used. Optic lobes were transferred to Eppendorf DNA LoBind tubes (Fisher Scientific) containing 150 μL DPBS + 0.04% BSA and dissociated by vigorous pipetting (30–50 strokes per round), interspersed with 1 min incubations on ice, until no large tissue fragments remained. Suspensions were monitored under a dissecting microscope, and additional pipetting was performed only when large clumps persisted. Small clumps were not forcibly dissociated, as these likely corresponded to neuropil fragments rather than cell bodies; minimizing pipetting improves data quality. Cell suspensions were filtered through a 20 μm strainer (pluriSelect), followed by a wash with an additional 50 μL DPBS + 0.04% BSA, which was collected with the flow-through (including recovery from the reverse side when necessary). To minimize cell loss, collection tubes and strainers were pre-coated with DPBS + 0.04% BSA for at least 20 min on ice. Cell suspensions were assessed using a Luna FL cell counter (Logos Biosystems) to estimate the proportion of single cells versus aggregates and to determine cell concentration, except for *D. virilis*, where concentration was measured using a 0.02 mm deep hemocytometer. Conditions were as follows (number of pooled optic lobes (OLs); enzyme(s); temperature; duration): *D. virilis* (11 OLs; collagenase + dispase; 25 °C; 15 min), *M. domestica* (6 OLs; collagenase; 28 °C; 10 min), *A. aegypti* (10 OLs; collagenase + dispase; 28 °C; 20 min), *B. mori* (8 OLs; collagenase + dispase; 30 °C; 40 min), *N. vitripennis* (12 OLs; collagenase + dispase; 30 °C; 10 min; surrounding sheath removed prior to dissociation), *T. castaneum* (60 OLs; collagenase + dispase; 30 °C; 15 min), *C. dipterum* (9 OLs; collagenase + dispase; 28 °C; 70 min), and *G. bimaculatus* nymphal stages: st1 (21 OLs; collagenase; 28 °C; 10 min), st3 (10 OLs; collagenase + TrypLE; 28 °C; 30 min), and st5 (5 OLs; collagenase + TrypLE; 28 °C; 35 min). Digestion was performed on isolated optic lobes for all species except *N. vitripennis*, for which whole brains were digested and optic lobes were subsequently isolated for downstream processing.

### Library preparation

Three independent single-cell mRNA-seq libraries were generated for each developmental stage from a single pooled single-cell suspension prepared by merging optic lobes from multiple individuals of the same species (i.e., the same suspension was split to generate the three libraries). In total, 30 libraries were produced (3 libraries per species, except *G. bimaculatus* where 9 libraries were prepared for three nymphal stages). Library preparation followed the 10x Genomics Chromium Next GEM Single Cell 3ʹ Reagent Kits v3.1 (Dual Index) protocol, which captures the 3ʹ end of polyadenylated mRNA transcripts. Libraries were sequenced by Novogene on an Illumina NovaSeq platform using PE150 (paired-end 2×150 bp), targeting ∼30,000 reads per cell.

#### GeneExt and Cell Ranger

GeneExt ^38^ was used to mitigate incomplete genome annotations in non-model species, where missing 3′ gene ends can reduce read/UMI assignment in 3′-based single-cell RNA-seq datasets. GeneExt performs peak calling on aligned reads to detect transcriptional signals extending beyond annotated gene boundaries, and extends nearby genes’ 3′ regions accordingly, thereby improving UMI capture; it can also use unassigned intergenic peaks to propose putative novel genes (orphans). GeneExt was run on the initial BAM files generated with cellranger v7.1.0 ^80^ (--expect-cells=15000) to produce an improved genome annotation file, and cellranger was then re-run for each library using the updated annotation to generate refined BAM files and gene expression matrices. To maximize evidence for extension calling, BAM files from all sequenced developmental and adult stages were merged per species using samtools v1.13 ^81^ and processed with GeneExt using a subsample parameter (--subsamplebam) of 100 million reads to accelerate computation as well as the orphan (--orphan) parameter to report candidate novel genes; orphan predictions were subsequently filtered out and retained only for future exploratory analyses. The final, filtered improved annotation (integrating information across stages) was used for the second cellranger run to generate updated BAM files and final gene expression matrices with enhanced transcriptional coverage (**Fig. S3**). Using this approach, GeneExt extended 3′ regions for 4260/15621 genes in *D. virilis* (median extension 562 bp), 4893/20747 in *M. domestica* (937 bp), 3994/19269 in *A. aegypti* (1427 bp), 5991/16687 in *B. mori* (3115 bp), 3094/14492 in *T. castaneum* (804 bp), 3876/15025 in *N. vitripennis* (1106 bp), 5629/17869 in *G. bimaculatus* (5423 bp), and 5403/16357 in *C. dipterum* (705 bp).

#### Single-cell mRNA sequencing data analysis

The libraries were analyzed using Seurat v5 ^82^. For each dataset, a Seurat object was created from the Cell Ranger gene-barcode count matrix (UMI expression matrix), using all droplets classified as cells by Cell Ranger’s cell-calling filters. Genes expressed in fewer than three cells were removed, and cells with fewer than 200 detected genes were excluded. Objects were further filtered by applying library-specific thresholds of genes per cell (nFeatures/cell) to optimize data quality. Filtering decisions were guided by inspection of nFeatures distributions visualized with histograms and violin plots (**Fig. S4**) for each library, and some low-complexity droplets were intentionally retained to maximize available information. DoubletFinder ^83^ was used to remove doublets in libraries with more than 20000 cells after filtering (*B. mori* and *M.domestica*). Standard Seurat preprocessing functions - including NormalizeData, FindVariableFeatures, and ScaleData - were applied with default parameters, followed by dimensionality reduction using PCA and UMAP (100-150 principal components depending on the dataset). The three libraries generated from the same pooled suspension for each species were integrated using the FindIntegrationAnchors and IntegrateData functions. After integration, residual groups of low-quality cells that showed poor integration and were characterized by reduced nFeatures compared to the main cell population were removed only for visualization purposes.

#### Phylome reconstructions

A phylome is the collection of phylogenetic trees for each gene in a genome. We reconstructed a phylome for each of the species included in the analysis and added *Daphnia pulex* to serve as an outgroup. Proteomes were retrieved from public repositories and formatted using phylomeDB-compliant proteome codes as follows (species; proteome code; source; phylomeID): *Drosophila melanogaster* (DROME; FlyBase v6.49; 938), *Drosophila virilis* (DROVI; NCBI assembly GCF_003285875.2; 939), *Musca domestica* (MUSDO; VectorBase-61; 940), *Aedes aegypti* (AEDAE; VectorBase-61; 941), *Bombyx mori* (BOMMO; Ensembl Metazoa release 55; 942), *Tribolium castaneum* (TRICA; Ensembl Metazoa release 55; 943), *Nasonia vitripennis* (NASVI; Ensembl Metazoa release 56; 944), *Gryllus bimaculatus* (GRYBI; *G. bimaculatus* genome resource; 945), *Cloeon dipterum* (proteome code 197152; NCBI assembly GCA_902829235.1; 946), and *Daphnia pulex* (DAPPPU; retrieved from phylomeDB; 947).

We used an automated pipeline that uses the same process to reconstruct gene trees as one would do manually^84^. First the proteome database was reconstructed using the 10 species and formatting the codes to phylomeDB format. Then a phylome was reconstructed starting from each one of the species. For each species a blastp search was performed between each gene in their genome and the proteome database. Blast results were filtered using an e-value threshold of 1e-05 and an overlap threshold of 50%. The number of hits was limited to the 200 best hits for each protein. Then six different multiple sequence alignments were reconstructed using three programs (Muscle v3.8.1551^85^, mafft v7.407^86^ and kalign v2.04^87^ and aligning the sequences in forward and in reverse. From this group of alignments a consensus alignment was obtained using M-coffee from the T-coffee package v12.0^88^. Alignments were then trimmed using trimAl v1.4.rev15 (consistency-score cut-off 0.1667, gap-score cut-off 0.9) ^89^. IQTREE v1.6.9^90^ was then used to reconstruct a maximum likelihood phylogenetic tree. Model selection was limited to 5 models (DCmut, JTTDCMut, LG, WAG, VT) with freerate categories set to vary between 4 and 10. The best model according to the BIC criterion was used. 1000 rapid bootstraps were calculated. All trees and alignments were stored in phylomeDB ^54^ with phylomeIDs ranging from 938 to 947 (http://phylomedb.org).

A species tree was reconstructed using a gene concatenation approach. To do that we first selected which genes in the phylome of *D. melanogaster* were found in single copy in all species. 547 such genes were selected. The alignments for those genes were then concatenated into a single multiple sequence alignment producing an alignment of 342,569 amino acid positions. IQTREE v1.6.9^90^ was used to reconstruct the species tree using LG as the model. 1000 rapid bootstrap calculations were performed.

All trees across the phylomes were rooted using the species tree reconstructed above. To do so, for each tree the seed species was taken and the other species were numbered from closest to farthest according to the species tree. Then the root was placed at a node containing the farthest related species in the gene tree. Once the trees were rooted, all possible orthology and paralogy relations were extracted from each tree using the species overlap algorithm as implemented in ETE3^91^. For each pair of sequences detected as orthologous in any of the trees we computed a score which divides the number of times the pair appeared as orthologs across all trees by the number of times the pair of sequences was found as either orthologs or paralogs (CS). If the CS was above 0.5 the pair of sequences were considered orthologous. Using this information, we classified each gene relative to *Drosophila* genes as one-to-one, one-to-many, many-to-one, many-to-many homologs, or as species-specific (**Table S1**).

#### Identification of transcription factors in insect genomes

A transcription factor (TF) candidate list was generated by screening the genome annotation file of each species using a curated reference list of *D. melanogaster* TF names and TF-associated keywords (“homeobox”, “zinc finger”, “bHLH/HLH”, “leucine zipper”, “POU”, “LIM”, “forkhead”, “nuclear receptor”, “DNA-binding”, “transcription”) as functional cues. Matches were performed in a case-insensitive manner while accounting for minor formatting differences such as hyphenation, and genes with at least one match were retained as putative TF candidates, with the corresponding TF term recorded. Additionally, we queried the similarity-relationships list across the phylomes to identify proteins showing sequence similarity to any known *D. melanogaster* TF, further expanding the candidate lists. This inclusive strategy allowed recovery of maximal TF-related information, accepting the possibility of false positives in order to capture as many potential candidate TFs as possible. In the resulting lists, *D. melanogaster* TF names were propagated to the corresponding orthologs in each species when orthology relationships were established.

#### Identification of temporally expressed transcription factors

Initially, we identified medulla neuroblast clusters in the integrated object based on expression of neuroblast markers such as *mira, dpn,* and *wor*, while lamina and lobula plate neuroblast pools were excluded using complementary markers. In *Cloeon* only *dpn* was used, because it was not possible to identify a corresponding locus to *mira* or *wor* in the genome, through annotation, or the available phylomes, or reciprocal blast, or Flybase homologs for other insect species. HCR-FISH validations allowed the identification of the limits between lamina and medulla clusters for *Aedes* and *Nasonia*, leading to subclustering to remove lamina cells. For each species, clusters of interest were extracted and developmental trajectories were inferred using Slingshot ^55^, which defines lineages from low-dimensional embeddings and orders cells in pseudotime. For each species, candidate temporal transcription factors (tTFs) were then investigated by testing for genes with pseudotime-associated expression changes using PseudotimeDE^56^, focusing on the species-specific TF candidate lists. Because our goal was to detect temporal restriction (genes expressed in only part of the trajectory) rather than broadly dynamic expression across the entire trajectory, we implemented additional filtering steps to prioritize genes with windowed expression. Pseudotime values were rescaled to 0–1 and cells were binned into 20 equally spaced pseudotime intervals to generate a cell-by-interval assignment matrix. Candidate tTFs were then filtered using log-transformed counts with two criteria: (a) a reach-zero filter retaining genes whose mean expression dropped to near-background levels in at least one pseudotime interval (mean < 0.1), while requiring expression in at least 5% of all cells; (b) a minimum-expression filter requiring sufficient overall expression (mean non-zero log-expression above a defined threshold; values of 0.4 and 0.45 were tested) and at least 5% non-zero cells to remove lowly expressed/noisy genes. The PseudotimeDE parametric p-value was also considered to account for uncertainty in pseudotime estimation. Exceptionally for *Gryllus*, only the approximation to 0 was applied, since the levels of expression for the TFs with temporal expression were lower in comparison to the other species. Genes passing both filters were retained and further inspected. Candidates with clearly temporally restricted patterns (i.e., not expressed throughout the trajectory) were prioritized as putative tTFs. In addition, we considered that bona fide tTFs are typically robustly expressed in neuroblasts relative to false-positive candidates, and the parametric p-value from pseudotimeDE was used as an additional selection criterion, printed at each plot of candidate tTFs (annotated as para).

#### Missing and *ey* annotation in the *Musca* genome

To determine whether tTFs known from the *Drosophila melanogaster* optic lobe were genuinely absent from other insect neuroblast temporal series or instead missing due to incomplete genome annotations, we applied different strategies depending on the availability of bulk RNA-seq datasets in public repositories.

##### a) Identification of *Musca eyeless*

Head RNA-seq datasets from *Musca domestica* were retrieved from the Gene Expression Omnibus (GEO) under accession numbers GSM3611342 and GSM3611234. Reads were assembled de novo using the Trinity transcriptome assembler ^92^ to generate a *Musca* head transcriptome assembly.

To identify the *Musca domestica eyeless (ey)* homolog, the *Drosophila melanogaster ey* protein sequence was used as a query in a tblastn search against the assembled *Musca* transcriptome. The transcript with the highest similarity score was selected as the putative *Musca Ey* candidate. Orthology was further assessed by blastp analysis of the predicted *Musca Ey* protein sequence against the *D. melanogaster* protein database. Reciprocal best-hit analysis identified *D. melanogaster eyeless* as the top match, supporting its annotation as the *Musca ey* ortholog. This assignment was further validated by phylogenetic analysis (**Fig. S10B**).

The identified *Musca ey* transcript sequence was subsequently used in a tblastn search against the *Musca domestica* genome assembly to determine the genomic location of the locus, which was then manually incorporated into the genome annotation file.

##### b) Identification of missing genes in other species

For several candidate genes absent from the *Gryllus* datasets, including *dmrt99B, erm, scro,* and *slp*, no bulk RNA-seq data were available in public repositories. To assess whether these genes were present in the species, tblastn searches were performed using the corresponding *D. melanogaster* Erm, Scro, and Slp protein sequences as queries against the available *Gryllus* transcriptome assembly. As no reliable hits were identified, these genes were considered absent from the currently available *Gryllus* transcriptomic resources.

#### Hybridization Chain Reaction Fluorescence *In Situ* Hybridization (HCR-FISH)

To generate the final list of candidate temporal transcription factors, we validated expression patterns by HCR-FISH for 100 genes (**Table S4**). Probe sets were either purchased from Molecular Instruments or designed with the HCR probe generator (https://github.com/rwnull/insitu_probe_generator) and synthesized by IDT. HCR-FISH was performed following published protocols with the addition of a blocking step ^94–96^. Briefly, brains from *D. melanogaster*, *D. virilis*, *M. domestica*, *A. aegypti*, *T. castaneum*, *N. vitripennis*, and *G. bimaculatus* were dissected in Schneider’s insect medium, fixed in 4% FA/PBS (from a 16% FA stock, methanol free, ThermoFisher) for 20 min, rinsed 3x and washed 2×15min in PBST (PBS + 0.1% Triton X-100). While *Drosophila* brains were processed directly, brains from the other species were dehydrated in 100% methanol to improve tissue penetration and allow long-term storage, and were rehydrated prior to the hybridization step through a methanol series in PBST (100%, 75%, 50%, 25% in PBST 0.1%). On day 1, samples were pre-hybridized in 200 µL pre-warmed probe hybridization buffer for 30 min at 37°C in glass staining dishes, ensuring tissues were fully immersed. Probe solution was prepared by adding 1–4 pmol of each probe set to 200 µL of pre-warmed hybridization buffer, supplemented with denatured salmon sperm DNA (0.1 mg/mL; Sigma D7656; boiled for 10 min and cooled on ice for ≥5 min) and fresh BSA (2 mg/mL), and samples were incubated overnight at 37°C in the probe and blocking solution (sealed). On day 2, samples were washed four times for 15 min each in pre-warmed probe wash buffer at 37°C, followed by three 5-min washes in 5×SSCT (0.3% Triton) at room temperature. Samples were then pre-amplified for 30 min in amplification buffer at room temperature. Hairpins were prepared by snap-heating 12 pmol each of h1 and h2 at 95°C for 90 s, cooling for 30 min at room temperature in the dark, diluting each in 100 µL amplification buffer, and combining to a final volume of 200 µL; samples were incubated overnight in hairpin solution at room temperature (protected from light). On day 3, hairpins were removed and samples were washed in 5×SSCT at room temperature (2 × 5 min, 2 × 30 min, and 1 × 5 min). The brains were mounted in Vectashield and imaged on a ZEISS LSM 980 laser-scanning confocal microscope, using a ×40 glycerol objective. Images were processed in Fiji.

## Data availability

All raw and processed data referenced were uploaded to GEO: accession number GSE333138. All related code is available in GitHub (https://github.com/NikosKonst/EvoTempoPattern).

## Acknowledgements

We thank Claude Desplan, Vanessa Ribes, Babis Galouzis, Félix Simon, and Carlos Martin Blanco for comments on the manuscript. We thank all colleagues who kindly provided animals and expertise for insect rearing: Leo Beukeboom, Anna Rensink, Daniel Bopp, Claudia Brunner-Barios, Siegfried Roth, Matthias Pechmann, Eloise Muller, Isabel Almudi, Gregor Bucher, Marie Vazeille and Anna Bella Failloux, and Manon Monier. We also thank members of the Konstantinides lab for valuable discussions, especially Rebekah Rickeburg for help with animal rearing and Isabel Holguera for sharing expertise in mounting of larval brains and imaging. We thank Joel Marchand for informatic assistance. We also acknowledge the ImagoSeine core facility at the Institut Jacques Monod, a member of France-BioImaging (ANR-10-INBS-04) and certified by GIS-IBiSA.

## Author contributions

N.K. and K.F. designed the project. K.F. established and maintained all animal cultures in the laboratory. K.F. designed protocols and performed experiments, imaging, and single-cell data generation and downstream computational analysis. K.F., C.L., and J.J.L. developed the code for temporal transcription factor selection. E.I. improved genome annotations and processed single-cell sequencing reads. K.F. and C.J.L.M. performed HCR-FISH and imaging. M.M.-H. and T.G. performed phylogenomic analyses. K.F. and N.K. interpreted the data and wrote the manuscript. All authors edited and approved the final manuscript.

Conceptualization: K.F., N.K.; Investigation K.F., E.I., C.J.L.M., C.L., M.M.-H.; Methodology K.F.; Funding acquisition N.K.; Supervision K.F., T.G, J.J.L., N.K.; Writing – original draft K.F., N.K.; Writing – review & editing K.F., E.I., C.J.L.M., C.L., M.M.-H., T.G, J.J.L., N.K.

## Additional Information

Correspondence and requests for materials should be addressed to Nikolaos Konstantinides.

## Supplementary Figure legends

**Fig. S1. Life cycle schematics of insects used in this study.**

(A) Schematics of the lifecycle of the holometabolous insect species that were used in this study. The optimal temperature, developmental time, and number of larval stages varies between insects.

(B) Schematics of the lifecycle of the hemimetabolous insect species that were used in this study. These insects lack larval and pupal stages. The number of nymphal stages and developmental time differs between the two insects.

**Fig. S2. EdU incorporation identifies the peak neuroblast proliferative stage.**

(A) EdU staining in *A. aegypti* early L4 stage.

(B) EdU staining in *B. mori* late L4 stage.

(C) EdU staining in *T. castaneum* L4 and L5 stages.

(D) EdU staining in *N. vitripennis* early L3 stage.

(E) EdU staining in *G. bimaculatus* stages 3, 4, 5, and 7.

(F) EdU staining in *C. dipterum* early and mid-late nymphs.

Medulla proliferating cells (cells marked in green) are circled with dashed lines. Scale bar, 20µm.

**Fig. S3. GeneExt pipeline and results.**

(A) GeneExt pipeline that was followed for the improvement of genome annotation in the eight insect species.

(B) Violin plots of gene extension lengths and numbers of extended genes across eight insect species.

**Fig. S4. Quality control and filtering of insect single-cell mRNA sequencing libraries.** For each of the thirty libraries, a histogram showing the distribution of detected features per droplet/cell (left) and violin plots of the number of detected features per cell before (middle) and after (right) filtering are shown. The red dotted line indicates the library-specific filtering threshold, droplets left to the dotted line were removed.

**Fig. S5. Cell-class identification across insect species by marker gene expression.**

UMAP projections are shown for *D. virilis* (A), *M. domestica* (B)*, A. aegypti* (C)*, B. mori* (D)*, T. castaneum* (E)*, N. vitripennis* (F)*, G. bimaculatus* (G), and *C. dipterum* (H), coloured by the expression of selected marker genes used to identify major neural cell classes, including neuroepithelial cells (NE), neuroblasts (NB), ganglion mother cells (GMCs), and glial cells. These markers enable the assignment of cells to distinct cell classes within each species.

**Fig. S6. Neuropil identification across insect species by marker gene expression.**

UMAP projections are shown for *D. virilis* (A), *M. domestica* (B)*, A. aegypti* (C)*, B. mori* (D)*, T. castaneum* (E)*, N. vitripennis* (F)*, G. bimaculatus* (G), and *C. dipterum* (H), coloured by the expression of selected marker genes used to identify different neuropils. These markers enable the assignment of cells to the lamina (highlighted in the blue circle) and lobula plate (highlighted in the purple circle).

**Fig. S7. Phylome reconstruction and ortholog identification.**

(A) Percentage of genes in the species of the x-axis that have at least one ortholog in the species of the y-axis. As the relationship between orthologs may not be one-to-one and the proteome sizes vary the graph is not symmetrical.

(B) Number of *D. melanogaster* genes that have one-to-one (dark blue), one-to-many (cyan), many-to-one (green), many-to-many (pink) orthologs or are *D. melanogaster*-specific (grey) for each of the species used in the phylome reconstruction.

**Fig. S8. Pipeline for the identification of candidate tTFs.**

(A) For each gene, expression was plotted along pseudotime. Two filters were applied to ensure temporal expression (i.e., present in specific segments of the trajectory but absent in others and adequate expression). Genes passing these criteria were further evaluated based on expression levels and PseudotimeDE p-values, widespread or localised expression and selected candidates were subsequently tested by HCR-FISH. In this example *tll* represents a confident candidate tTF, while *elB* requires further validation, since it is lowly expressed. *ap* does not pass the gene expression filter so is discarded, and *bbx* though it passes the filter for temporality has widespread expression and is considered a false positive, therefore discarded.

(B) HCR-FISH analysis of *elB* and *tll* (purple) in the developing optic lobes of *D. virilis*. Neuroblasts are visualized by *dpn* expression (white), the dashed line delineates the neuroblast region. The expression pattern of *elB* is not temporal, as opposed to that of *tll*; therefore, this candidate tTF was discarded. For each panel a dashed line delineates the medulla neuroblast region, visualised by. Images are z-projections of slices containing neuroblasts. Scale bar, 20µm.

**Fig. S9. Expression of all candidate tTFs along the neuroblast trajectory.**

Dynamic gene expression along the neuroblast pseudotime trajectory (left) and UMAP visualization of gene expression in neuroblasts (right) are shown for all candidate tTFs in all eight species. Genes highlighted in red or green were tested by HCR-FISH; among these, genes highlighted in red represent species-specific tTFs. Genes highlighted in light or dark grey were not cross-validated by HCR-FISH; those in dark grey correspond to *D. melanogaster* tTF orthologs. Genes below the horizontal dark line were eliminated from the candidate tTF list either because cross-validation did not support their temporal expression or because they were expressed at low levels. The asterisk (*) refers to non 1:1 ortholog genes.

**Fig. S10. Identification of the *ey* gene in *Musca domestica*.**

(A) Pipeline to computationally identify the *ey* trascript in *Musca* genome. Available RNA-seq data from *Musca* head were used to generate a transcriptome. tblastn of *Drosophila* Ey against *Musca* transcriptome revealed a highly confident hit. Reciprocal blast against *Drosophila* returned as best hit Ey, and subsequent search in *Musca* genome revealed the *ey* gene.

(B) Phylogenetic reconstruction of Ey and Toy proteins of all 8 species, including the newly identified *Musca* Ey, positions this hit with the other Ey orthologs and not with Toy ortholog proteins. *Gryllus*, *Aedes*, and *Cloeon* do not have an identified 1:1 Toy ortholog, or homolog. Nodes are collapsed when bootstrap values are below 50, the branches are thicker when the bootstrap is high and are coloured in blue for speciation events and in red for duplication events. Scale bar represents the estimated number of substitutions per site along each branch.

(C) HCR-FISH in the eye-antennal imaginal disk of L3 larvae further confirms that *ey* is restricted in the eye region, while *hth* counterstains the whole structure. Images are z-projections of the slices that express *ey*.

(D) UMAP plot of *ey* in the *Musca* dataset shows its absence of expression in the neuroblasts, while neuronal expression can be seen in the tips of some of the neuronal trajectories.

(E) HCR-FISH in the *Musca* developing optic lobe of L3 larvae confirms that *ey* expression is absent from neuroblasts, while being present in neurons. Images are z-projections of the slices that express *ey*.

Scale bar, 20 μm.

**Fig. S11. Experimental validation of conserved temporal transcription factors.**

HCR stainings from **Fig.3** are presented with individual channels separated and merged with neuroblast markers per species. Additional conserved temporal gene expression patterns are presented per species.

(A) In *D. melanogaster, hth* (left, green), *tll* (left, magenta), and *scro* (right, magenta) HCR-FISH in neuroblasts (*dpn,* white) agrees with the antibody stainings ^11^.

(B) HCR-FISH expression of *hth* (left, green), *hbn* (left, magenta), and *tll* (right, magenta) in *D. virilis* neuroblasts (*dpn*, white).

(C) HCR-FISH expression of *hth* (left, green), *tll* (left, magenta), *erm* (middle, magenta) and *Bar-H* (right, magenta) in *M. domestica* neuroblasts (*dpn*, white).

(D) HCR-FISH expression of *hth* (left, green), *slp* (middle-left, green), *hbn* (middle-left, magenta), *Bar-H* (middle-right, magenta), and *tll* (right, magenta) in *A. aegypti* neuroblasts (*dpn*, white).

(E) HCR-FISH expression of *hth* (left, magenta), *D* (left, green), and *ey* (right, magenta) in *G. bimaculatus* neuroblasts (*mira*, white).

(F) HCR-FISH expression of *opa* (left, green), *scro* (left, magenta), *Bar-H* (middle, magenta), and *tll* (right, magenta) in *N. vitripennis* neuroblasts (*dpn*, white).

(G) HCR-FISH expression of *dmrt99B* (left, green), *tll* (left, magenta), *hth* (middle-left, magenta), *Bar-H* (middle-left, green), *opa* (middle, magenta), *slp* (middle, green), *hbn* (middle-right, green), *opa* (middle-right, magenta), *Bar-H* (right, green), and *tll* (right, magenta) in *T. castaneum* neuroblasts (*wor*, white).

Scale bar, 20 μm.

**Fig. S12. Experimental validation of new temporal transcription factors.**

(A) HCR-FISH shows *Ets65A* (magenta) expression in old *A. aegypti* neuroblasts (*dpn*, white).

(B) HCR-FISH shows *cas* (left, magenta) and *grh* (right, magenta) expression in mid and old *N. vitripennis* neuroblasts (*dpn*, white), respectively.

**Fig. S13. The absence of the *ey* temporal window leads to the absence of its neuronal progeny in *Musca*.**

UMAP projections showing the expression of *ey*, *erm*, *vvl*, *kn*, *TfAP-2*, and *svp* in developing visual brains of *M. domestica* (top) and *D. melanogaster* (bottom). In the *D. melanogaster ey* panel neuronal populations generated from different neuroblast temporal windows (Hth, Hth/Opa, Erm/Ey, and Ey/Hbn windows) are circled, and annotated based on known combinations of the plotted markers than can distinguish them from each other. In *M. domestica*, all corresponding neuronal populations are detected except those expressing *ey* (no *ey+ Lim3+ tup-* population), consistent with the absence of an Ey temporal window. NotchON neurons expressing *vvl* are markedly fewer in *M. domestica* than in *D. melanogaster* and co-express *erm*, indicating they derive from the Erm temporal window rather than the Ey window.

## Table Legends

**Table S1: One-to-one, one-to-many, many-to-one, many-to-many homologs, and species-specific genes for each insect**

**Table S2: Transcription factor lists for each insect species.**

**Table S3: Candidate temporal transcription factor lists for each insect species.**

**Table S4: HCR-FISH probe sets used for validation of candidate temporal transcription factors**

**Table S5: Insect strains, genomes, and reagents**

