## Supplementary figures and images for "Evolutionary dynamics of temporal transcription factor series in the insect visual brain"

### Figure S1

Supplementary Figure 1

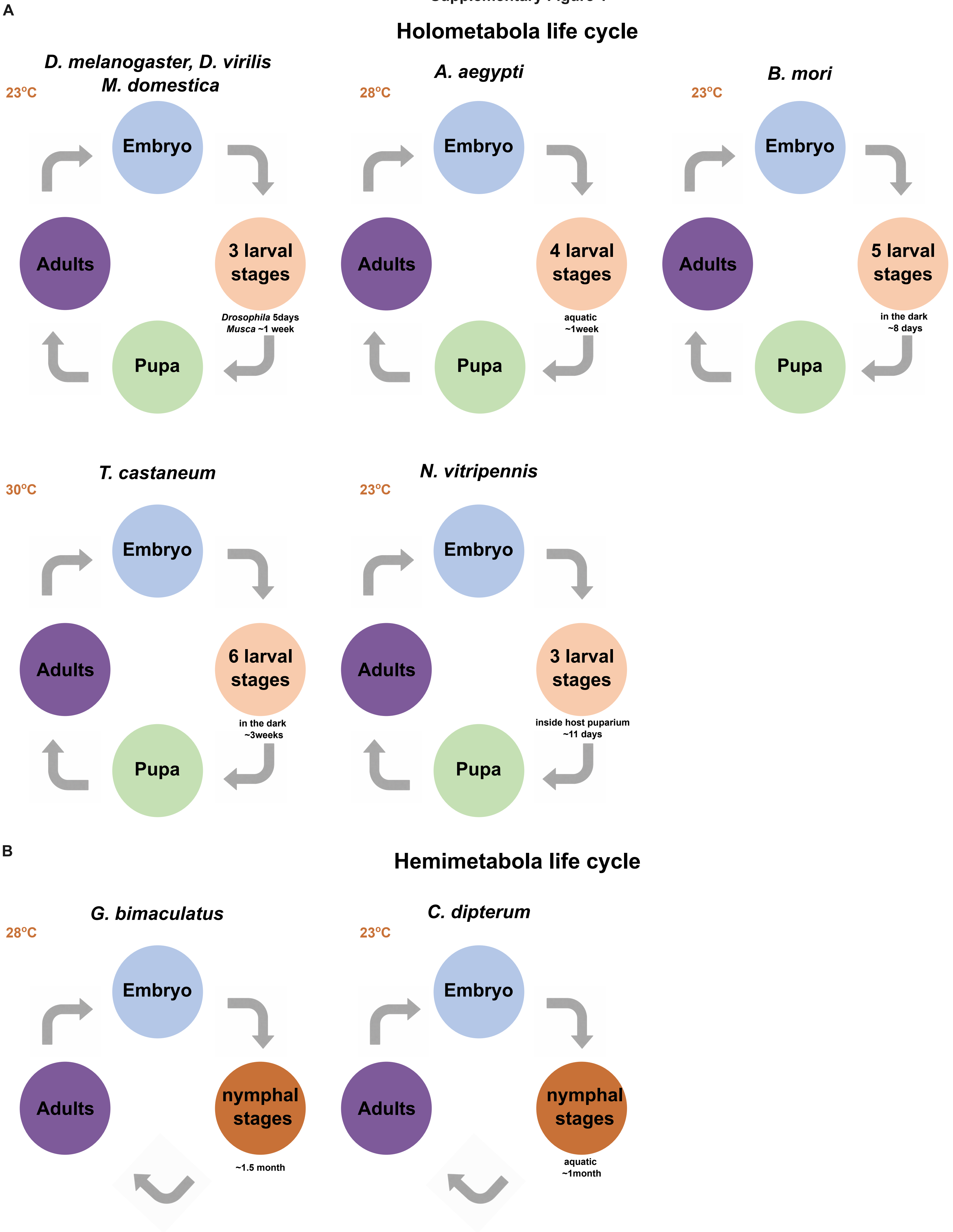

### Figure S2

##### Supplementary Figure 2

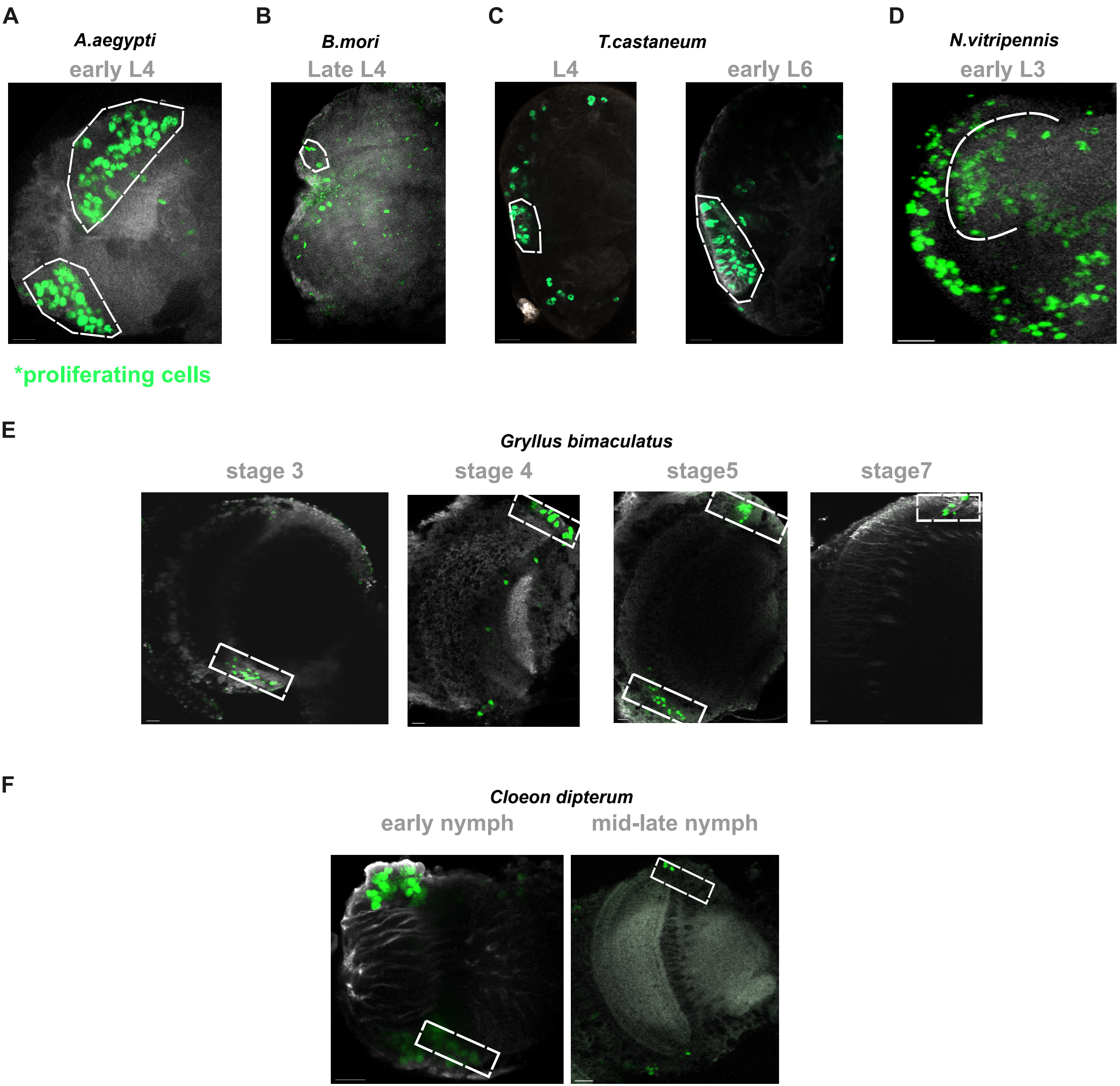

### Figure S3

Supplementary Figure 3

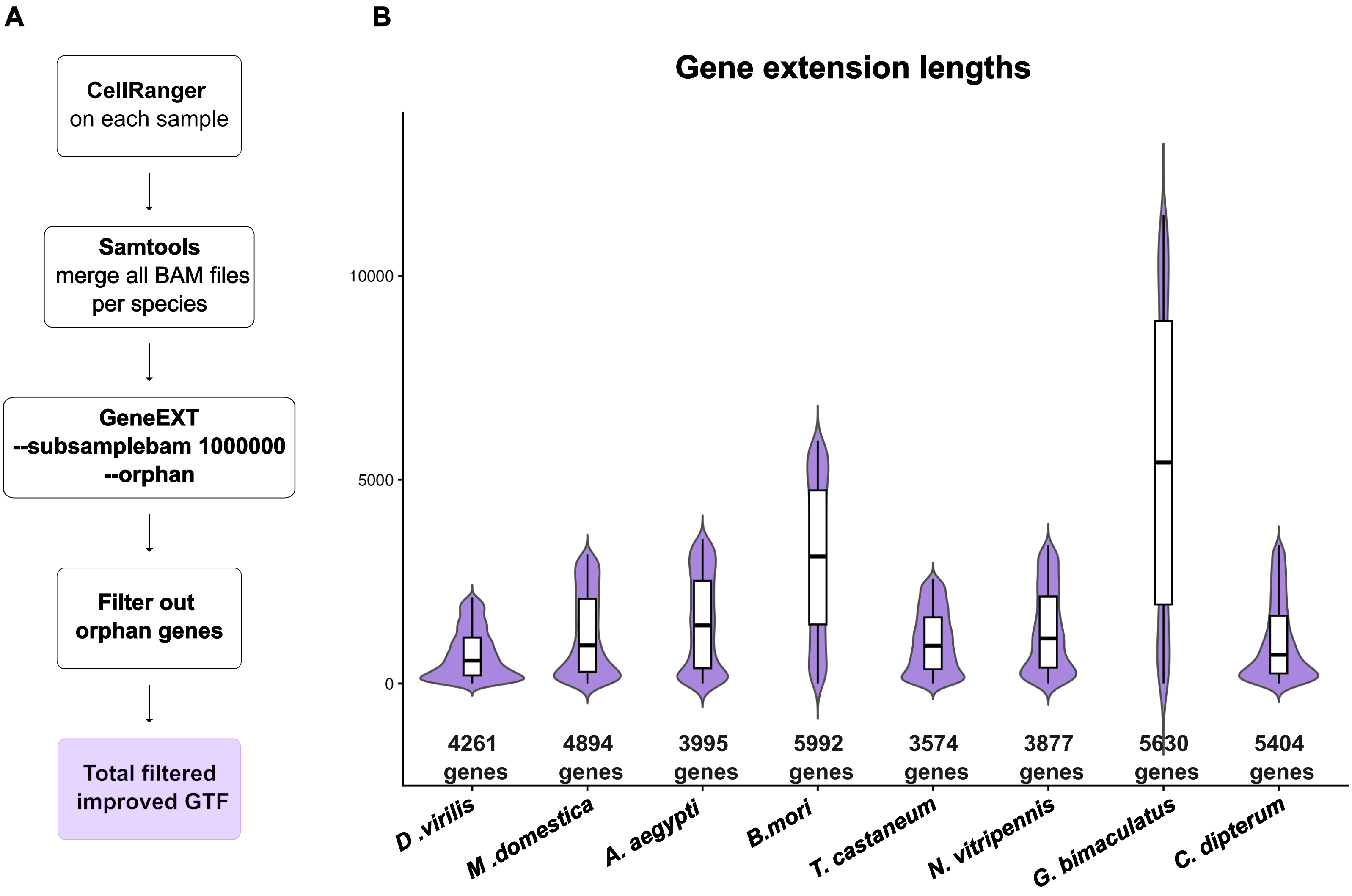

### Figure S5

#### Supplementary Figure 5

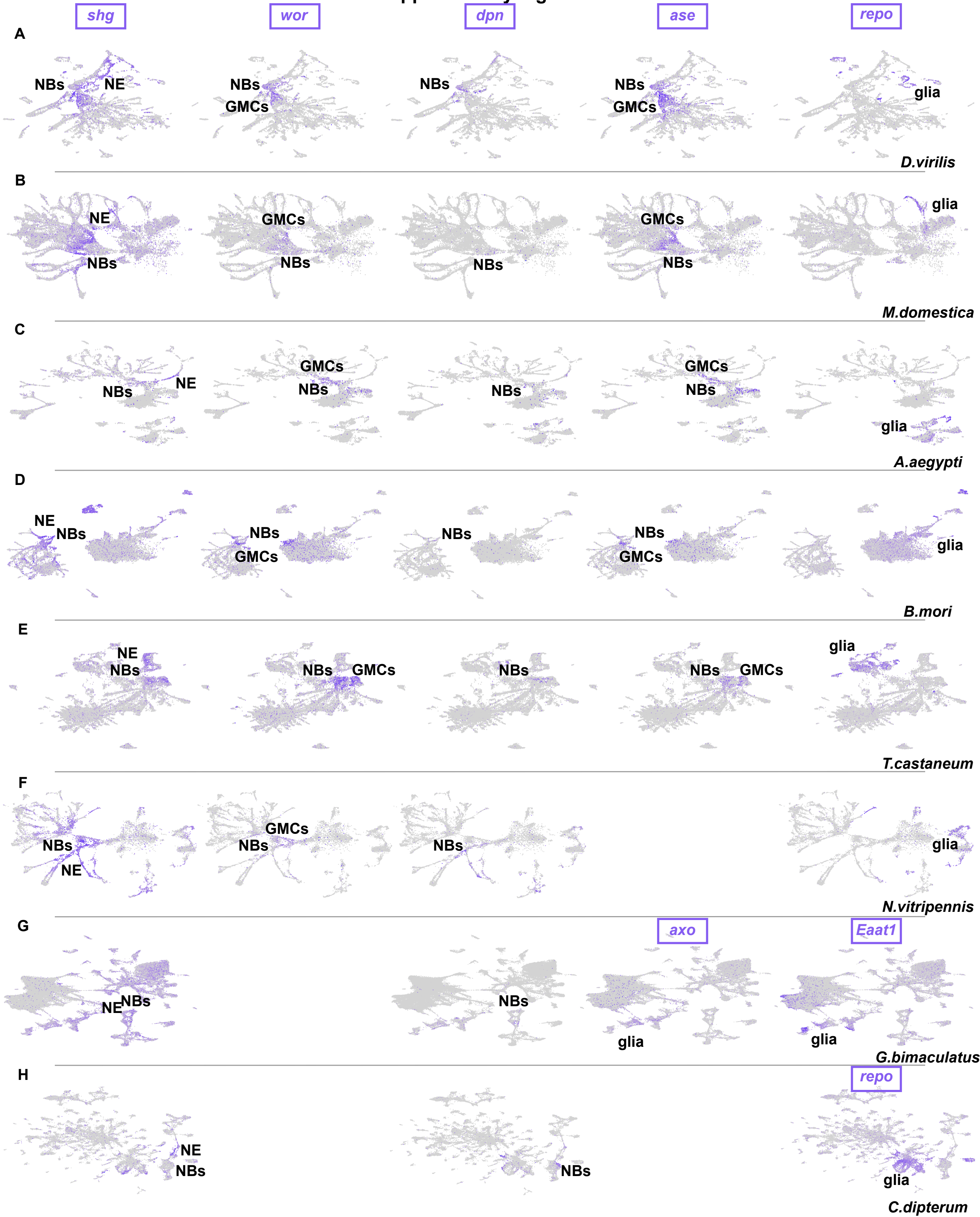

### Figure S6

#### Supplementary Figure 6

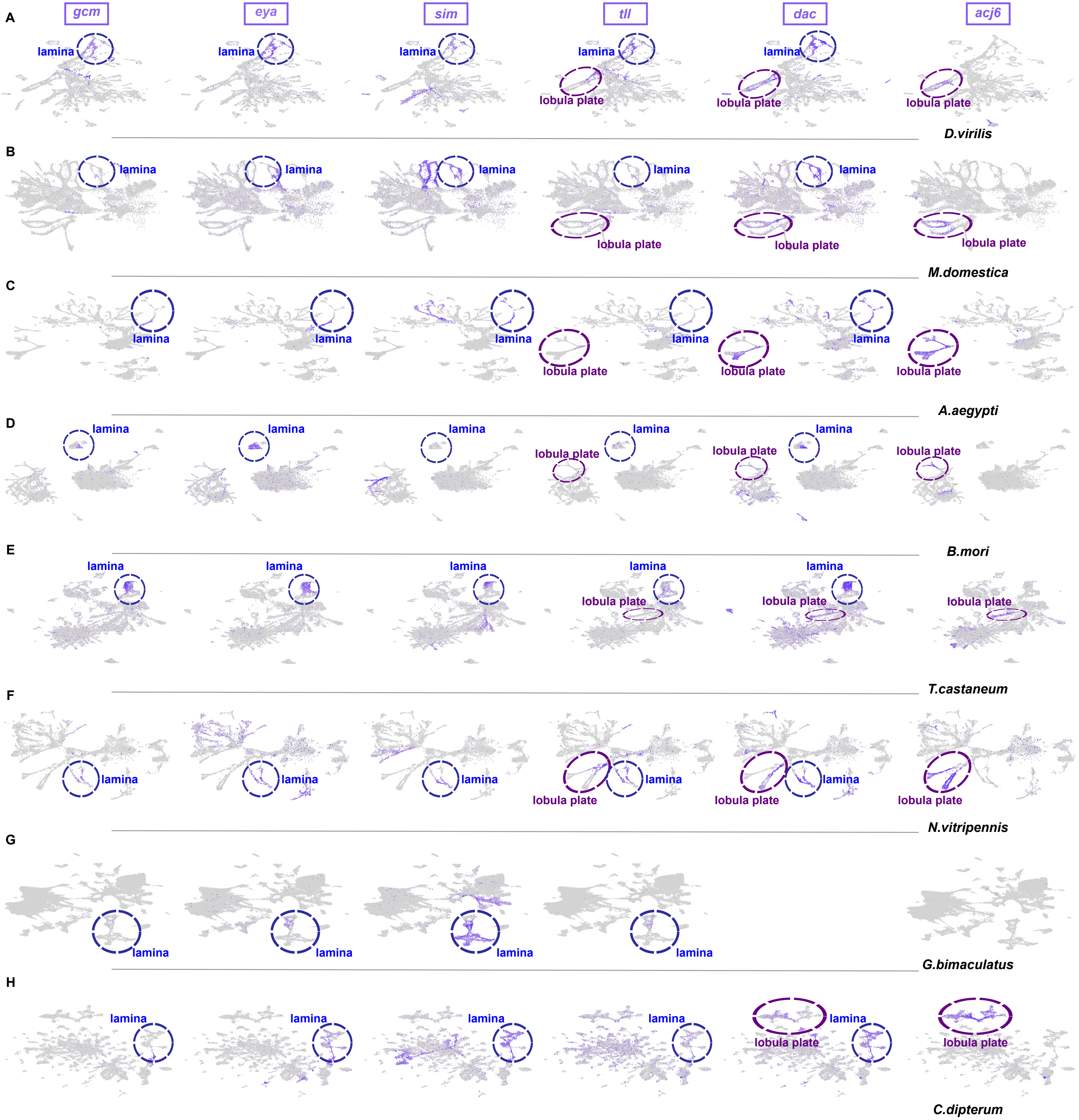

### Figure S7

Supplementary Figure 7

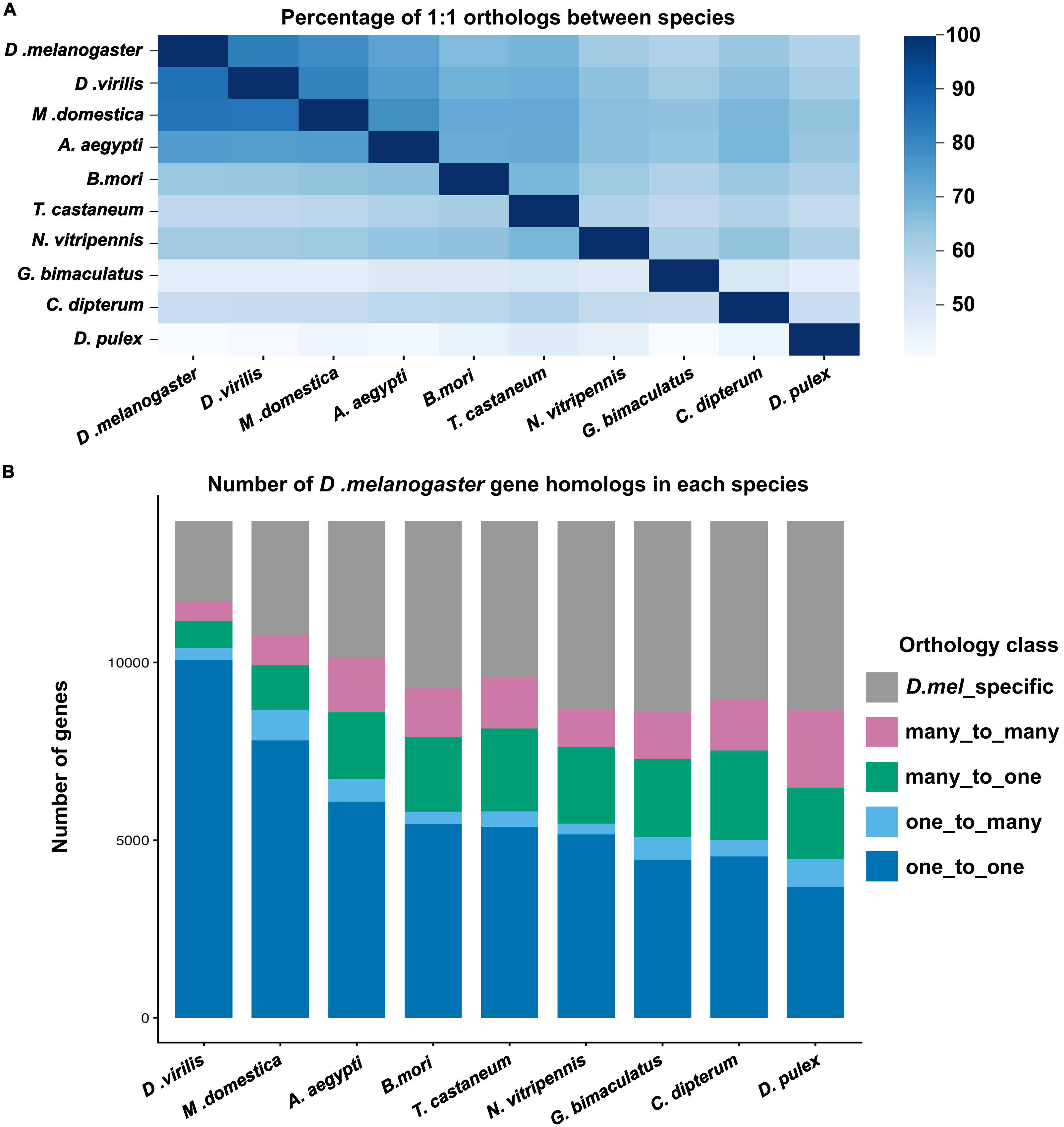

### Figure S8

##### Supplementary Figure 8

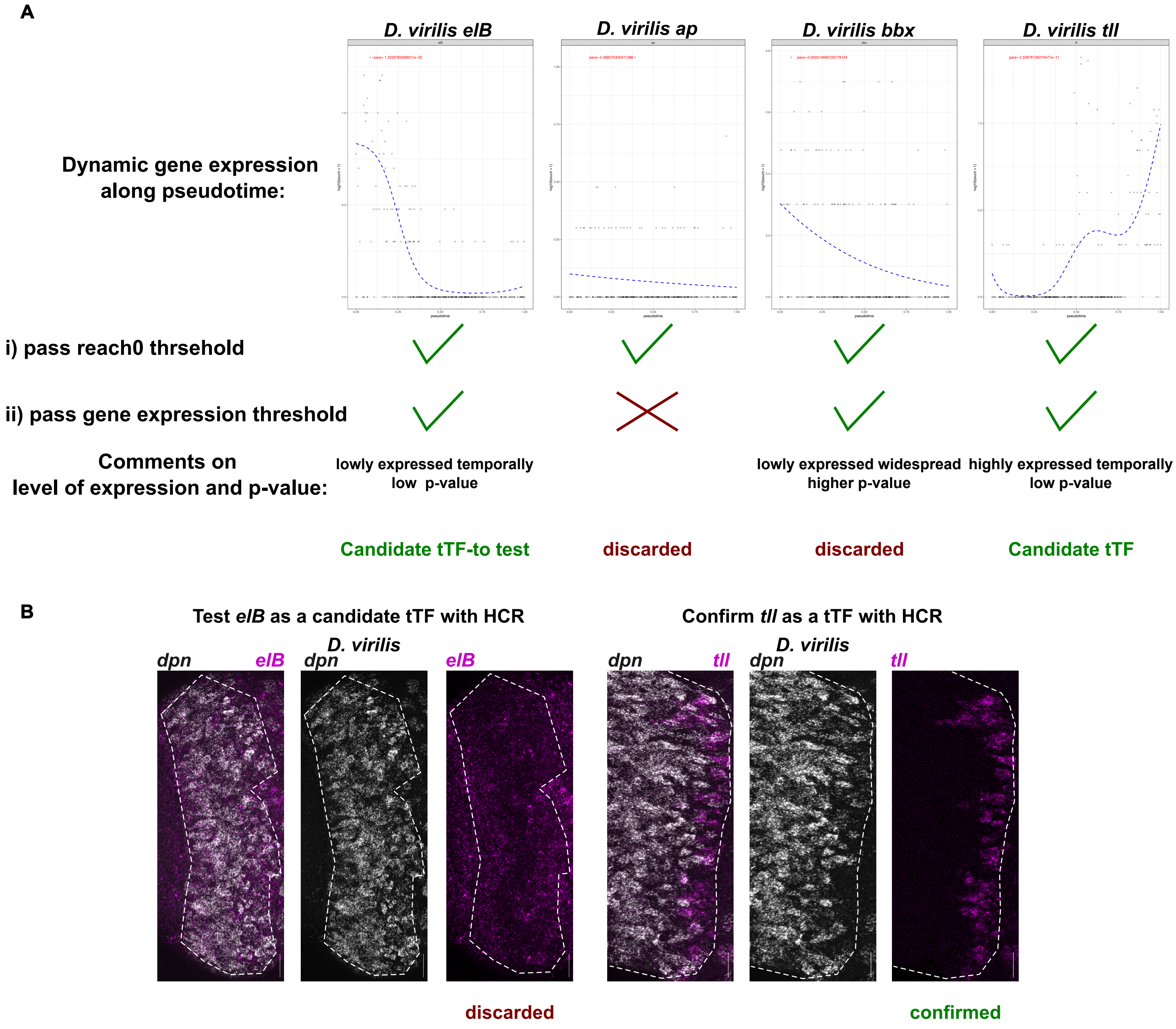

### Figure S10

Supplementary Figure 10

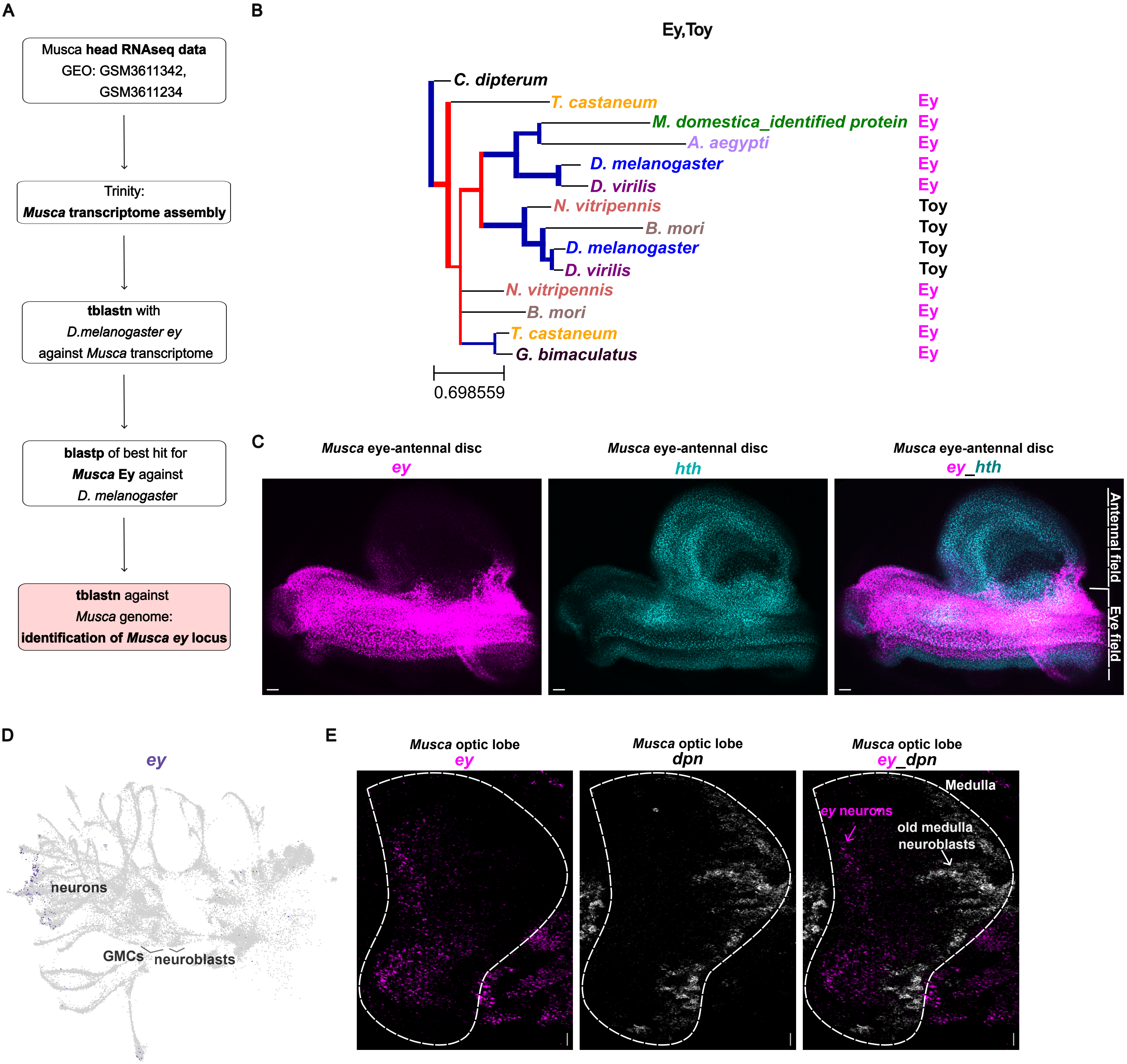

### Figure S11

#### Supplementary Figure 11

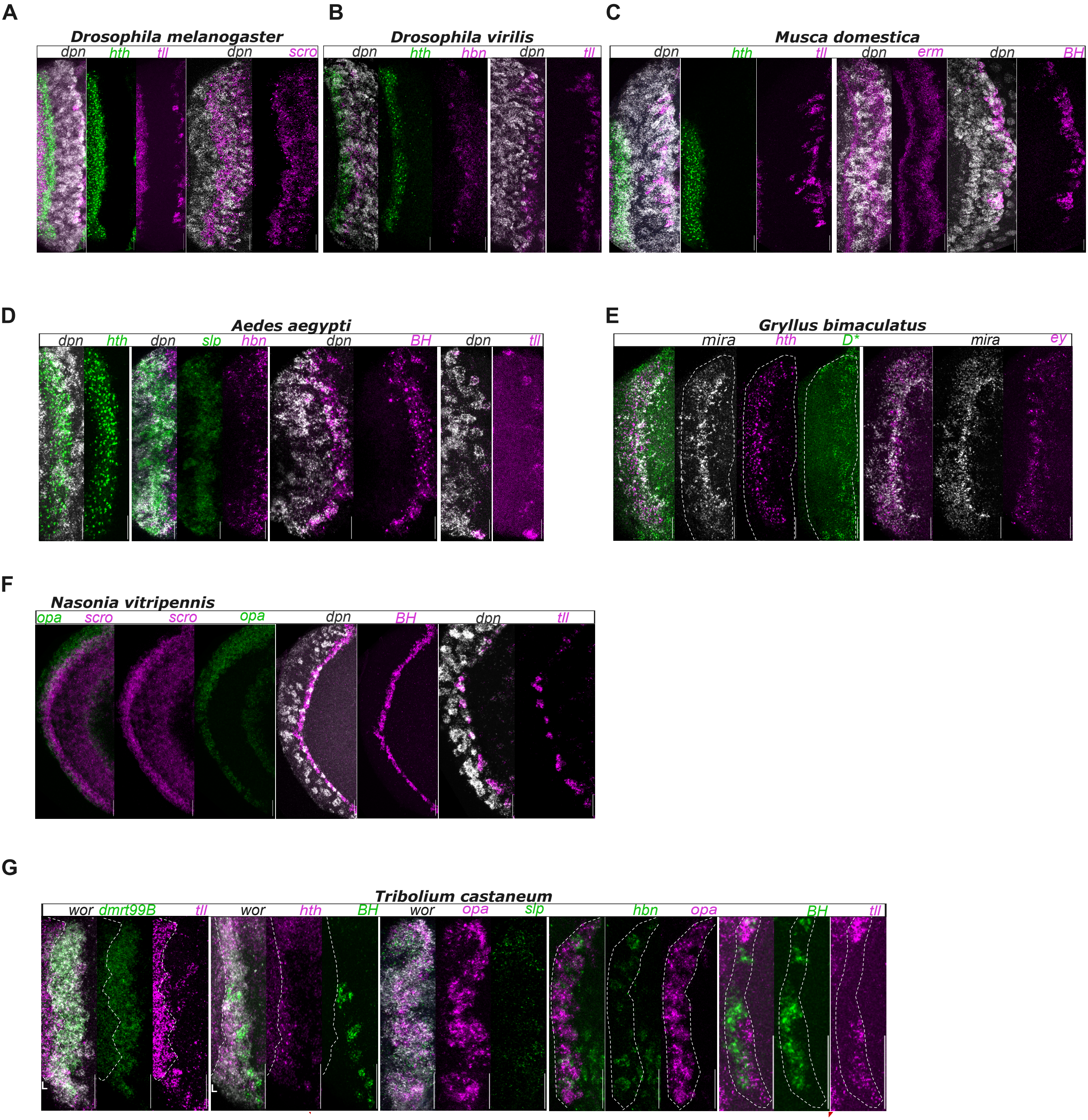

### Figure S12

#### Supplementary Figure 12

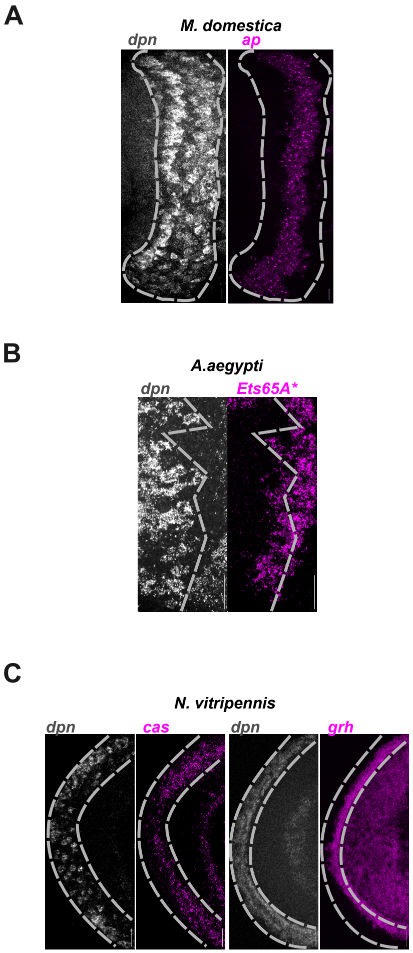

### Figure S13

Supplementary Figure 13

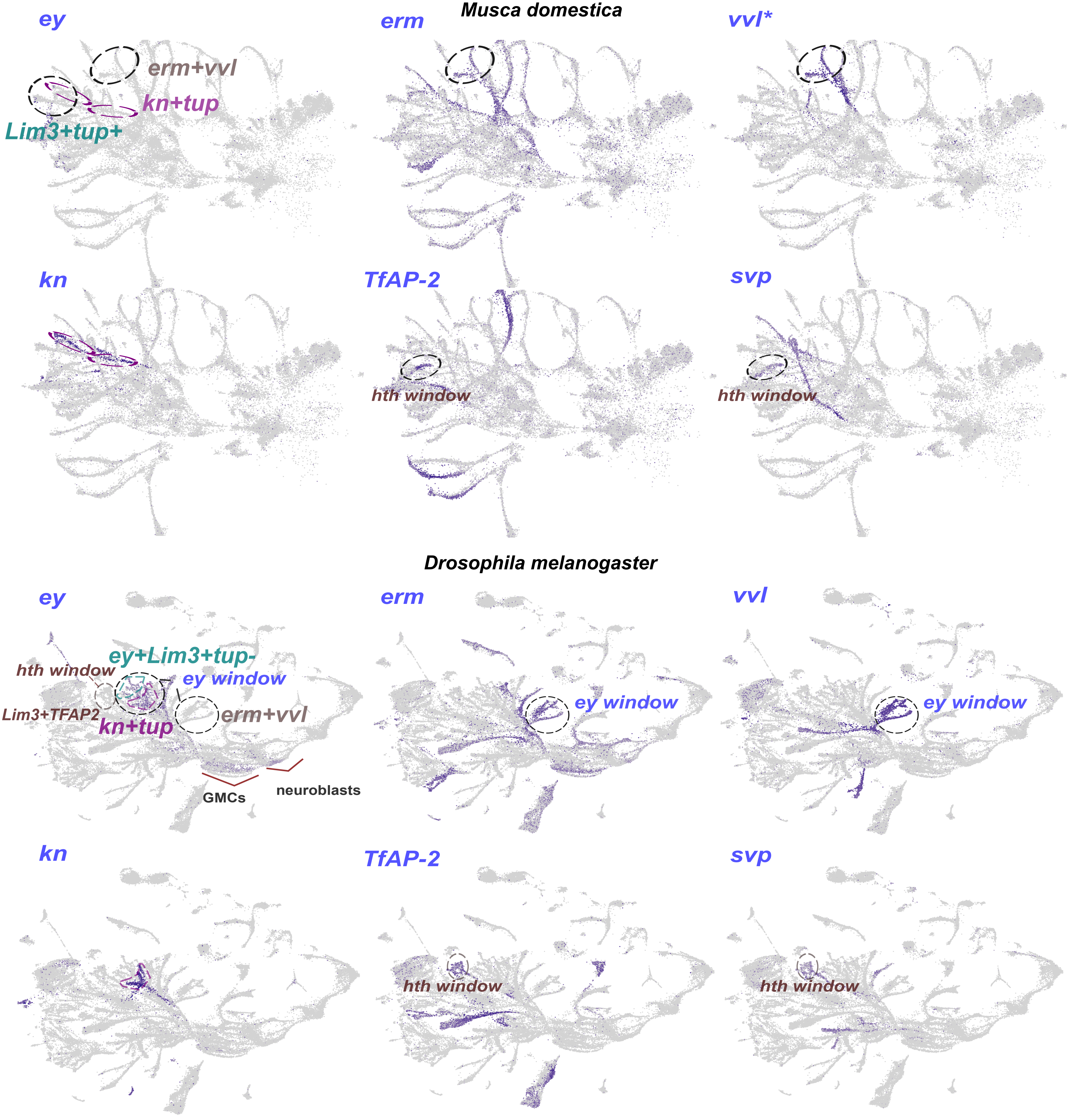
