## Supplementary material for "Evolutionary dynamics of temporal transcription factor series in the insect visual brain": Figure S4

### Supplementary Figure 4

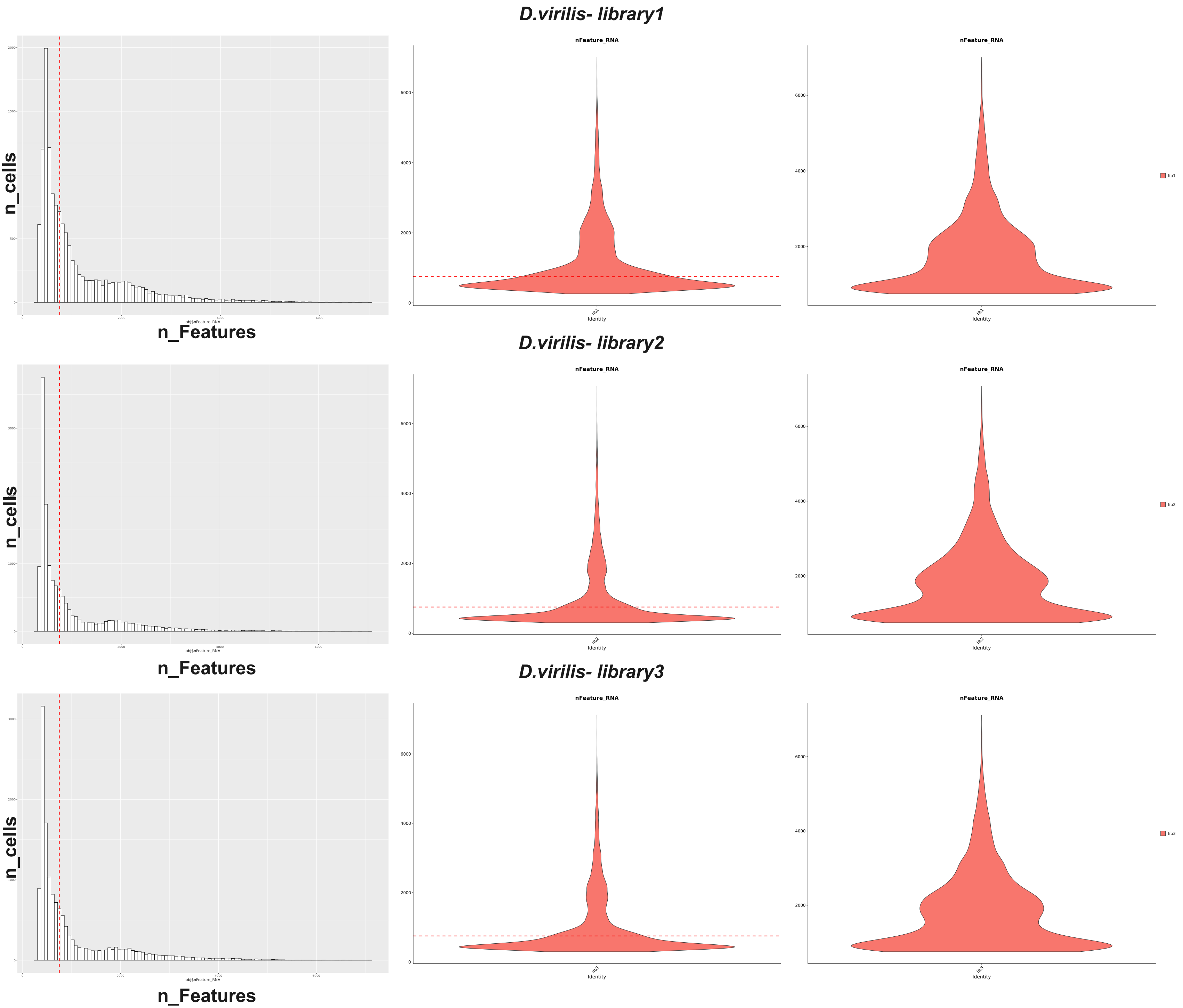

#### *M.domestica - library1*

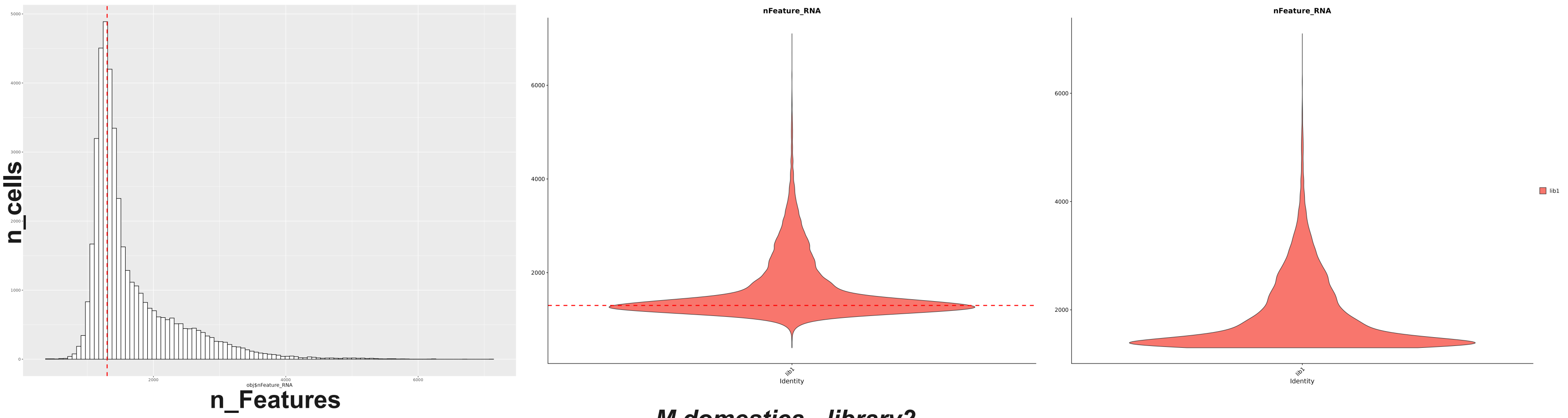


#### *M.domestica - library2*

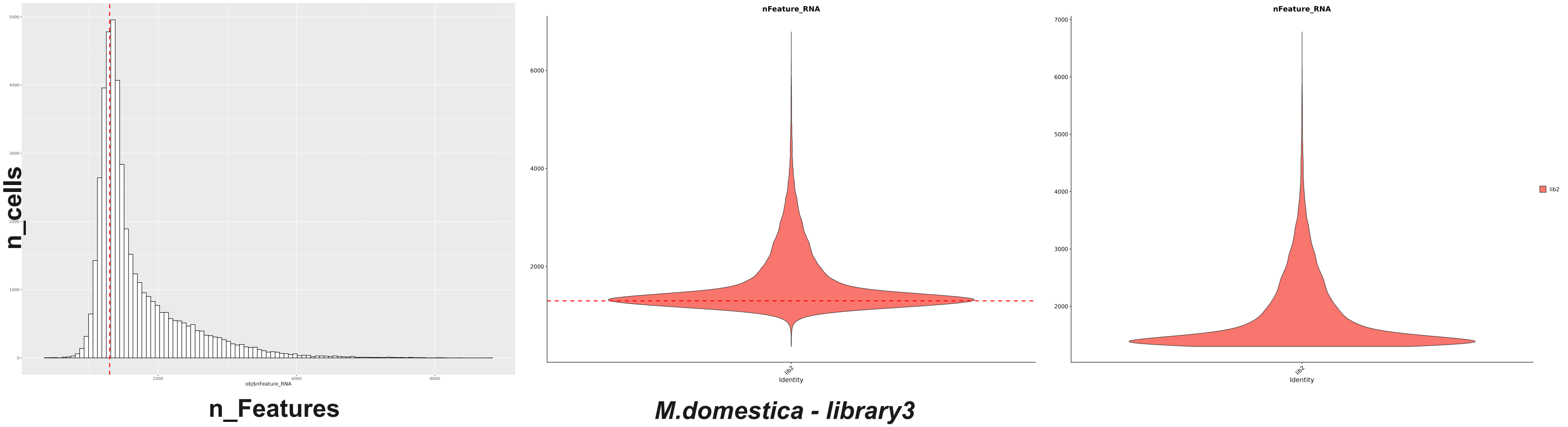


#### *M.domestica - library3*

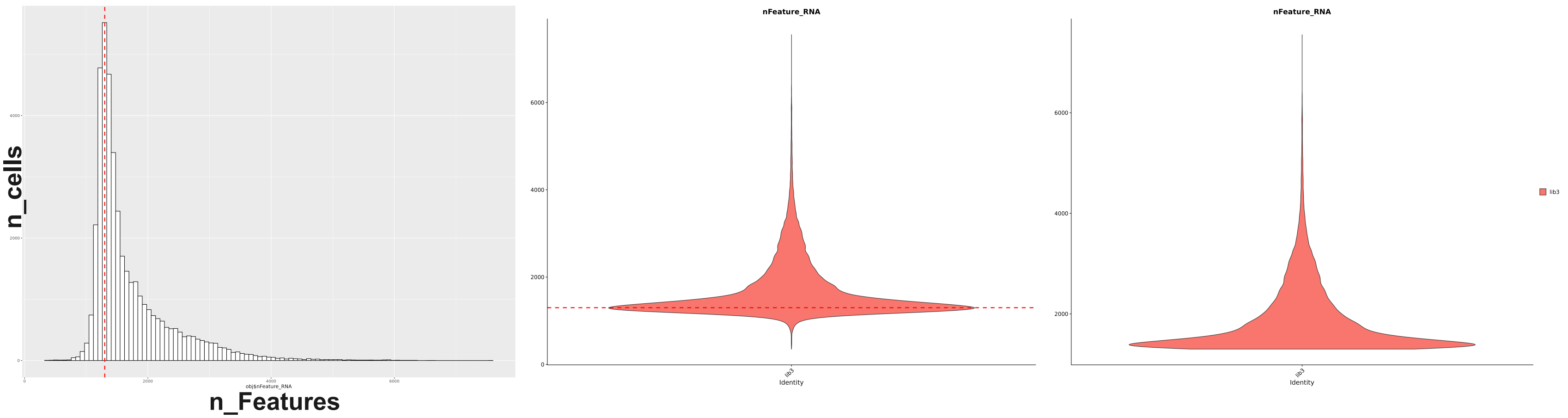

#### *A. aegypti - library1*

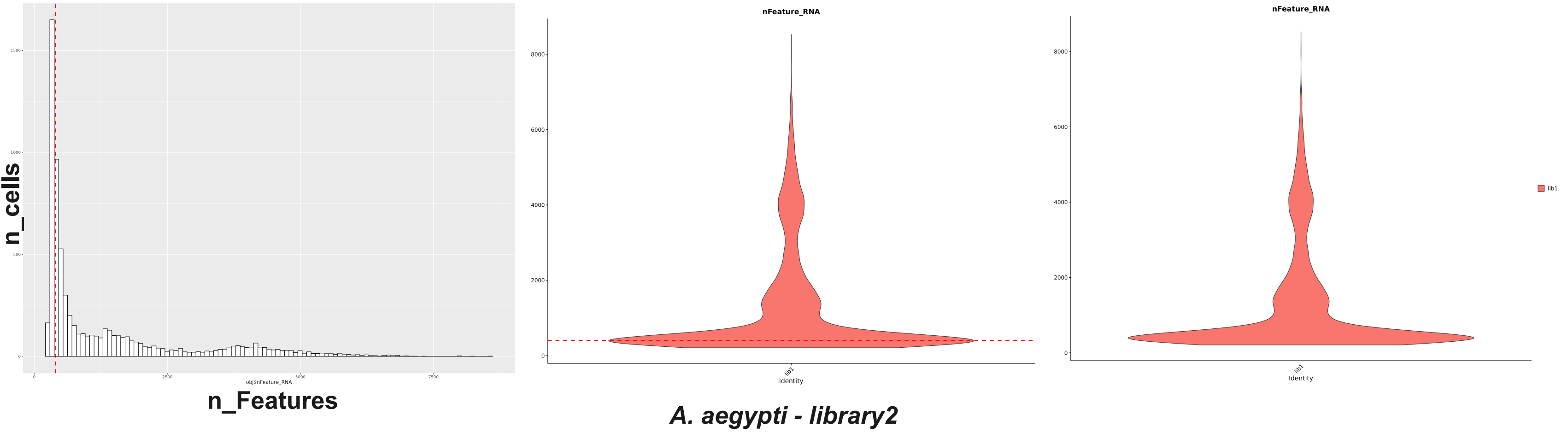


#### *A. aegypti - library2*

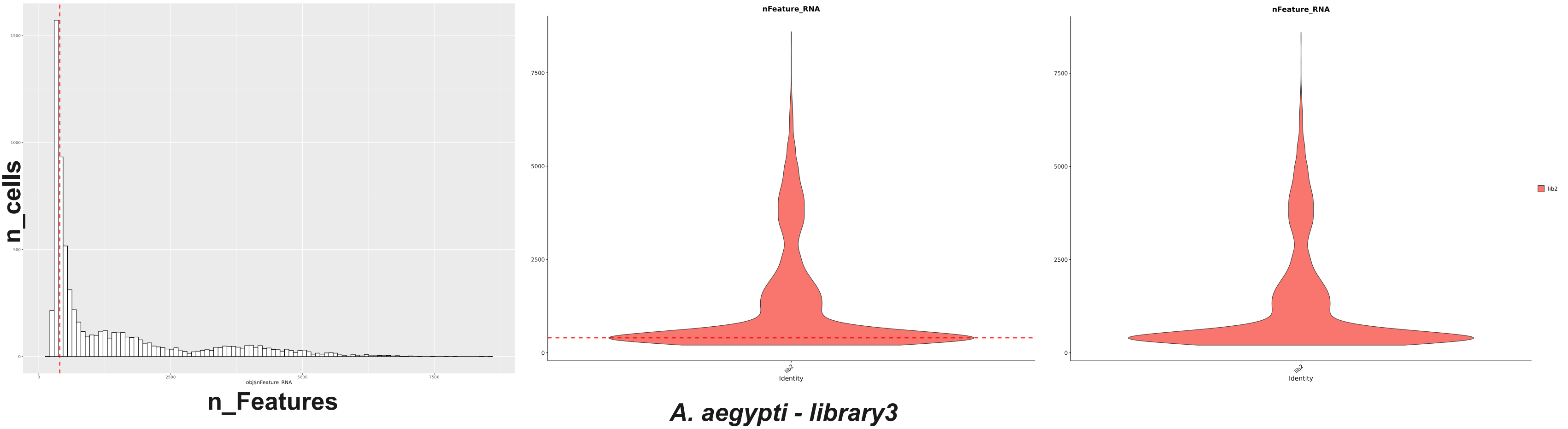


#### *A. aegypti - library3*

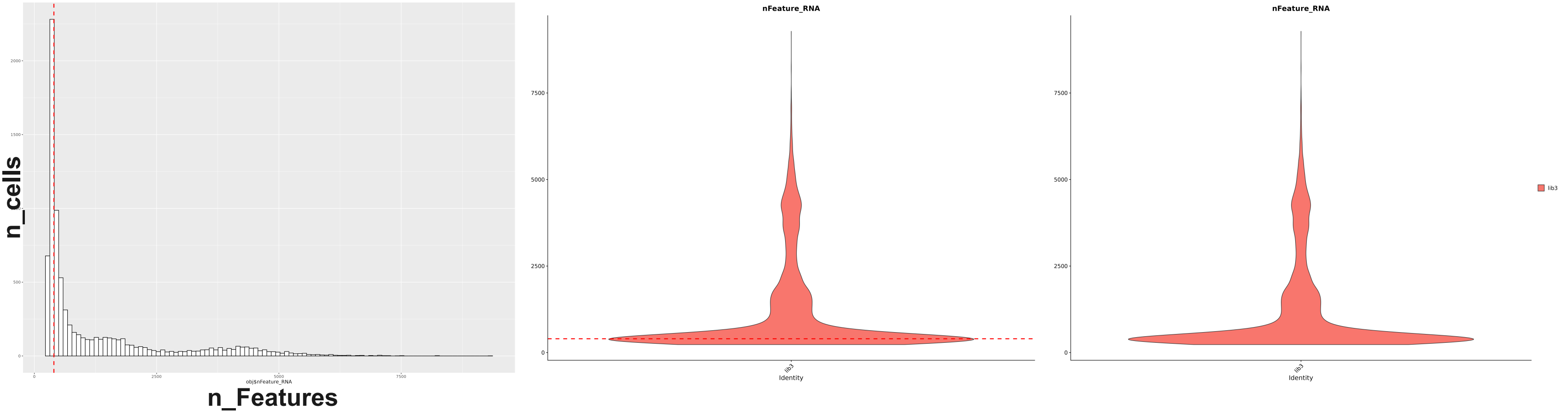

#### B. mori - library1

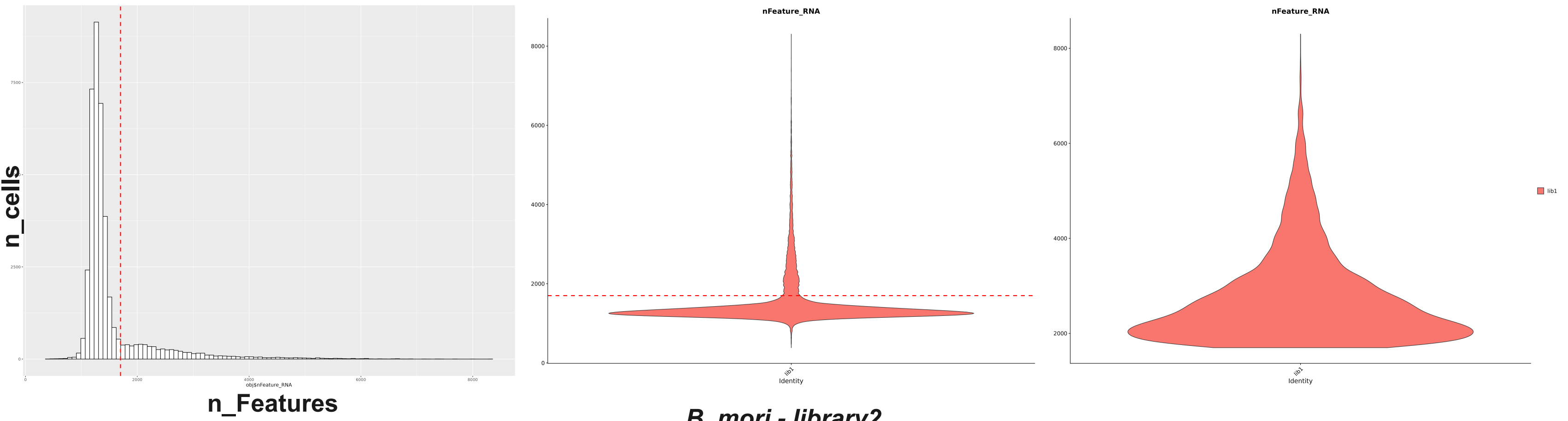


#### B. mori - library2

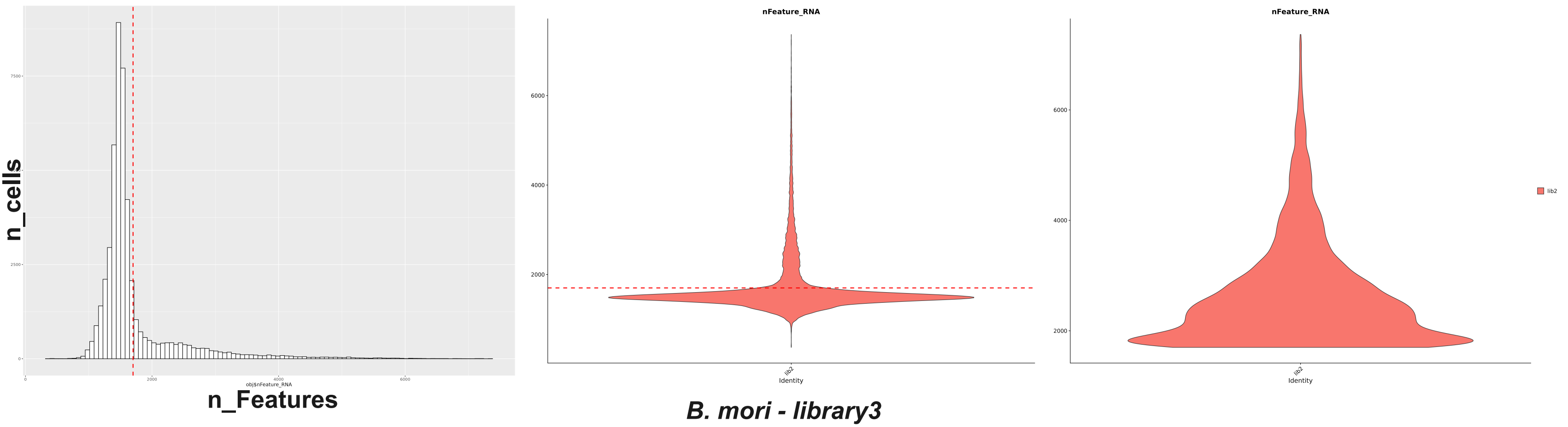


#### B. mori - library3

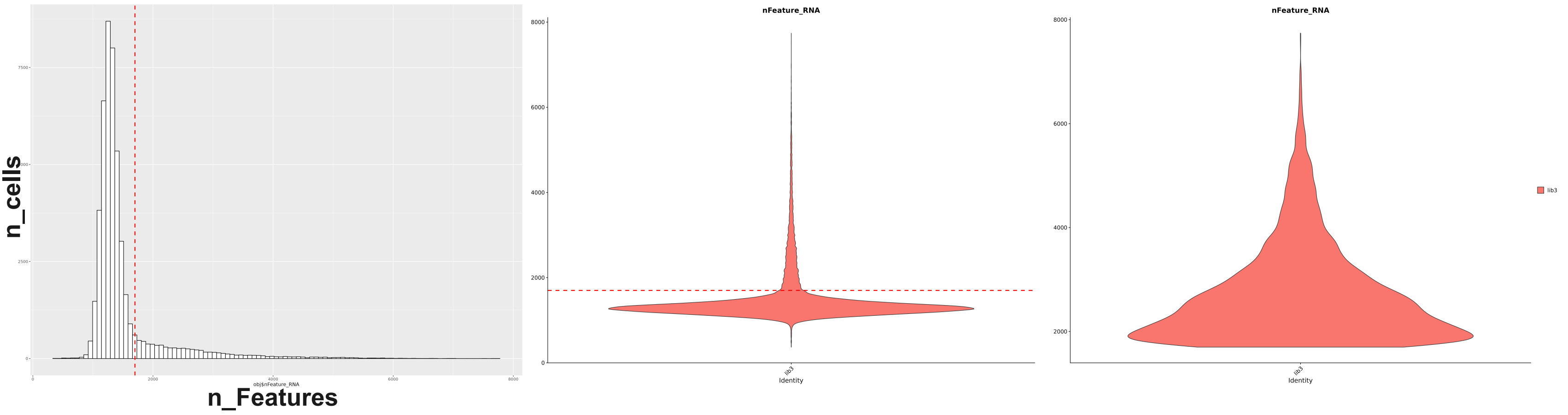

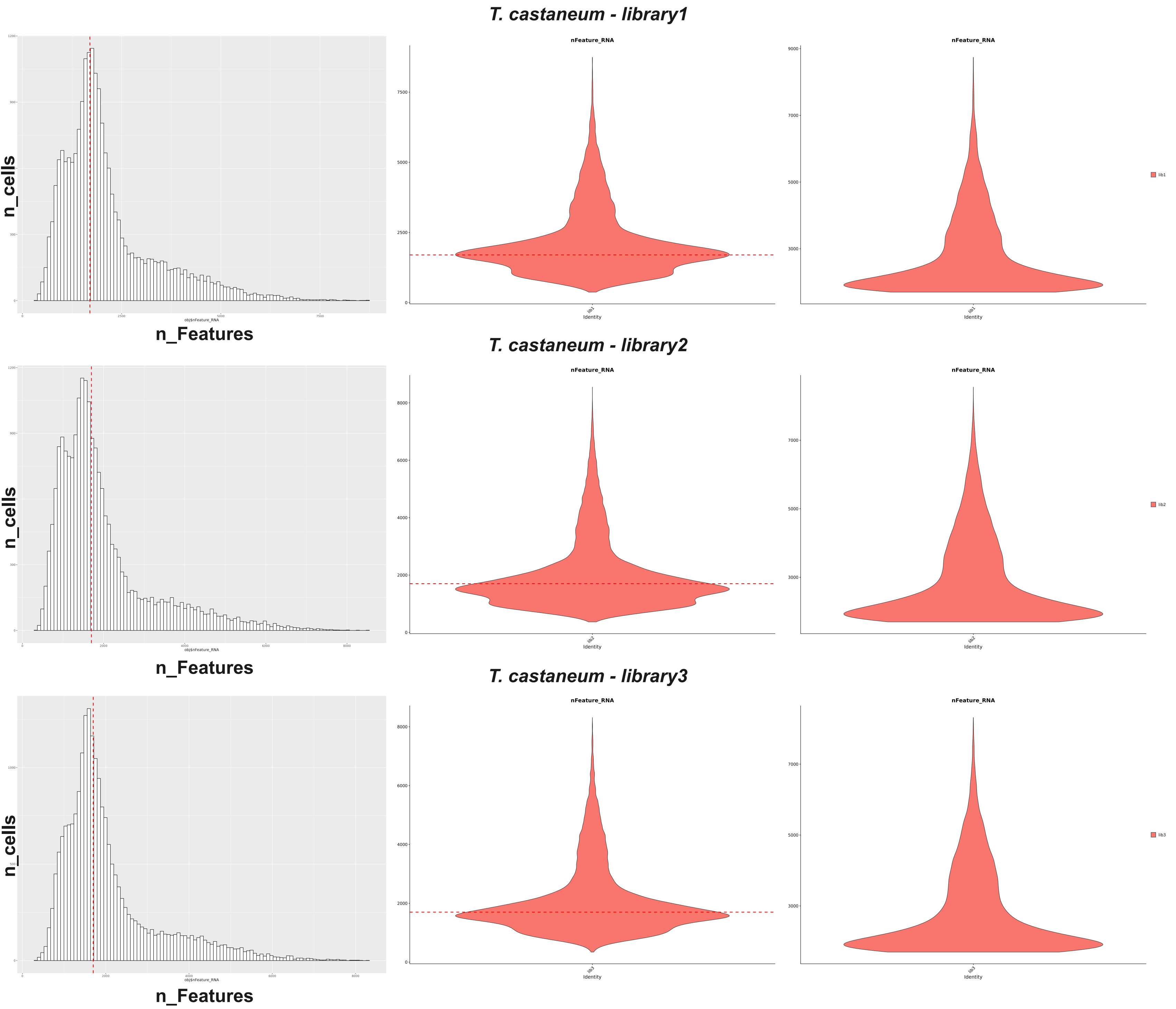
T. castaneum - library1
n_cells
n_Features
obj$nFeature_RNA
nFeature_RNA
Identity
lib1
T. castaneum - library2
n_cells
n_Features
obj$nFeature_RNA
nFeature_RNA
Identity
lib2
T. castaneum - library3
n_cells
n_Features
obj$nFeature_RNA
nFeature_RNA
Identity
lib3

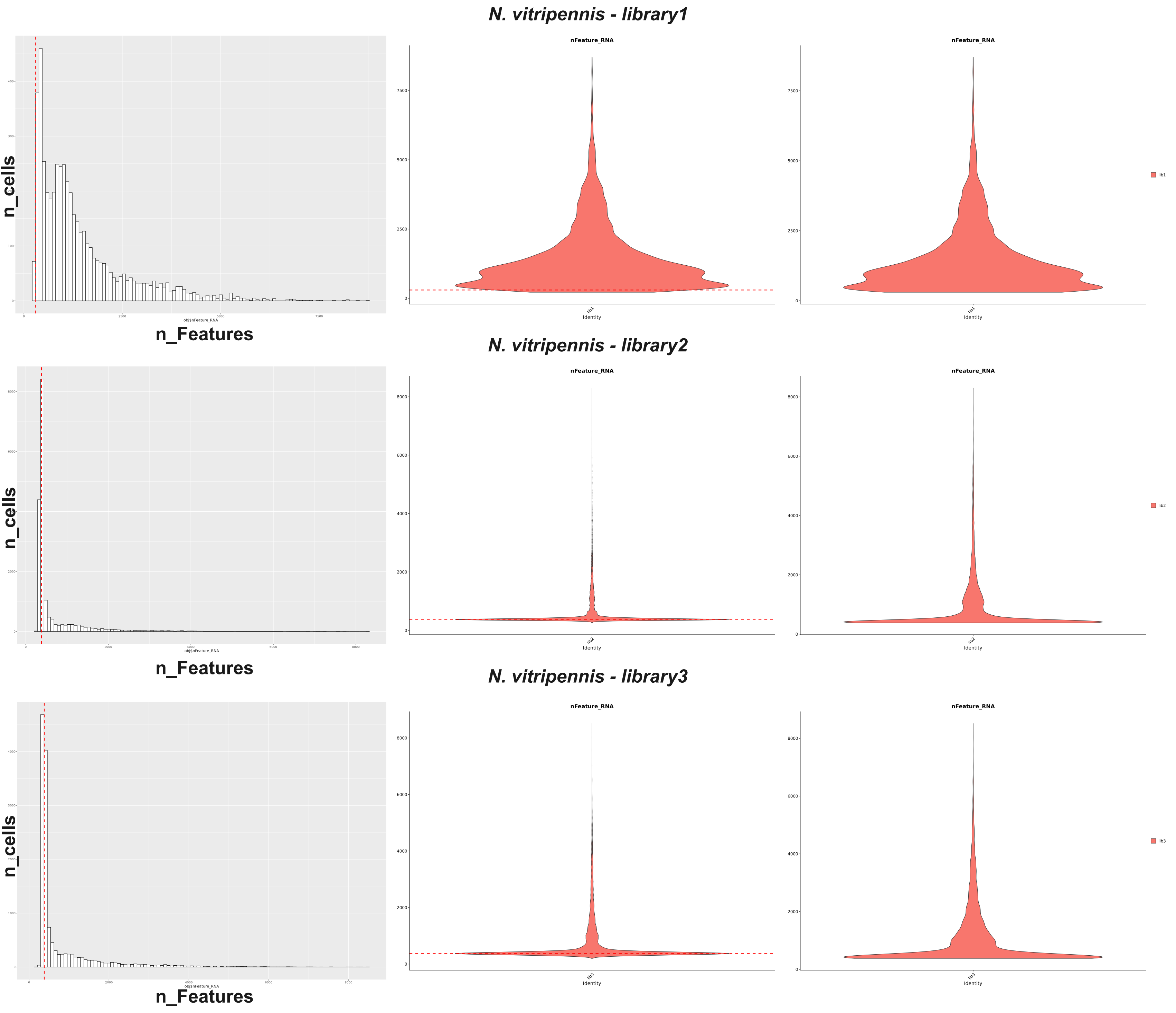

N. vitripennis - library1
nFeature_RNA
nFeature_RNA
n_cells
n_Features
obj$nFeature_RNA
Identity
lib1
N. vitripennis - library2
nFeature_RNA
nFeature_RNA
n_cells
n_Features
obj$nFeature_RNA
Identity
lib2
N. vitripennis - library3
nFeature_RNA
nFeature_RNA
n_cells
n_Features
obj$nFeature_RNA
Identity
lib3

#### *C. dipterum - library1*

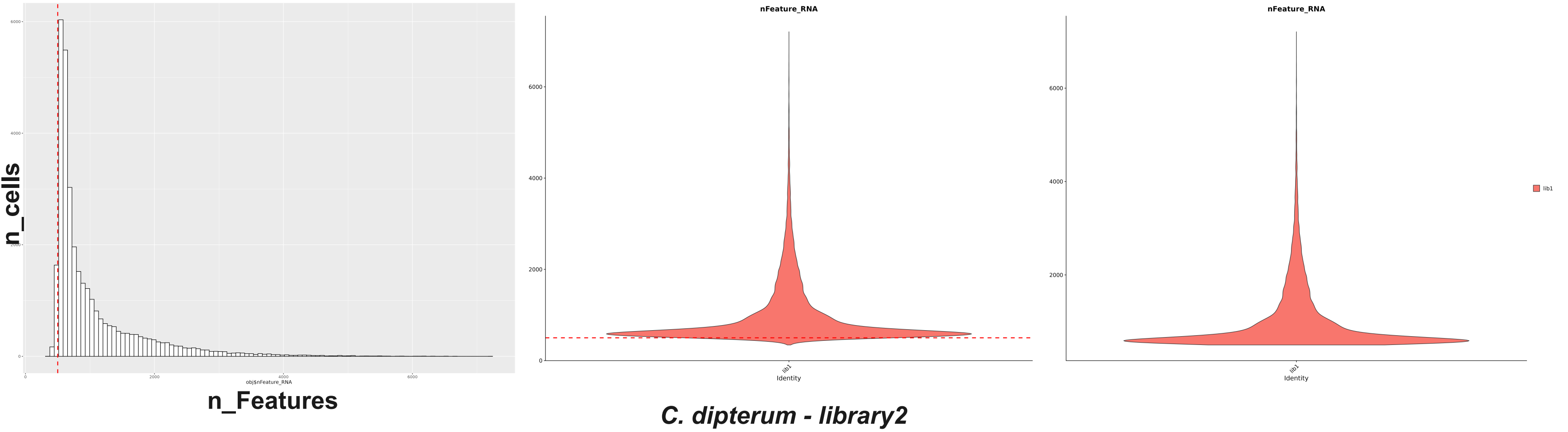


#### *C. dipterum - library2*

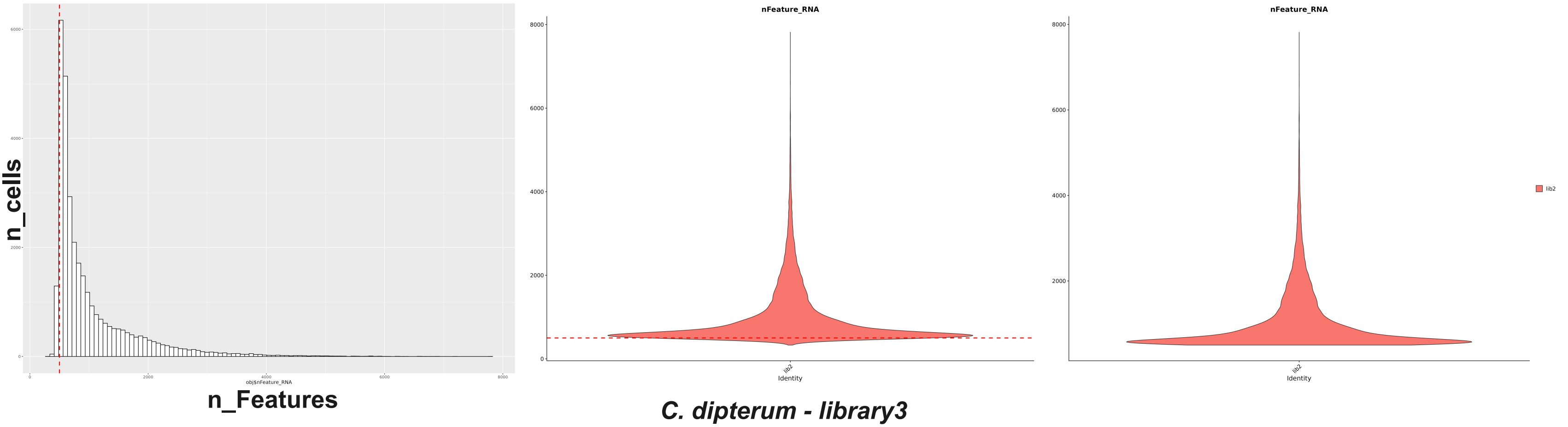


#### *C. dipterum - library3*

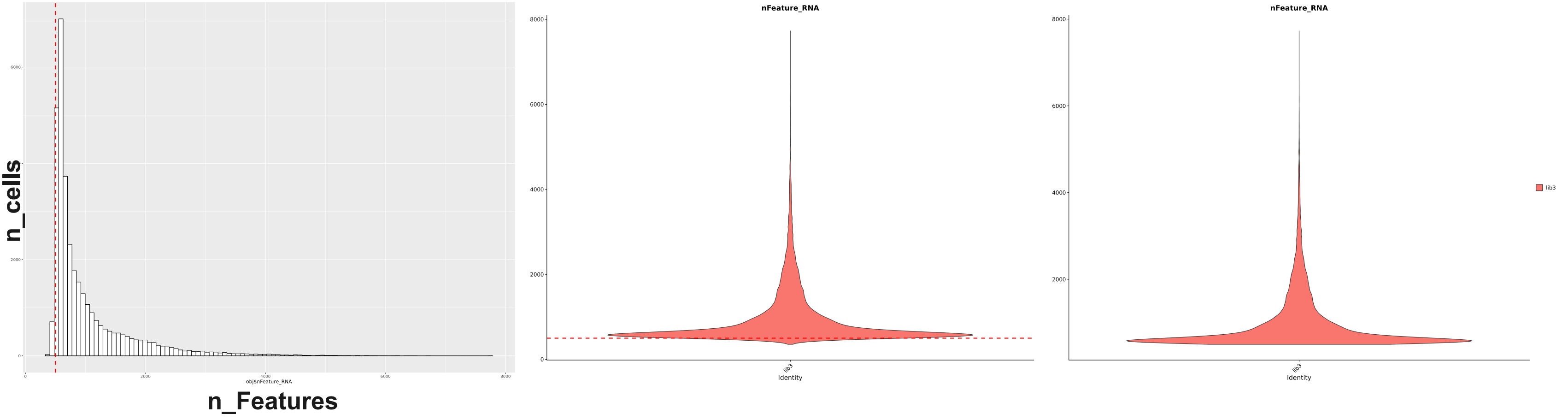

#### *G. bimaculatus stage1 - library1*

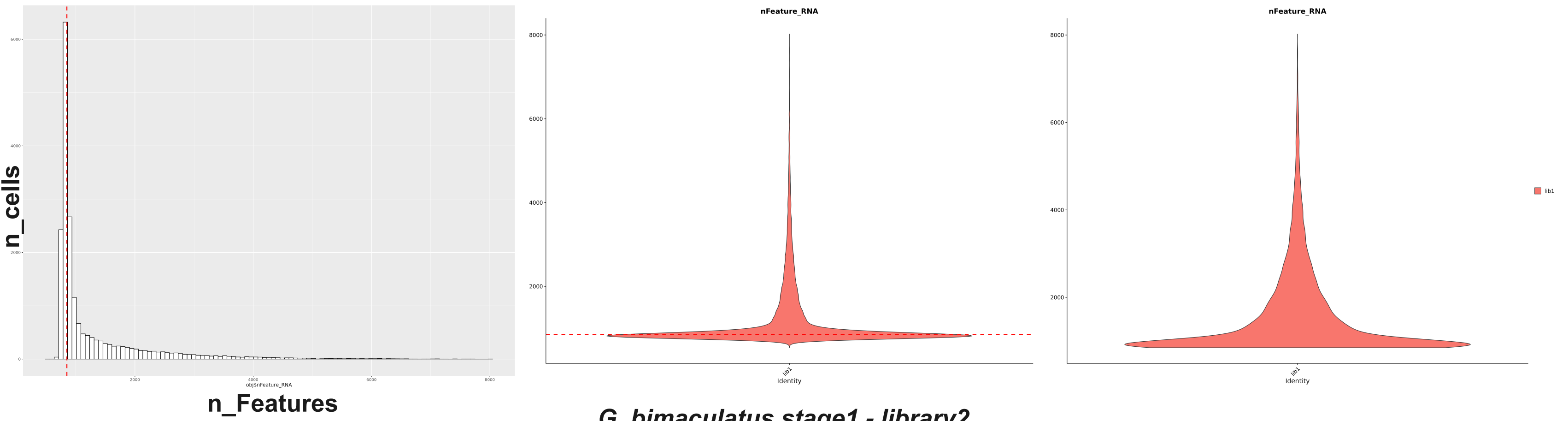


#### *G. bimaculatus stage1 - library2*

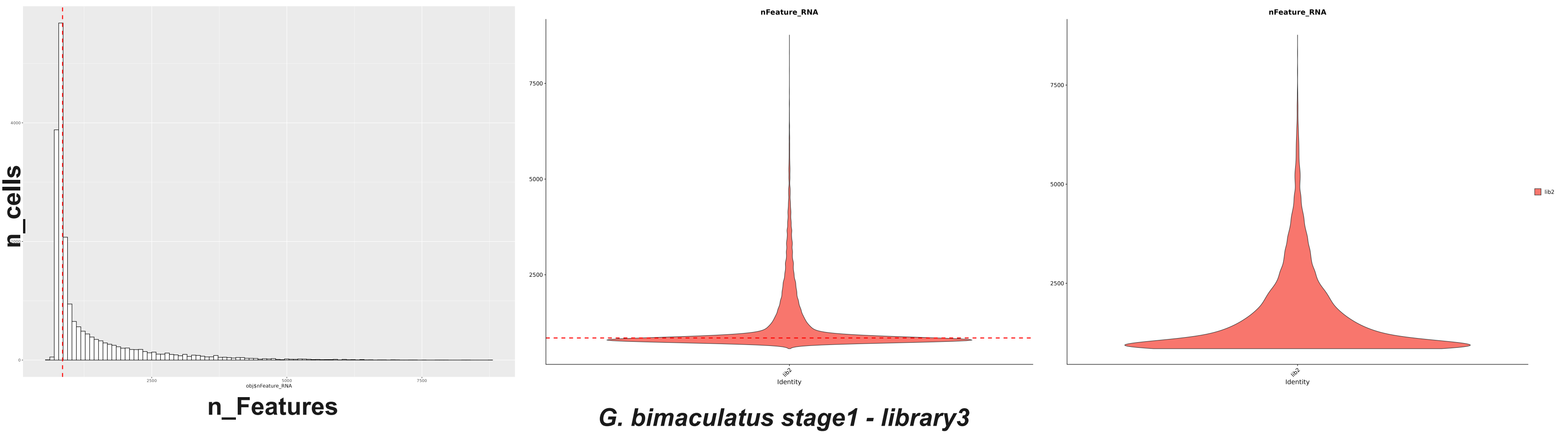


#### *G. bimaculatus stage1 - library3*

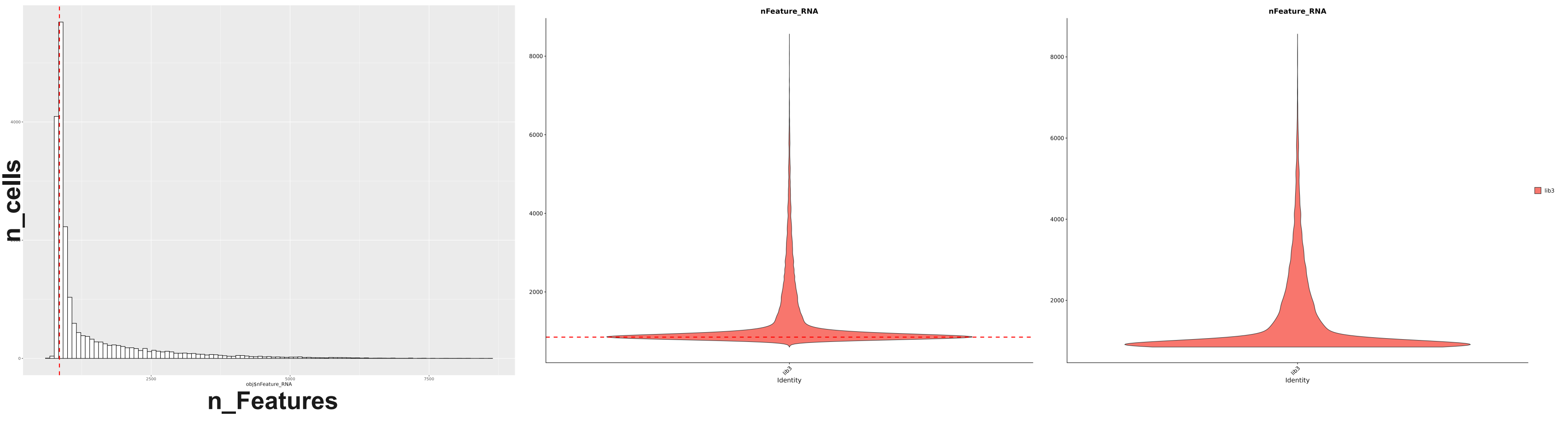

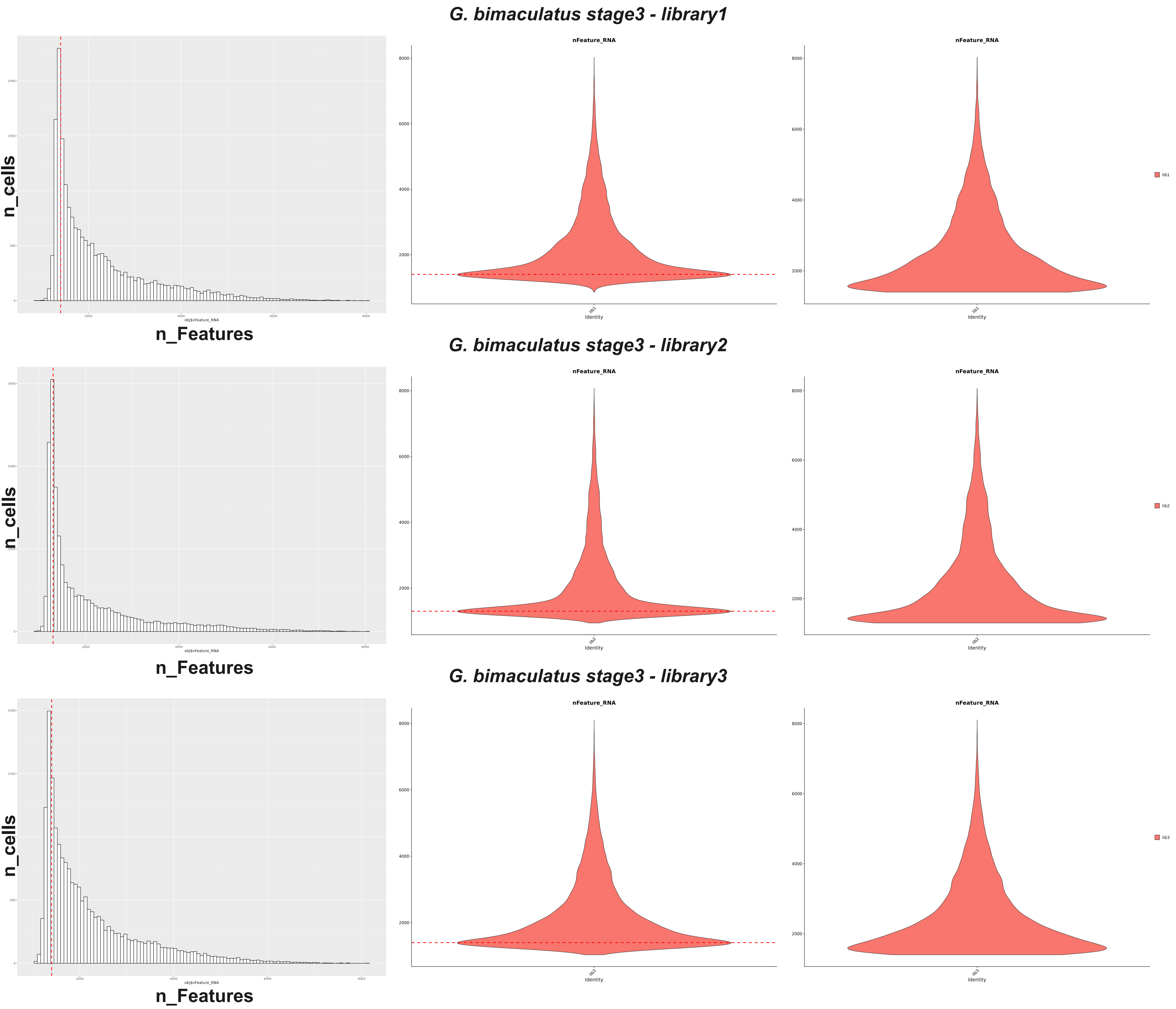
G. bimaculatus stage3 - library1
n_cells
n_Features
obj$nFeature_RNA
nFeature_RNA
8000
6000
4000
2000
lib1
Identity
G. bimaculatus stage3 - library2
n_cells
n_Features
obj$nFeature_RNA
nFeature_RNA
lib2
Identity
G. bimaculatus stage3 - library3
n_cells
n_Features
obj$nFeature_RNA
nFeature_RNA
lib3
Identity

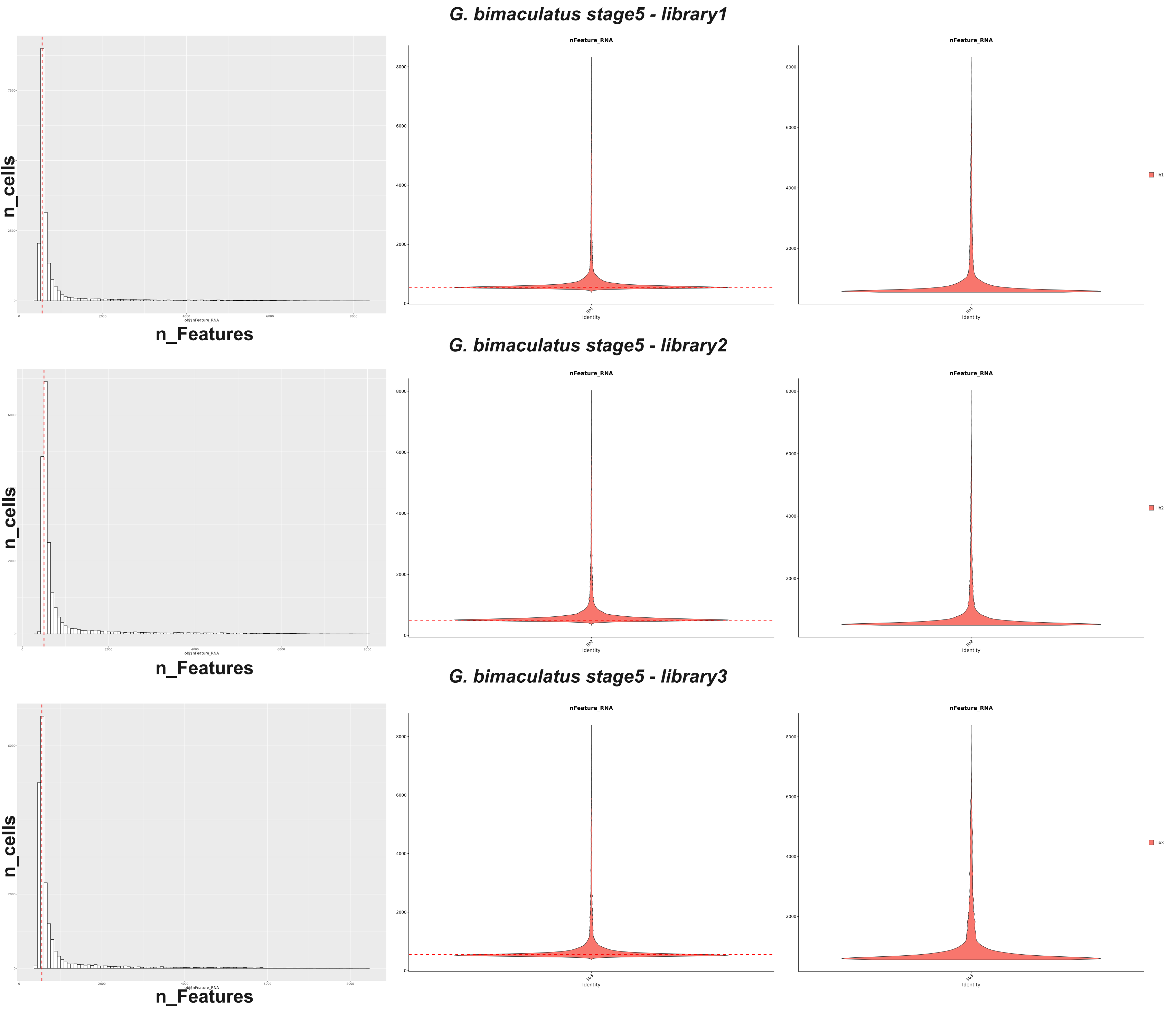
G. bimaculatus stage5 - library1
n_cells
n_Features
obj$nFeature_RNA
nFeature_RNA
Identity
lib1
G. bimaculatus stage5 - library2
n_cells
n_Features
obj$nFeature_RNA
nFeature_RNA
Identity
lib2
G. bimaculatus stage5 - library3
n_cells
n_Features
obj$nFeature_RNA
nFeature_RNA
Identity
lib3
