## Supplementary material for "Evolutionary dynamics of temporal transcription factor series in the insect visual brain": Figure S9

### Supplementary Figure 9

#### *D.melanogaster*

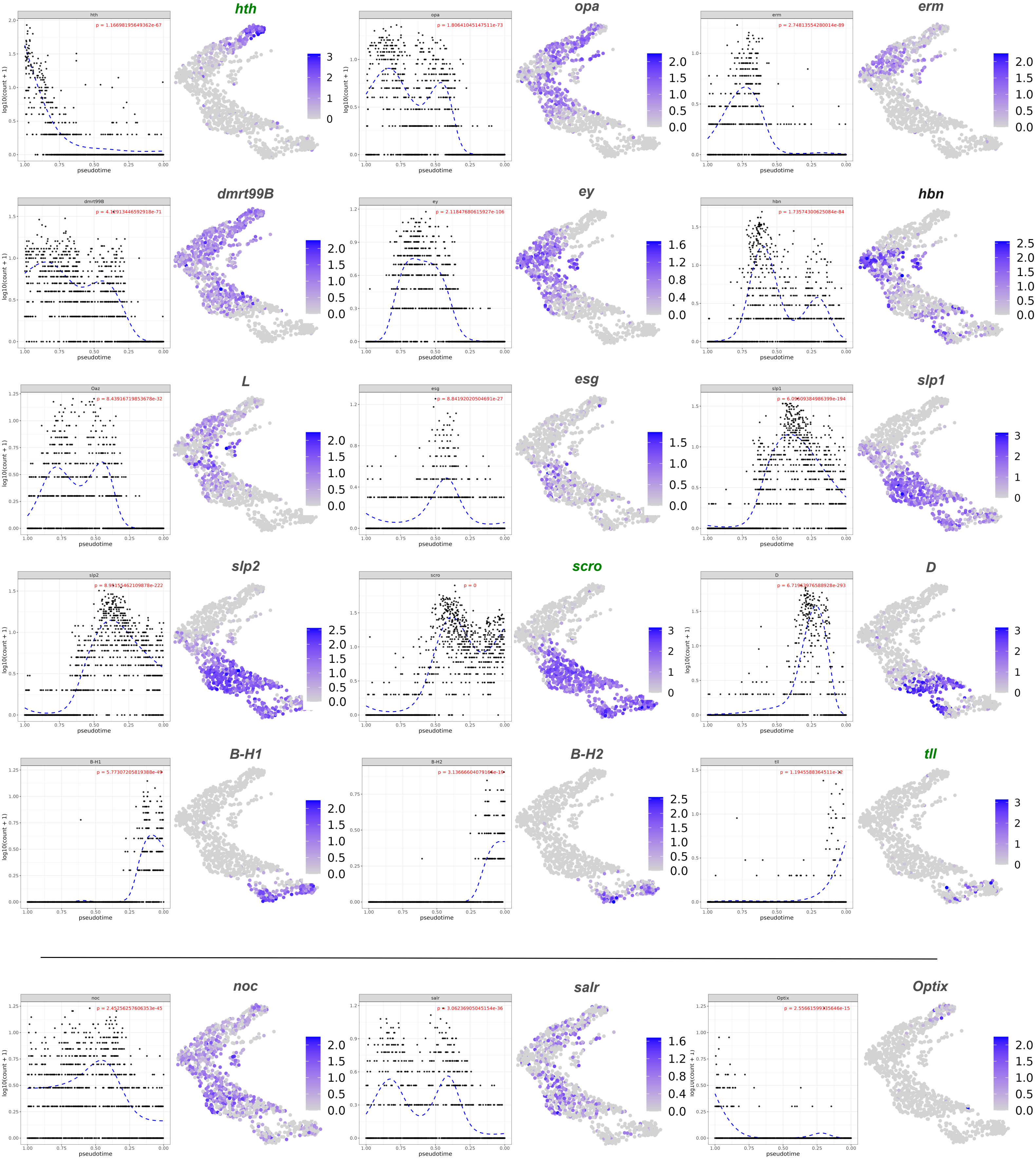

### Supplementary Figure 9

#### D.virilis

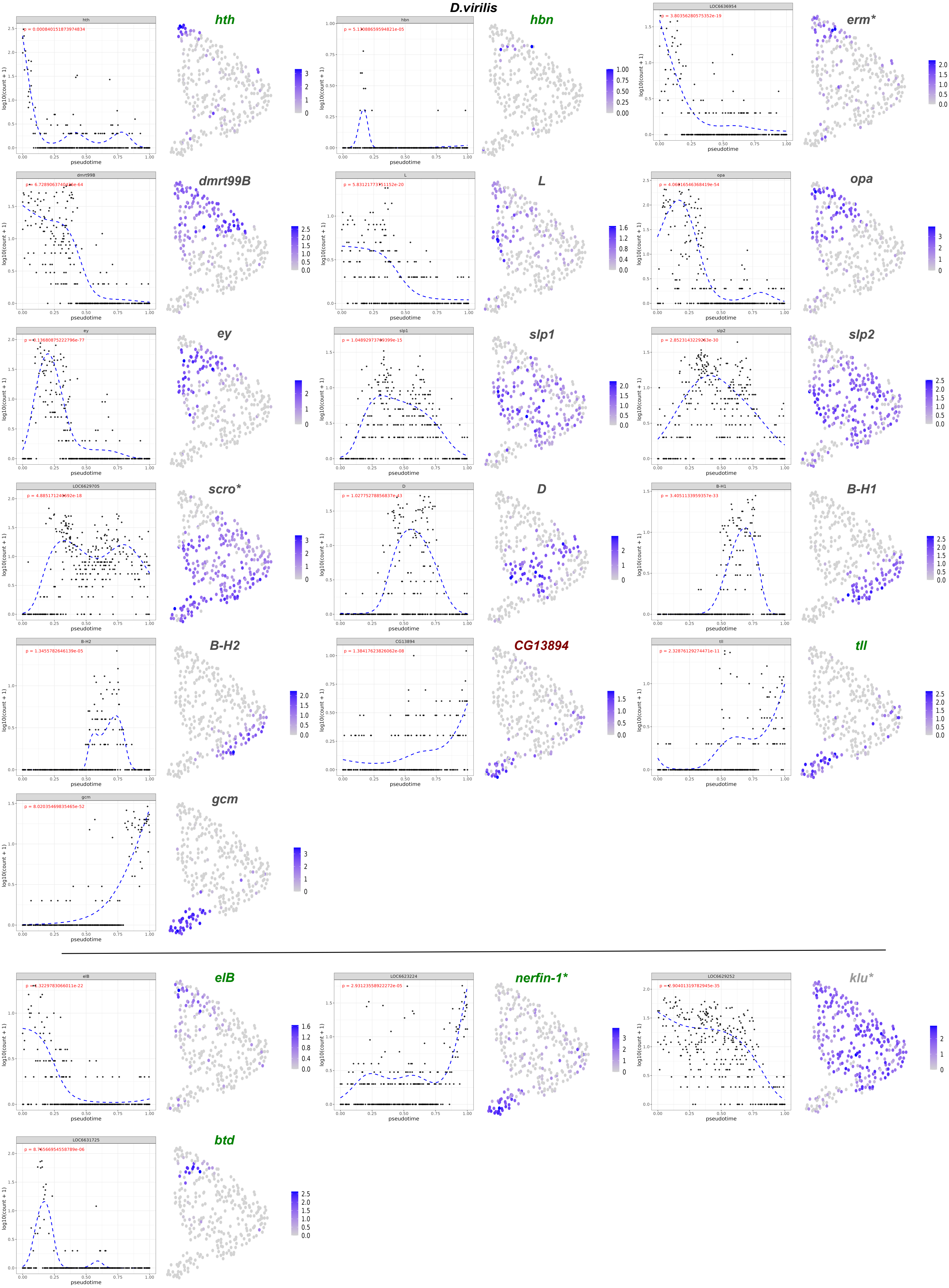

### Supplementary Figure 9

#### *M.domestica*

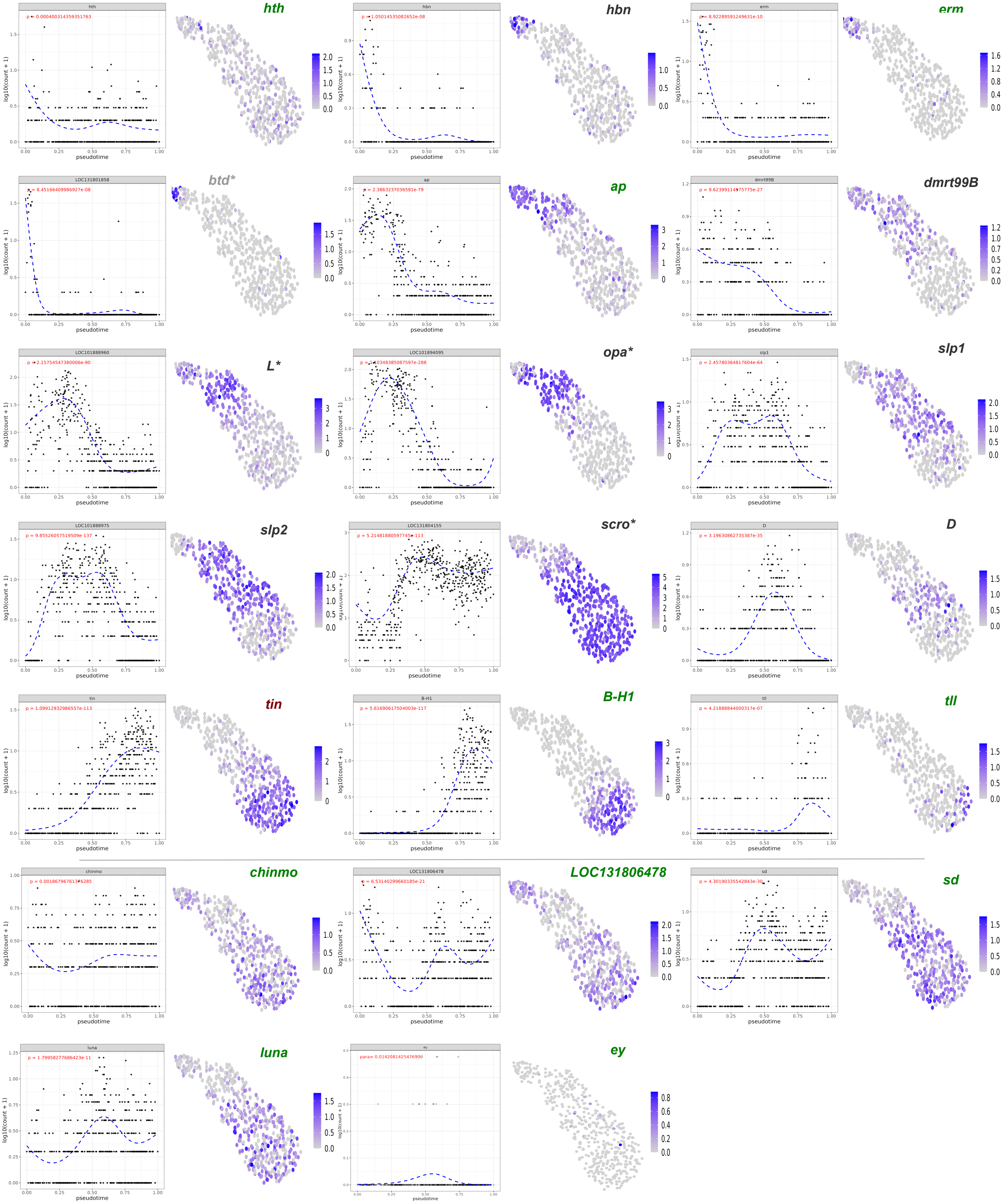

### Supplementary Figure 9

#### *A.aegypti*

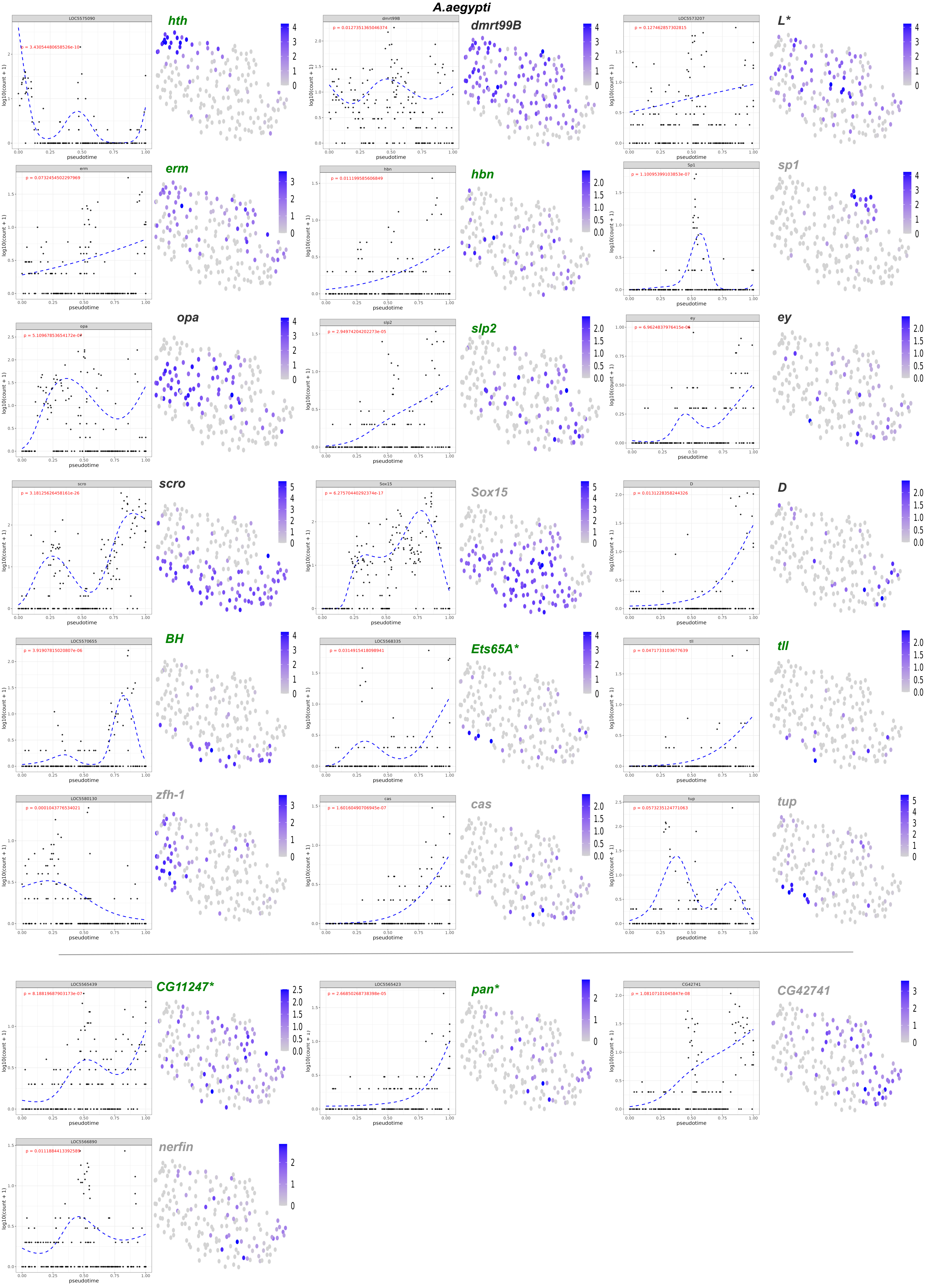

### Supplementary Figure 9

#### *B.mori*

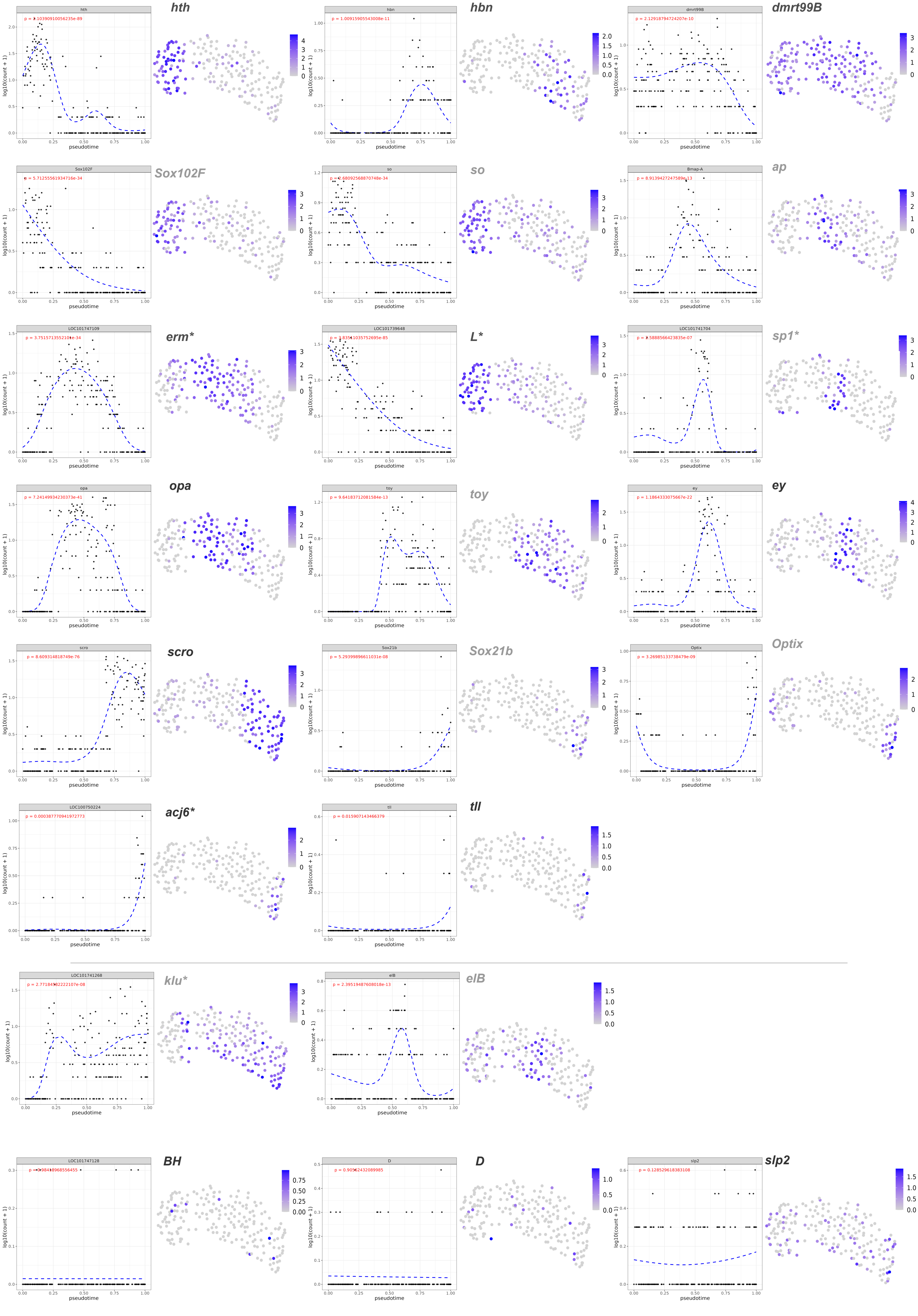

### Supplementary Figure 9

#### *T.castaneum*

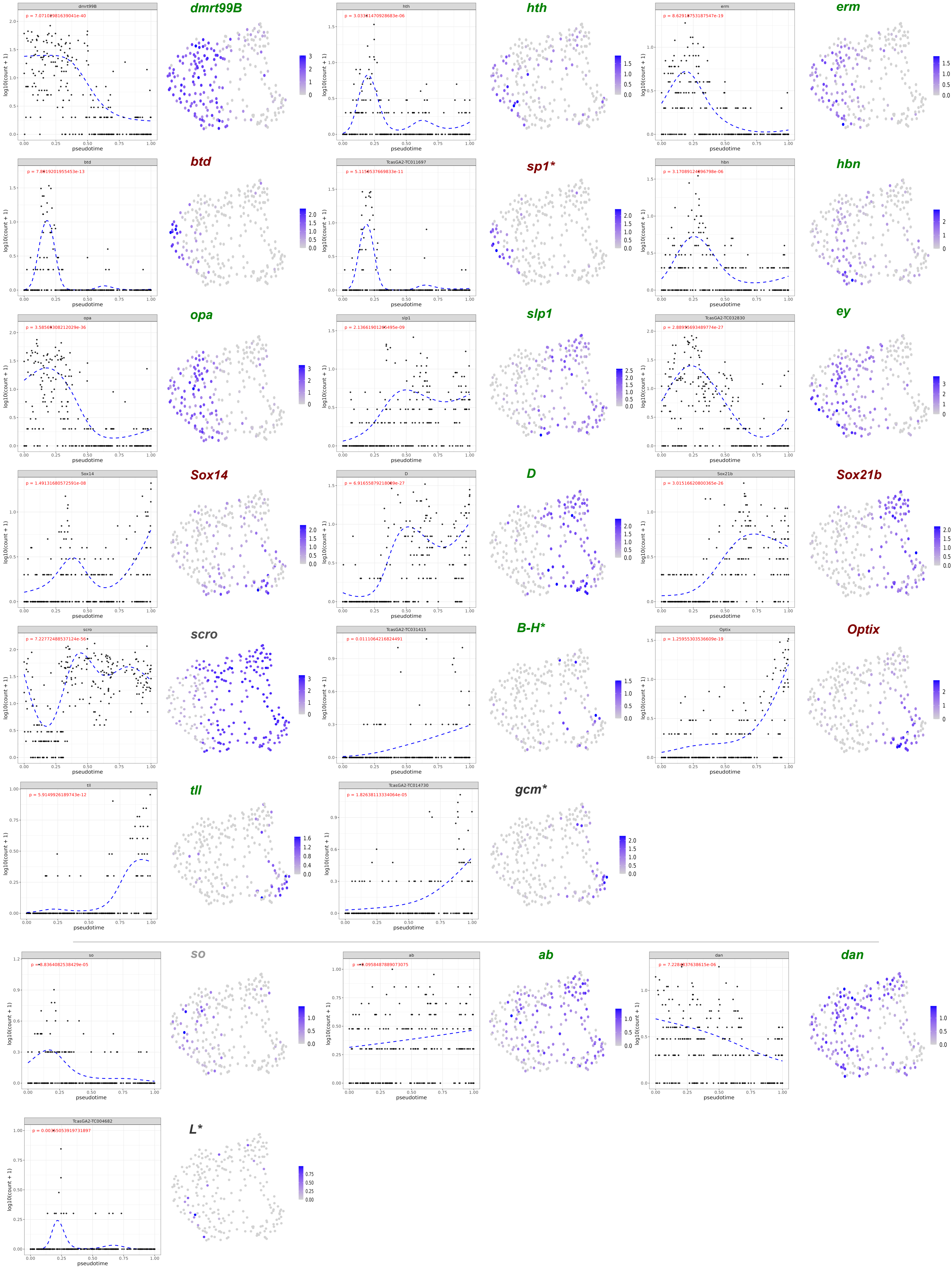

### Supplementary Figure 9

#### *N.vitripennis*

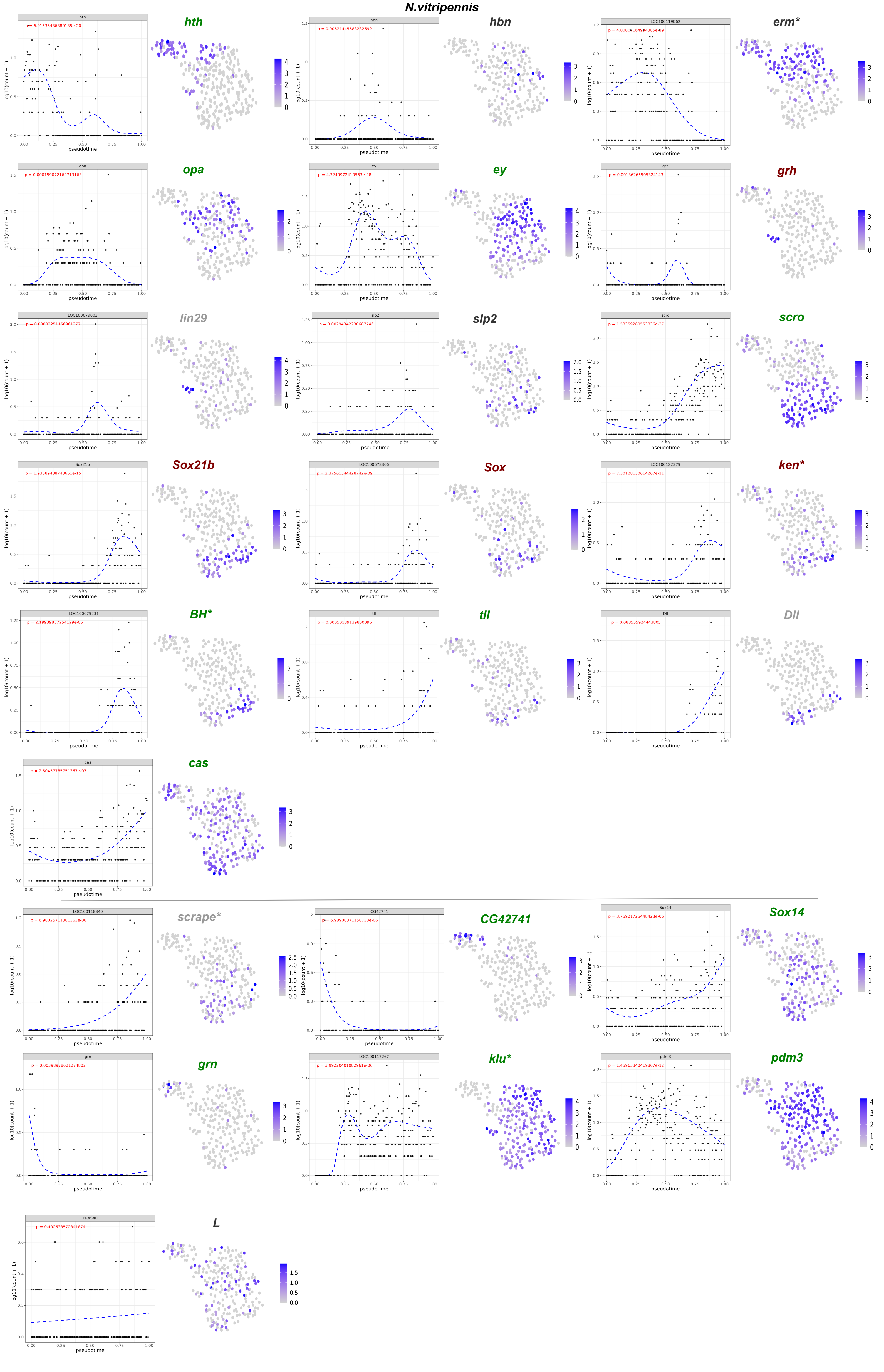

### Supplementary Figure 9

#### *G. bimaculatus*

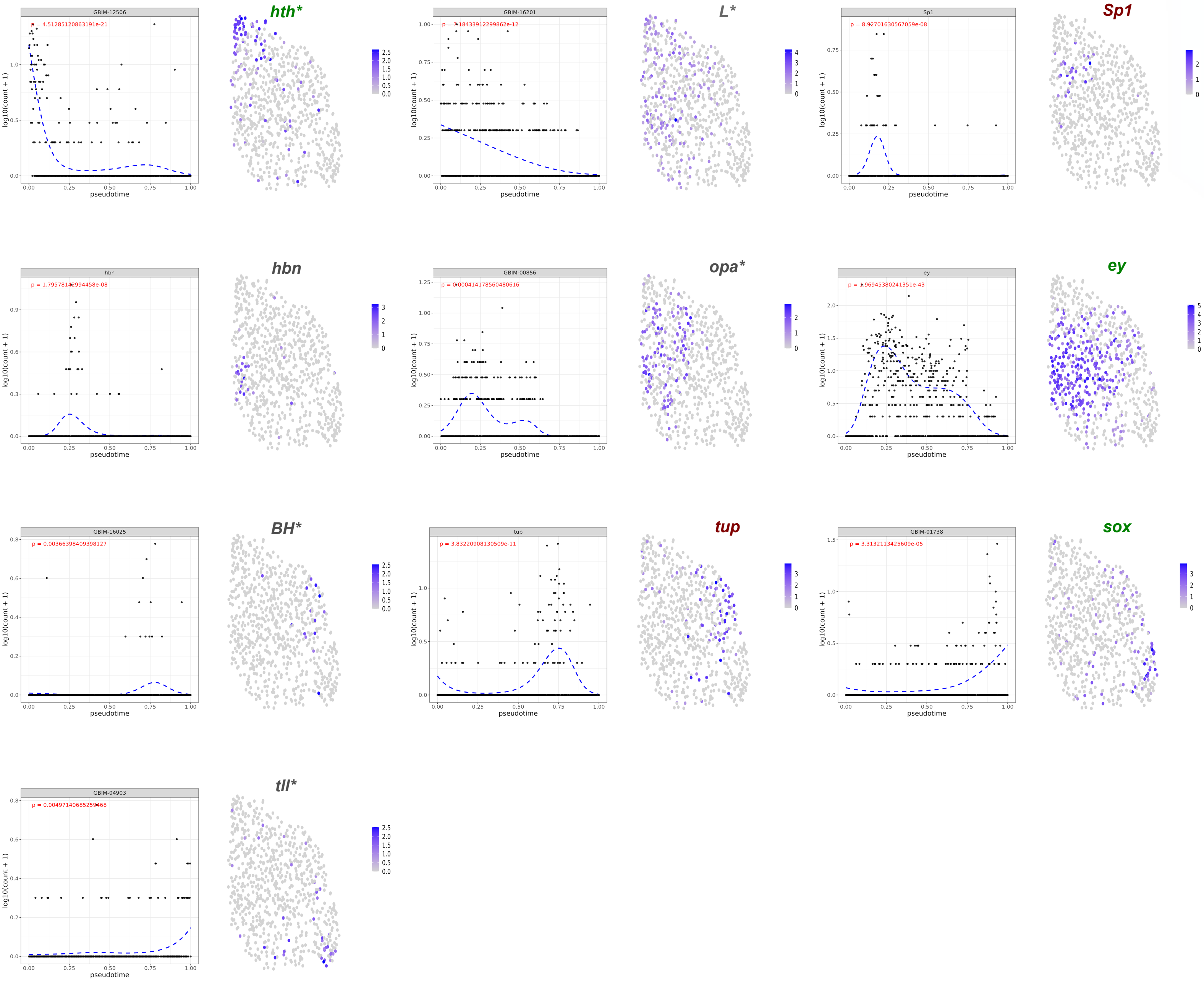

### Supplementary Figure 9

#### *C.dipterum*

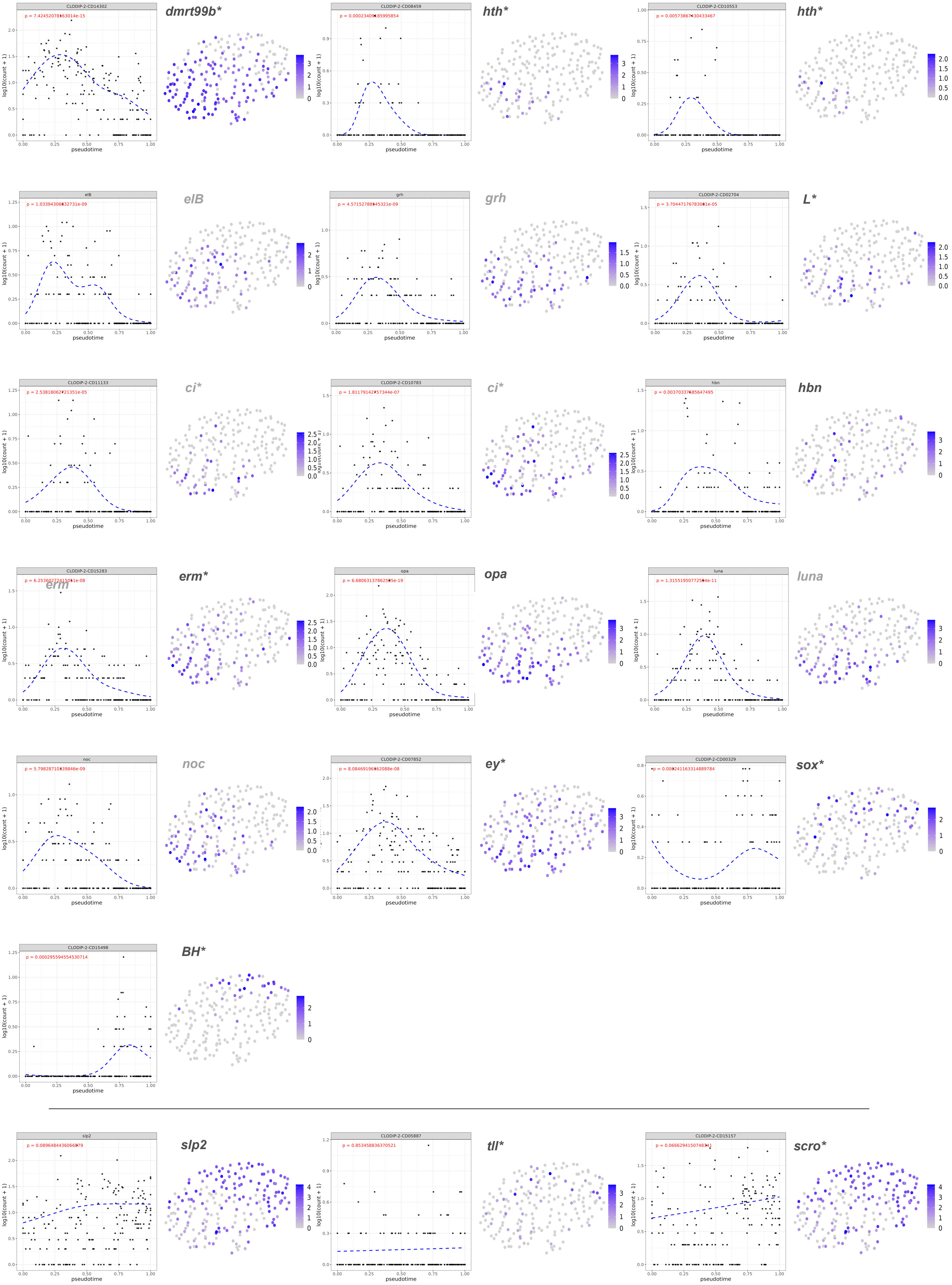
