## Supplementary material for "Evolutionary dynamics of temporal transcription factor series in the insect visual brain": Table S5

**Table S4: Insect strains, genomes, and reagents**

| REAGENT or RESOURCE | SOURCE | IDENTIFIER |
| --- | --- | --- |
| Insect strains | |  |
| *D. melanogaster* | Desplan lab | Canton-S |
| *D. virilis* | Courtier lab | 15010-1051.86 |
| *M. domestica* | Bopp lab | M3 strain |
| *A. aegypti* | Failloux lab | Paea strain |
| *B. mori* | Silkworm Genetic Resource Database | p50 strain |
| *T. castaneum* | Bucher lab | San Bernardino strain |
| *N. vitripennis* | Beukeboom lab | AsymCx wild-type strain |
| *G. bimaculatus* | Roth lab | White-eyed wild-type strain |
| *C. dipterum* | Almudi lab | Wild-type strain |
| Genomes |  |  |
| *D. virilis* | NCBI | GCF_003285735.1_DvirRS2 |
| *M. domestica* | NCBI | GCF_030504385.1_Musca_domestica.polishedcontigs.V.1.1 |
| *A. aegypti* | NCBI | GCF_002204515.2_AaegL5.0 |
| *B. mori* | NCBI | GCF_014905235.1_Bmori_2016v1.0 |
| *T. castaneum* | NCBI | GCA_000002335.3_Tcas5.2 |
| *N. vitripennis* | NCBI | GCF_009193385.2_Nvit_psr_1.1 |
| *G. bimaculatus* | NCBI | GCA_017312745.1_Gbim_1.0 |
| *C. dipterum* | NCBI | GCA_902829235.1_CLODIP2 |
| Antibodies |  |  |
| Anti-acetylated tubulin | Sigma Aldrich | 06-896 |
| Reagents |  |  |
| Click-iT EdU Imaging Kit | Molecular Probes | C10086 |
| Collagenase, Type I, powder | Gibco | 17100017 |
| Dispase de type II, poudre | Gibco | 17105041 |
| TrypLE | Gibco | A12177-01 |
| Shneiders insect medium | Gibco | 21720024 |
| Grace’s insect medium | Gibco | 11667037 |
| Formaldehyde (37%), 10-15% Methanol as stabilizer | Sigma-Aldrich | 252549 |
| Formaldehyde (16%), Methanol free | ThermoFischer | 28908 |
| Triton-X-100 | Sigma-Aldrich | X100 |
| Vectashield | Eurobio-scientific | H-1000-10 |
| D-PBS | Gibco | 14190144 |
| Bovine Serum Albumin | Sigma-Aldrich | A9418 |
| Saline Sodium Citrate | Invitrogen | AM9770 |
| Salmon sperm DNA | Sigma-Aldrich | D7656 |
| Probe hybridization buffer | Molecular Instruments |  |
| Probe wash buffer | Molecular Instruments |  |
| Amplification buffer | Molecular Instruments |  |
| Hoechst | Miltenyi Biotec | 130-111-569 |
| 20 μm strainer | pluriSelect | 43-10020-70 |
| Eppendorf DNA LoBind tubes | Sigma-Aldrich | EP0030108051-250EA |
| Next GEM Single Cell 3ʹ Reagent Kits v3.1 | 10x Genomics | 1000269 |
| Next GEM Chip G Single Cell Kit | 10x Genomics | 1000127 |
| Dual Index Kit TT Set A | 10x Genomics | 1000215 |
